# *In-silico* Analysis of SARS-Cov2 Spike Proteins of Different Field Variants

**DOI:** 10.1101/2023.01.22.525048

**Authors:** Muhammad Haseeb, Afreenish Amir, Aamer Ikram

## Abstract

**Background:** Coronaviruses belong to the group of RNA family of viruses which trigger diseases in birds, humans, and mammals, which can cause respiratory tract infections. The COVID-19 pandemic has badly affected every part of the world, and the situation in the world is getting worse with the emergence of novel variants. Our study aims to explore the genome of SARS-,CoV2 followed by *in silico* analysis of its proteins.

**Methods:** Different nucleotide and protein variants of SARS-Cov2 were retrieved from NCBI. Contigs & consensus sequences were developed to identify variations in these variants by using SnapGene. Data of variants that significantly differ from each other was run through Predict Protein software to understand changes produced in protein structure The SOPMA web server was used to predict the secondary structure of proteins. Tertiary structure details of selected proteins were analyzed using the online web server SWISS-MODEL.

**Findings:** Sequencing results shows numerous single nucleotide polymorphisms in surface glycoprotein, nucleocapsid, ORF1a, and ORF1ab polyprotein. While envelope, membrane, ORF3a, ORF6, ORF7a, ORF8, and ORF10 genes have no or few SNPs. Contigs were mto identifyn of variations in Alpha & Delta Variant of SARs-CoV-2 with reference strain (Wuhan). The secondary structures of SARs-CoV-2 proteins were predicted by using sopma software & were further compared with reference strain of SARS-CoV-2 (Wuhan) proteins. The tertiary structure details of only spike proteins were analyzed through the SWISS-MODEL and Ramachandran plot. By Swiss-model, a comparison of the tertiary structure model of SARS-COV-2 spike protein of Alpha & Delta Variant was made with reference strain (Wuhan). Alpha & Delta Variant of SARs-CoV-2 isolates submitted in GISAID from Pakistan with changes in structural and nonstructural proteins were compared with reference strain & 3D structure mapping of spike glycoprotein and mutations in amino acid were seen.

**Conclusion:** The surprising increased rate of SARS-CoV-2 transmission has forced numerous countries to impose a total lockdown due to an unusual occurrence. In this research, we employed *in silico* computational tools to analyze SARS-CoV-2 genomes worldwide to detect vital variations in structural proteins and dynamic changes in all SARS-CoV-2 proteins, mainly spike proteins, produced due to many mutations. Our analysis revealed substantial differences in functional, immunological, physicochemical, & structural variations in SARS-CoV-2 isolates. However real impact of these SNPs can only be determined further by experiments. Our results can aid *in vivo* and *in vitro* experiments in the future.

## Introduction

On December 31, 2019, COVID-19 was initially discovered in Wuhan, China. The condition became severe when many infected cases were reported in the “Huanan Seafood Market” (Shaikh et al., 2020). HKU1, HCoV229E, HCoVOC43, HCoVNL63, and HCoV229E are coronaviruses generally responsible for only minor common cold & respiratory infections in newborn infants and the elderly (Wormser & Aitken, 2010). Based on genetic material properties, the coronavirinae family contains four genes: Alpha, Beta, Gamma, and Delta coronavirus. Coronaviruses are RNA viruses that can trigger an infection in mammals, humans, & birds. They cause respiratory infections like the common cold, SARS, & MERS (Pal et al., 2020). SARS-Cov2 is a positive-polarity single-stranded RNA virus It started spreading immensely fast and became a worldwide pandemic within a few months. Its transmission between individuals and populations is relatively easy because of its transmission pattern, which is in the form of direct body contact and through respiratory droplets from an infected individual (Lauer et al., 2020b). The virus has a 2 to 14-day incubation period and causes severe respiratory problems. Its symptoms are high-grade fever, nonproductive cough, pharyngitis, muscle joint pains, runny nose, diarrhea, shortness of breath, and in certain circumstances, loss of sensations such as taste and smell are lost (Lauer et al., 2020a).

The four critical structural proteins found in the virion are N-Protein (Nucleocapsid), M-protein (Transmembrane), E-Protein (Envelope), and S-Protein (Spike). However, structural proteins’ direct assembly is not required to form the whole virion infection in some coronaviruses; other proteins with overlapping compensatory roles may be expressed (Arias-Reyes et al., 2020; Perlman & Netland, 2009; Vlasova et al., 2007). The SARS-CoV-2 genome comprises two enormous linear ORFs, ORF1a and ORF1b ORF1a is transcribed into polyproteins 1a and 1b, which are produced via one ribosomal frameshift. For replication and transcription, there are 16 non-structural proteins (NSPS) (Graham et al., 2008).

But worldwide spread of COVID-19 has upraised significant concerns about viral evolution and adaptation. How it spreads worldwide, encountering various host immune systems and countermeasures determined by mutations, deletions, and recombination.

SARS-Cov2 varies from earlier strains by having numerous hazardous residues in the Corona Virus receptor-binding region (especially Gln493), which delivers valuable communication with ACE2 human receptors (Zhang et al., 2020). Understanding surface receptors and their variations in the field is also crucial for developing a stable vaccine strain. Therefore, we will employ *in silico* screening of the whole genome for significantly spiked protein SARS-Cov2 of coronavirus variants from published data worldwide to understand viral variation patterns. As a result, SARS-CoV-2 positive samples were sequenced to investigate the genetic diversity during the fourth wave of the epidemic in Pakistan.

There is a pressing need to combat COVID-19, and we need quick and reliable approaches in this regard. SARS-Cov2 varies from earlier strains by having numerous residues in the coronavirus receptor-binding region, which enables its binding with ACE2 human receptors. The change in closeness perhaps explains why this virus is more transmissible than other viruses. Using *in silico* techniques, we will study the pattern of variation in the proteins, especially the spike proteins, of various SARS-Cov2 strains reported across the globe. This research will help lay the basis for future research.

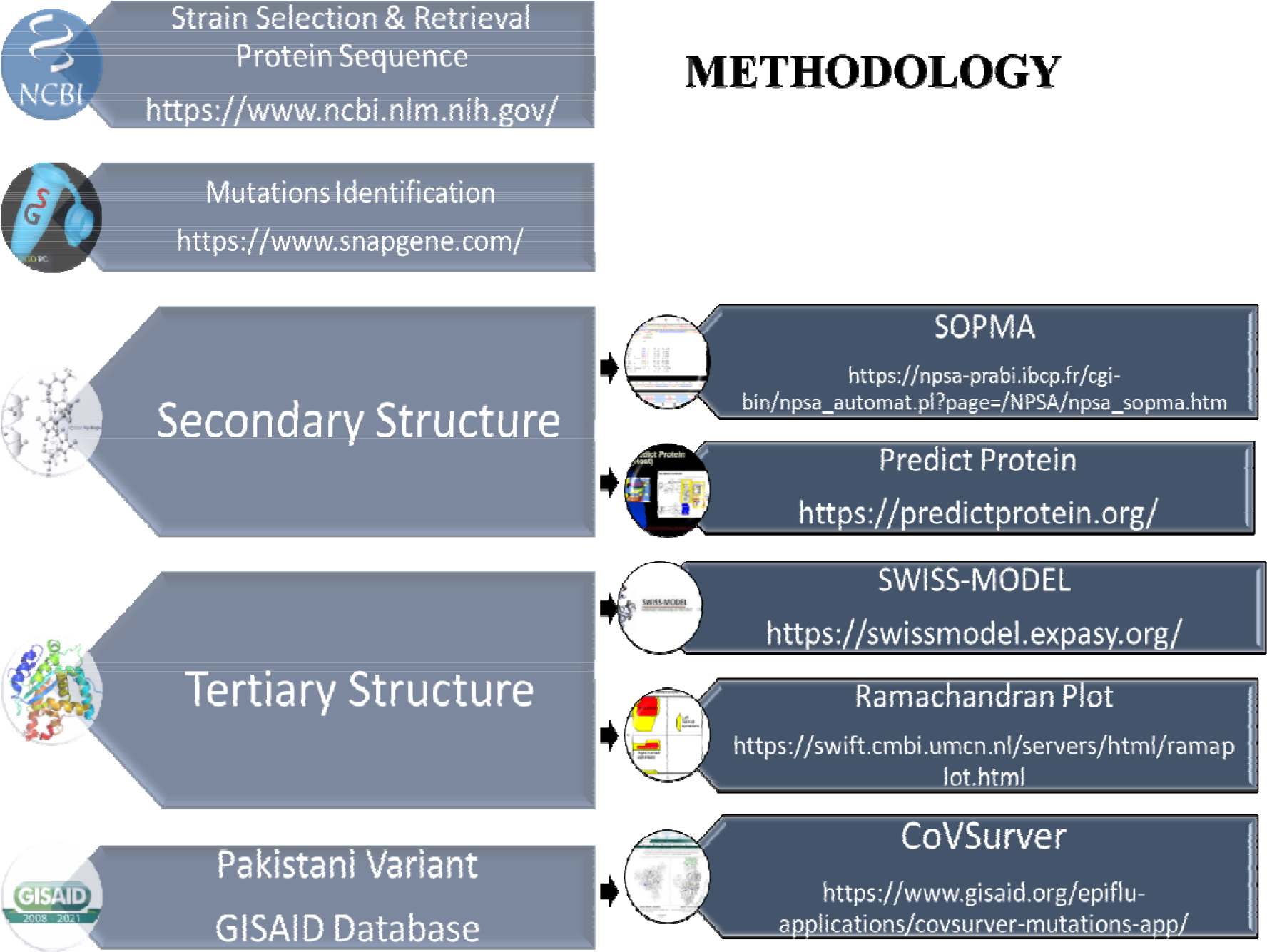

## Results

### Mutations Identified in the Sequenced SARS-CoV-2 Genomes

Sequencing results demonstrated numerous single nucleotide polymorphisms (SNPs) in Surface glycoprotein, Nucleocapsid & ORF1a polyprotein 1ab (ORF1ab), and ORF1ab polyprotein. These mutation hotspots may be of particular importance in adapting SARS-CoV-2s to the human host. While the Envelop, Membrane, ORF3a, ORF6, ORF7a, ORF8, and ORF10 genes have no or few SNPs, limited alterations in these proteins might mean that they have conserved activities that are required for viral transmission.

Contigs were made for the identification of variations in Alpha Variant (B.1.1.7) and Delta Variant (B.1.617.2) of SARs-CoV-2 with reference strain (Wuhan) by using SnapGene (https://www.snapgene.com/) version 5.3 software, and consensus sequences were developed as shown in Supplementary file.

**Figure 12:**
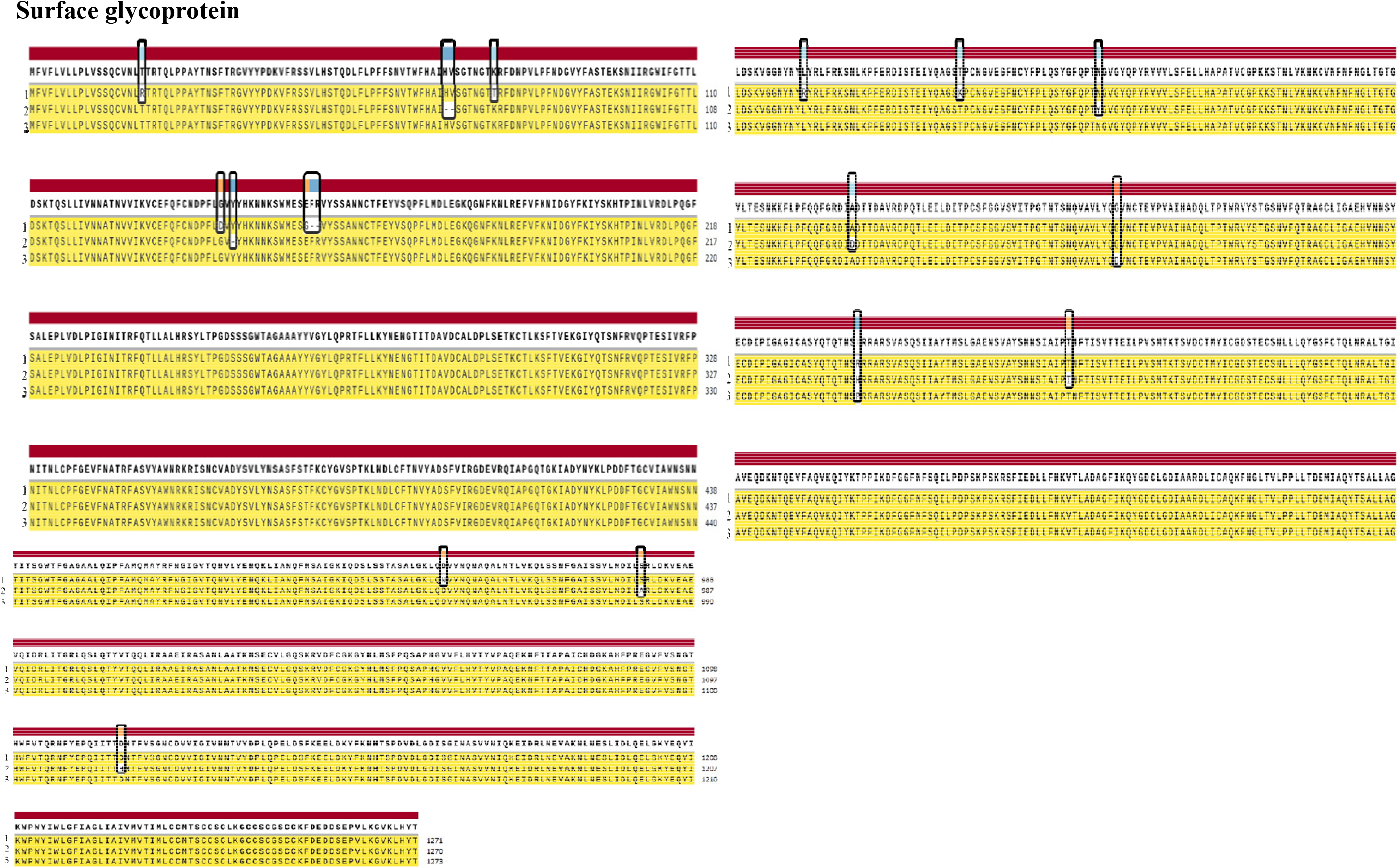
Schematic view of contigs of the Surface glycoprotein of 1. Alpha Variant (B.1.1.7) (UDQ41838.1) and 2. Delta Variant (B.1.617.21) (UDU36746.1) of SARs-CoV-2 with 3. reference Strain (Wuhan) (YP_009724390.1)

### Protein Secondary Structure

The secondary structures of the envelope protein, membrane glycoprotein, nucleocapsid phosphoprotein, ORF10 protein, ORF1a polyprotein, ORF1ab polyprotein, ORF3a protein, ORF6 protein, ORF7a protein, ORF7b protein, ORF8 protein, and surface glycoprotein of SARS-COV-2 were predicted by using the sopma secondary structure prediction method https://npsa-prabi.ibcp.fr/cgi-bin/npsa_automat.pl?page=/NPSA/npsa_sopma.html By keeping default parameters, a number of conformational states: 4 (Helix, Sheet, Turn, Coil), similarity threshold: 8 window width: 17, and were further compared with the reference strain of SARS-CoV-2 (Wuhan) proteins, which all showed similarity & variance to each other as shown in *Table 1* (Combet et al., 2000)

**Table 1.**
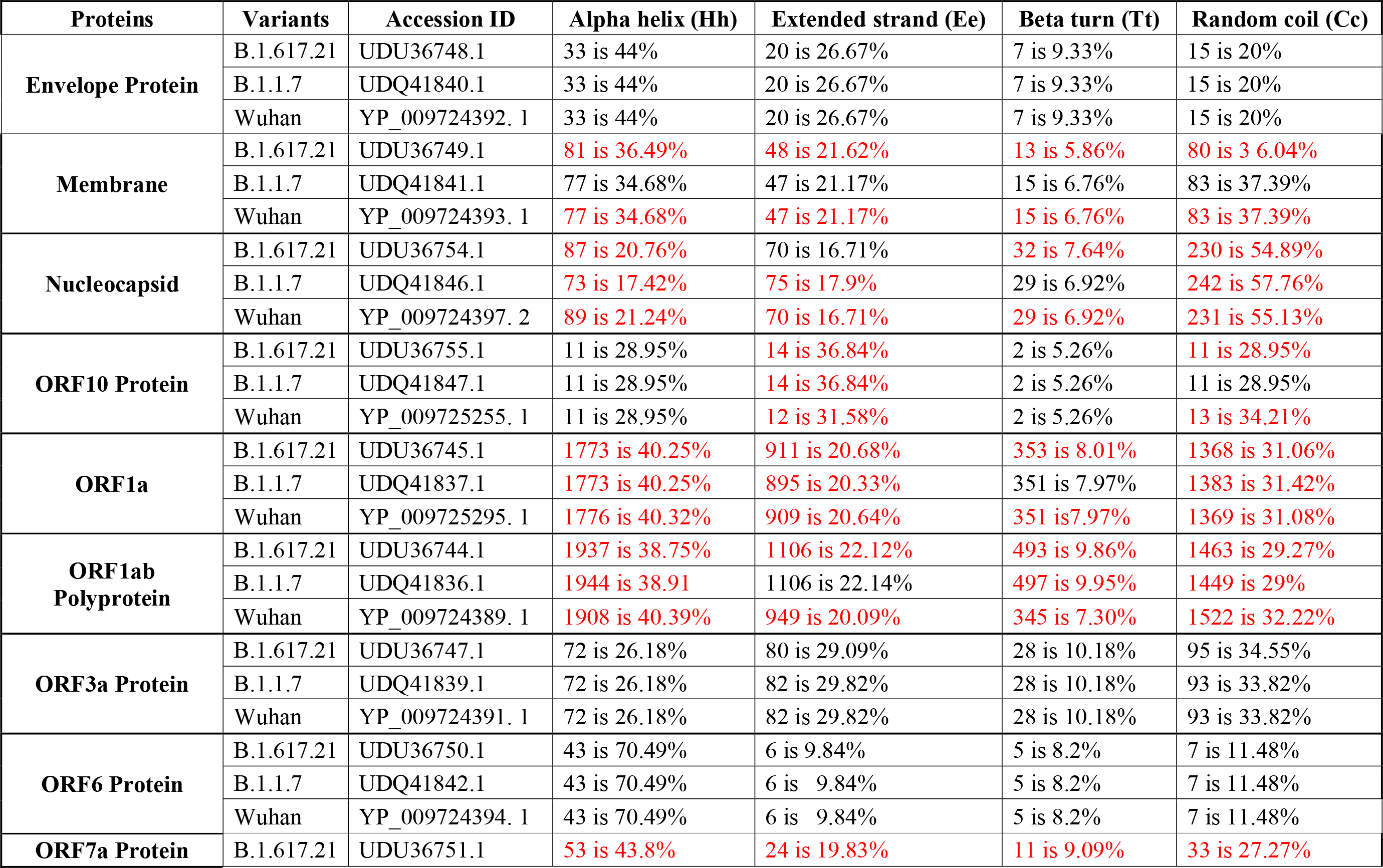
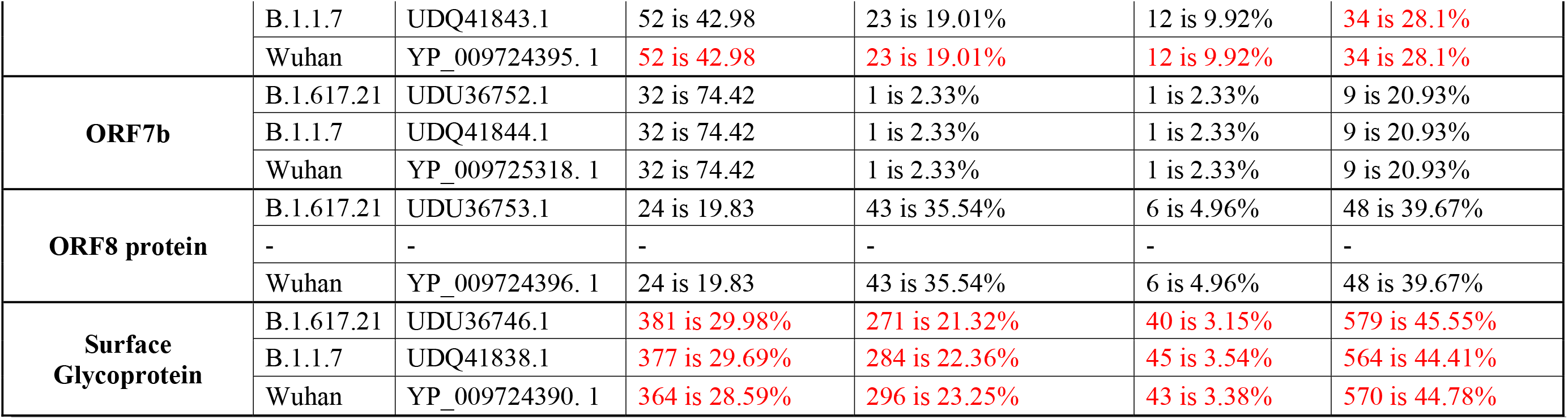
Comparison of the secondary structures of proteins of SARS-CoV strains B.1.617.21 and B.1.1.7 with a reference strain of SARS-CoV-2.

PredictProtein (https://predictprotein.org/) was used for the prediction of protein structural and functional features It was used for only those proteins with a drastic change in the secondary structure, as shown in supplementry material.

**Figure 17.**
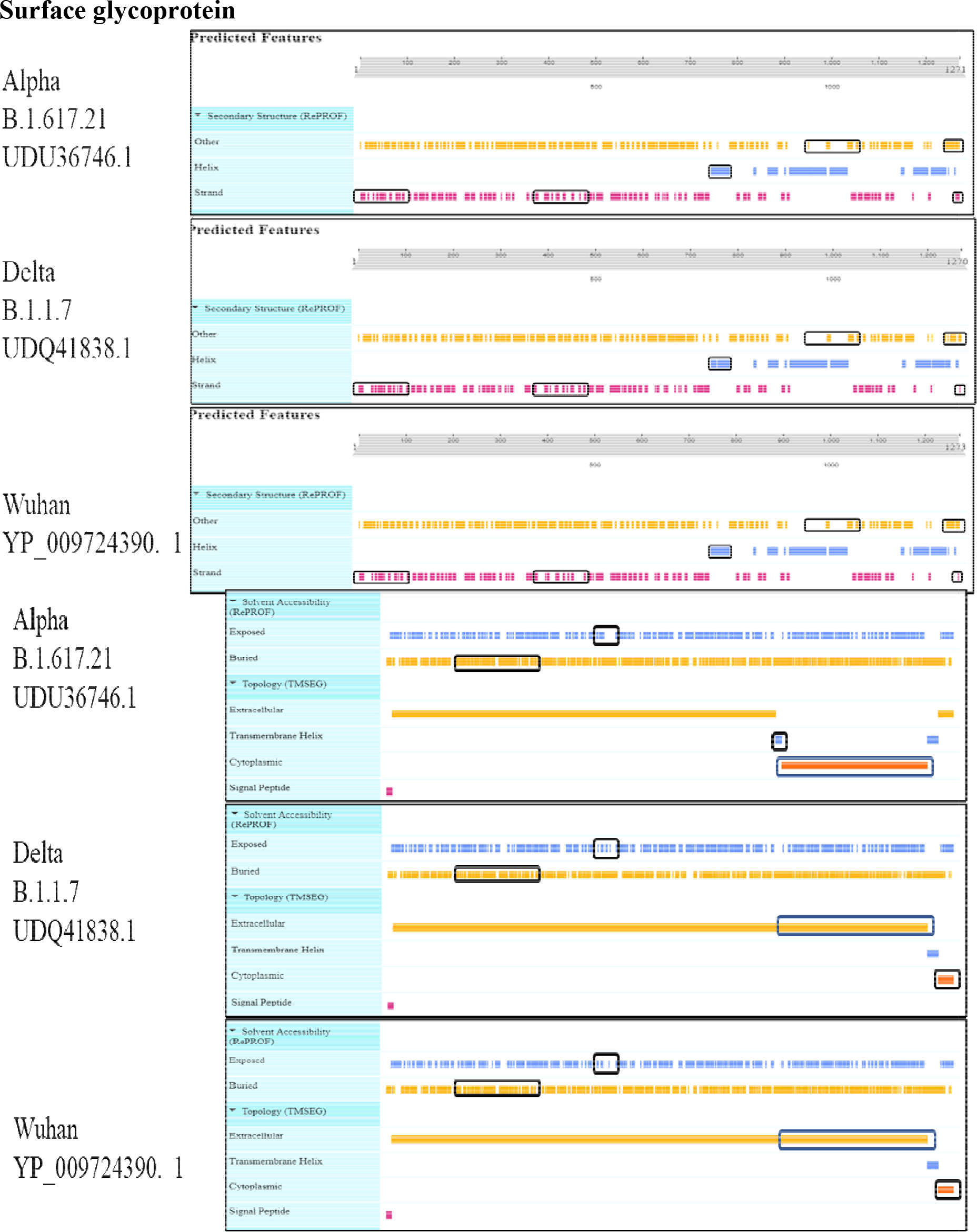
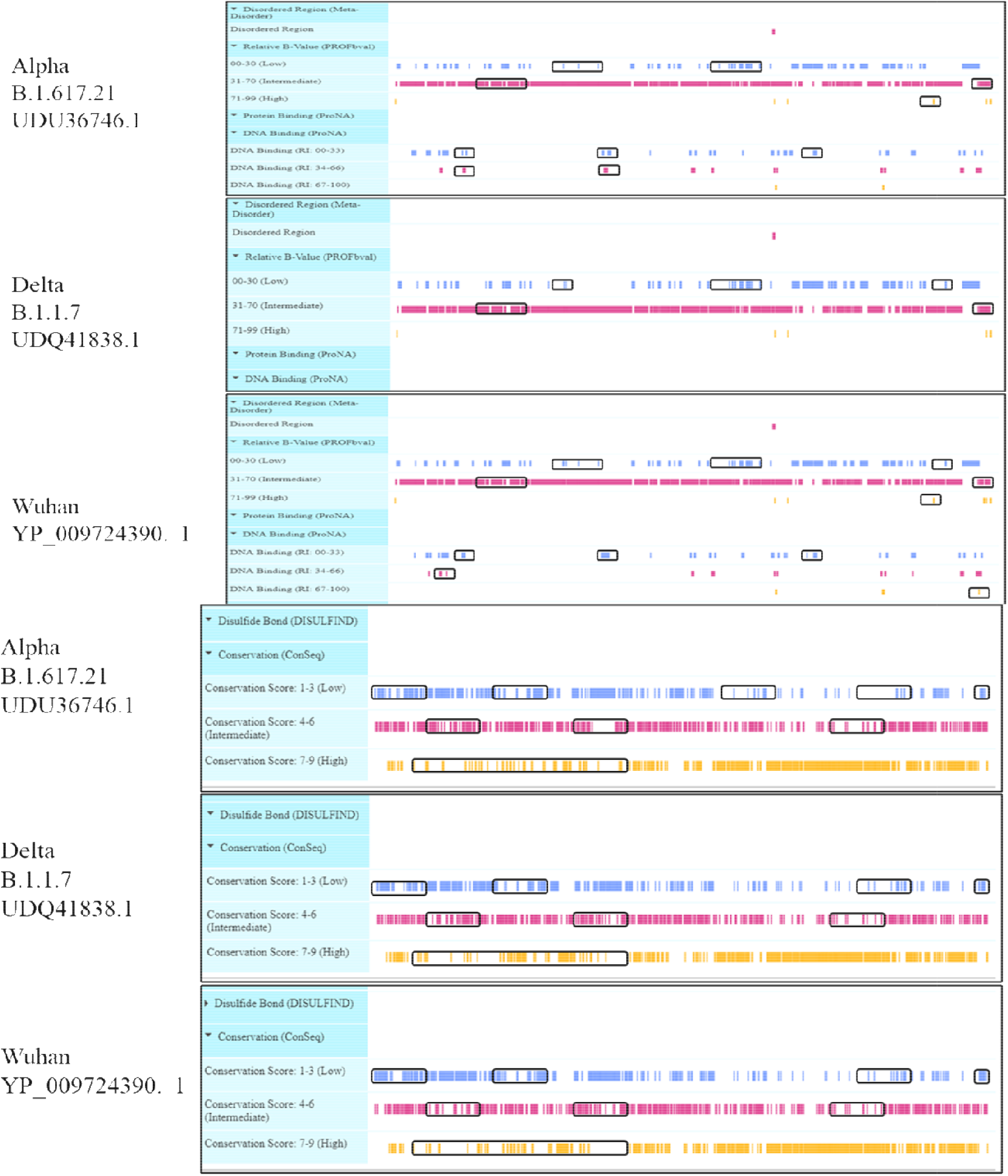
Viewer lays out predicted features of protein structural and functional features of Surface glycoprotein

### Tertiary Structure

Tertiary structures details of spike proteins was analyzed by using online software SWISS-MODEL (https://swissmodel.expasy.org/interactive) and Ramachandran plot (https://swift.cmbi.umcn.nl/servers/html/ramaplot.html). By Swiss-model, comparison of tertiary structure model of SARS-COV-2□spike protein of Alpha & Delta Variant was done with reference strain (Wuhan) as shown in fig (Waterhouse et al., 2018).

**Figure 18.**
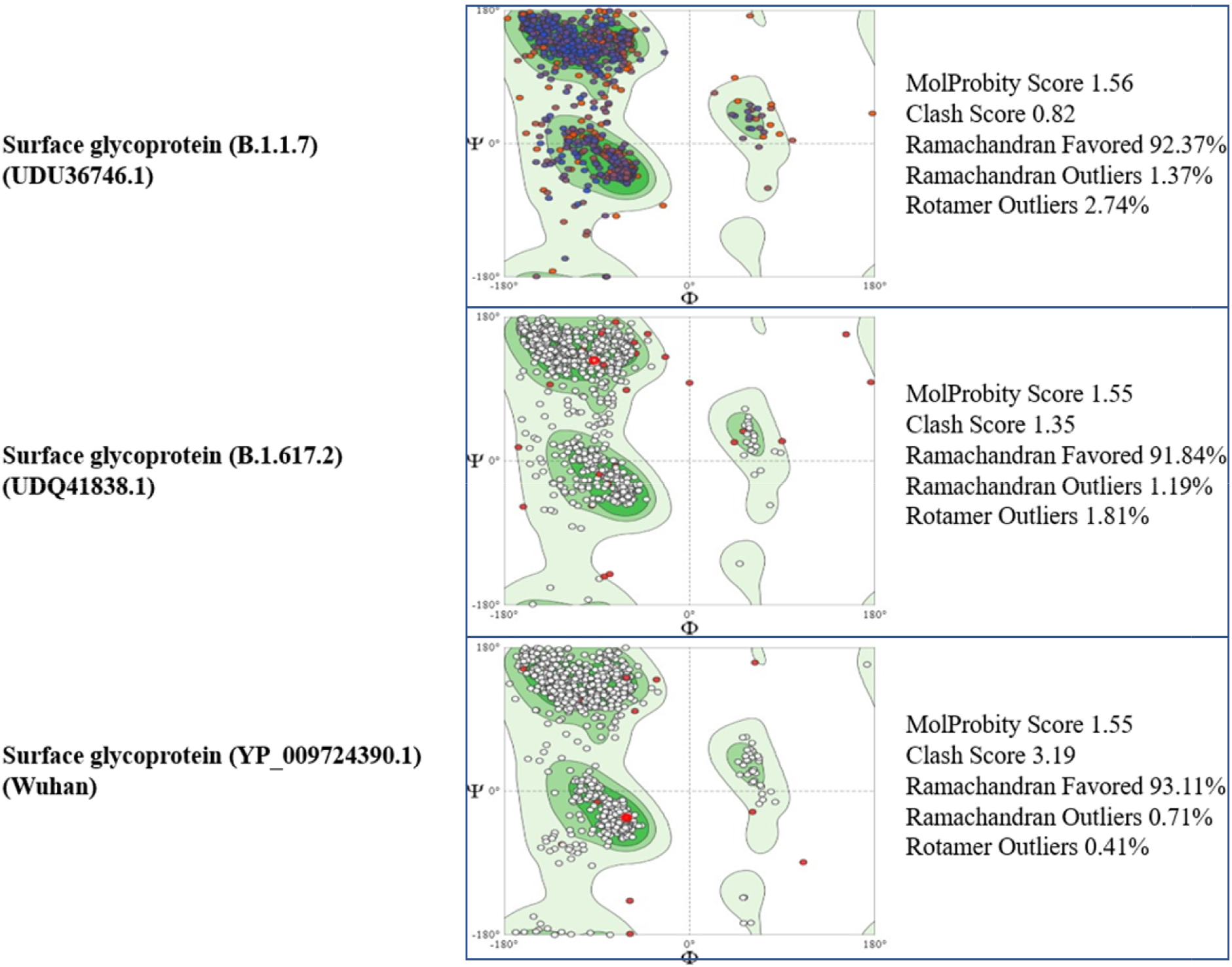
Comparison of Ramachandran Plots and MolProbity results of Surface glycoprotein

**Figure 19.**
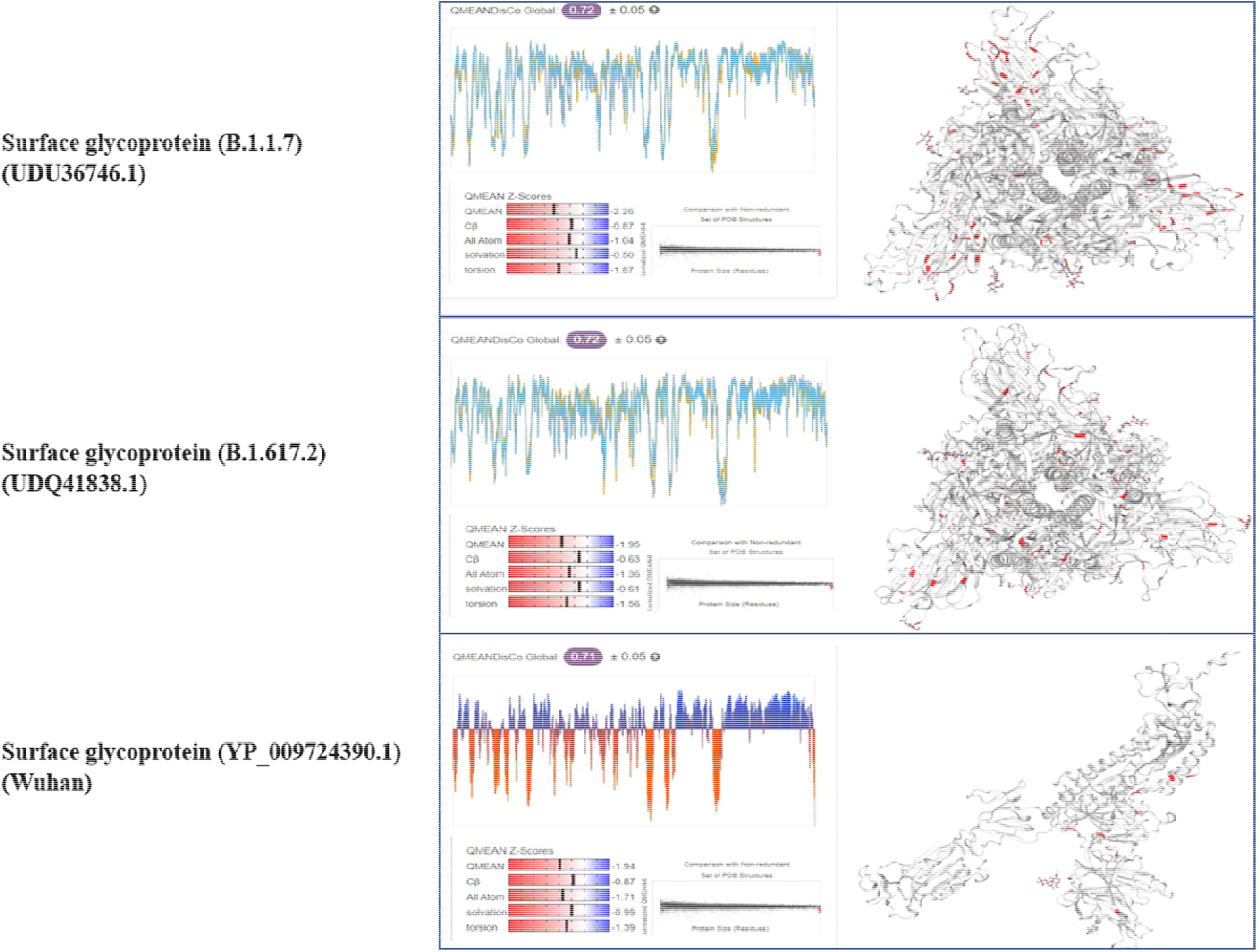
Comparison of Quality Estimate and Cryo-EM structure of PCoV_GX of spike glycoproteins

### Comparison of Pakistani Variants with reference Strain

Alpha & Delta Variant of SARs-CoV-2 isolates submitted in GISAID (https://www.epicov.org/epi3/frontend#43f233) from Pakistan with changes in amino acids in structural and nonstructural protein compared with Reference strain (hCoV-19/Wuhan/WIV04/2019) as shown in Table 2 & 3D structure mapping of spike glycoprotein and mutations in amino acid were seen as shown in supplementary material.

**Figure 20.**
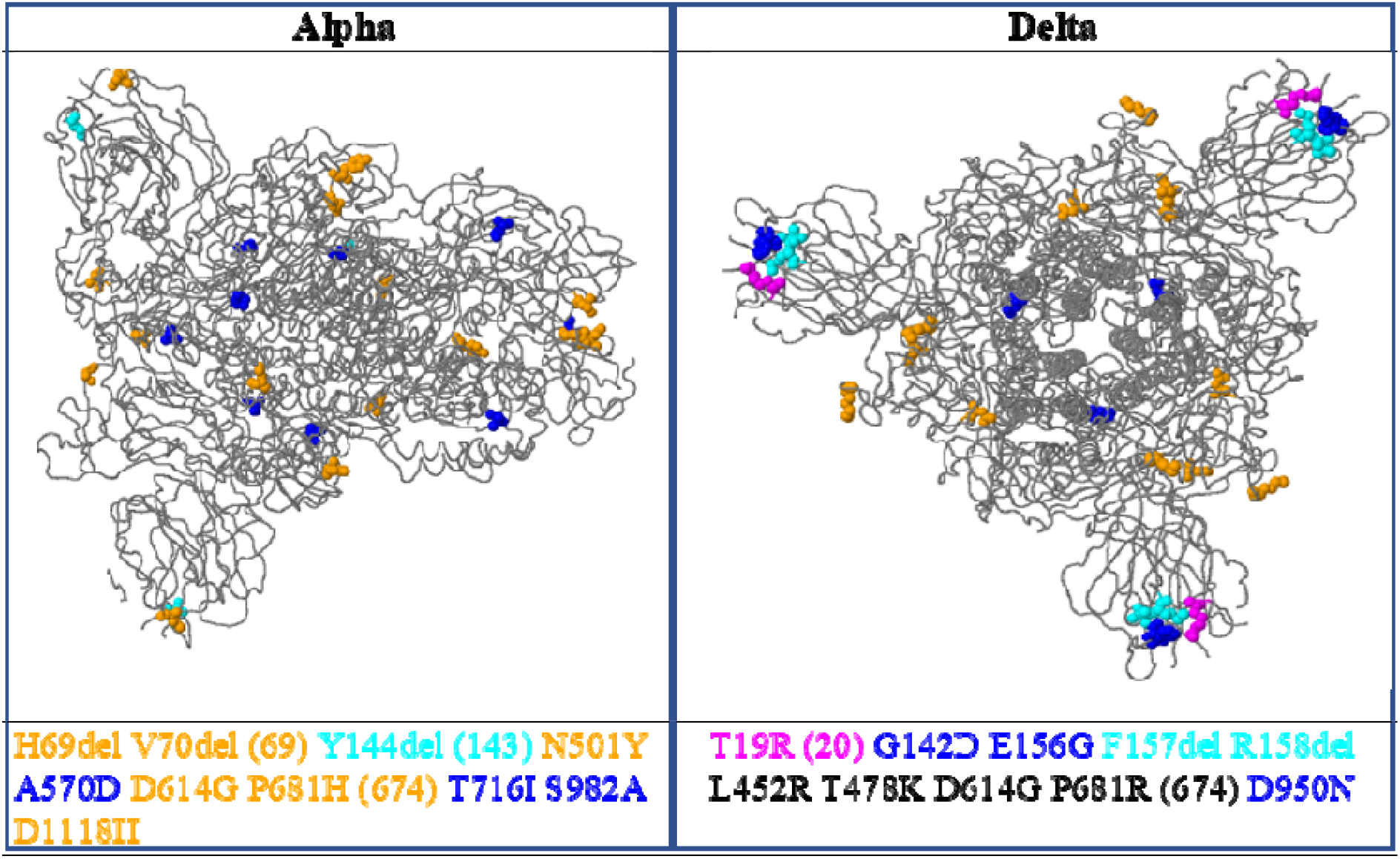
3D structural visualization of spike glycoprotein with amino acid changes shown as colored balls

## Discussion

In this research, we studied two complete sequences of Alpha and Delta Variant of SARS-CoV-2 with Wuhan variant as reference sequence. Several mutations have been noticed in the COVID-19 proteins as new variants emerge. These variations have several adverse effects on structure and function of COVID-19 proteins, making it challenging to administer the COVID-19 complex. Such mutations are discussed here.

Coronaviruses have the biggest genome size ranging from 26.4 to 31.7 kb of all RNA viruses (Mihindukulasuriya et al., 2008; Woo et al., 2009). It’s enormous gene size allows for greater flexibility in integrating and modifying genes (Mihindukulasuriya et al., 2008; Woo et al., 2009, 2010). In RNA viruses, mutation frequency is relatively high, which increases virulence and leads to the development of new species (Duffy, 2018). The greater rate of mutation within the genomes of viruses in various geographical areas is also one of reason that COVID-19 is liable for the changes in death rate & disease symptoms (Angeletti et al., 2020). In COVID-19, we found a few additional single amino acid changes in the Alpha & Delta Variant when compared to reference strain (Wuhan) as shown in above figures.

Virus particles contain the RNA genetic material and the structural proteins required for host cell entry. Once within the cell, the infecting RNA encodes structural proteins that form viral particles, nonstructural proteins that regulate viral assembly, transcription, replication, and control of the host cell, and accessory proteins whose role is unknown. The largest genome, ORF1ab, has overlapping open reading frames that encode the polyproteins PP1ab and PP1a. The polyproteins are degraded into 16 nonstructural proteins known as NSP1-16. A −1 ribosomal frameshifting occurrence is required to produce the longer (PP1ab) or shorter (PP1a) protein. Based on similarities to other coronaviruses, the proteins include the papain-like proteinase protein (NSP3), 3C-like proteinase protein (NSP5), RNA-dependent RNA polymerase (NSP12, RdRp), helicase (NSP13, HEL), endoRNAse (NSP15), 2’-O-Ribose-Methyltransferase (NSP16), and other non-SARS-CoV-2 non-structural proteins. They have the functions of viral transcription, replication, proteolytic processing, host immune response suppression, and host gene expression suppression.

Previous research has revealed that mutations in non-structural protein 2 & 3 play a crucial role in infectious capacity & are mainly accountable for SARS-CoV-2 differentiation process (Angeletti et al., 2020) and Coronavirus nucleocapsid protein is required for RNA replication, genome packing & transcription. (Masters, 2019). The envelope protein is involved in the viral genome assembly & the development of ion channels (IC), which are critical for virus-host connection and are primarily related to pathogenesis (Arias-Reyes et al., 2020; Ruch & Machamer, 2012).

Spike protein is about 180 to 200 kDa and has1273 amino acids (Hoffmann et al., 2020). To avoid the host’s immunological reaction, many polysaccharide molecules cover the spike protein’s surface (Watanabe et al., 2020). RBD of spike protein is the area that mainly interacts with ACE2, which leads to entry of virus into host cell (Angeletti et al., 2020; Tai et al., 2020). For several years, theoretical or experimental techniques have been used to predict protein stability (Sanavia et al., 2020). According to prior studies, a single point mutation in RBD disrupts the antigenic structure, affecting RBD binding to ACE2 (Babcock et al., 2004; Prabakaran et al., 2006). Furthermore, *in silico* investigations demonstrated that point mutations inside the RBD of spike glycoprotein had a stabilizing impact on spike protein and were discovered to enhance protein stability.

The potential changes in spike protein of COVID-19 discovered in alpha variants were H69del, V70del (69), Y144del (143), N501Y, A570D, D614G, P681H (674), T716I S982A, D1118H and potential changes in spike protein in delta variant were T19R (20), G142D, E156G, F157del, R158del, L452R, T478K, D614G, P681R (674) and D950N.

Some mutations may play a vital role in binding to human ACE2 receptors & many mutations may have enhanced the surface glycoprotein’s binding affinity. Additional beta strands and hydrogen bonds are shown in both the predictions and the 3D models. These extra mutations may produce structural conformational alterations and a greater binding affinity. Biologists and others can utilize these methods to gain a preliminary knowledge of SARS-CoV-2 mutations and their links to SARS-CoV & other related viruses. Initial observations can then be investigated further using various specialized tools and methods. Particularly computational techniques are promising. Machine learning, artificial intelligence, data integration and mining, visualization, computational and mathematical modelling for critical biochemical interactions, and disease control mechanisms can provide cost- and time-efficient solutions.

Researchers are collecting data related to Coronavirus in order to fully understand the spread of disease, pathogenesis & biology in order to eliminate it. (Douglas et al., 2020). The explosive growth of structural and genomic databases, combined with computational approaches, contributes to the discovery and manufacture of novel vaccination candidates. Modern breakthroughs in immunological bioinformatics have resulted in a variety of tools and web servers that can help cut the time and cost of manufacturing traditional vaccinations.

In our findings total forty substantial mutations were seen in alpha and ninety-two substantial mutations in delta variant in genomic sequences of Pakistani SARS-CoV-2 strains when compared with the Wuhan reference strain covering whole viral genome. Additional evaluation found that the majority of these changes were associated around a few viral genomic regions including spike, nucleocapsid protein, NS3, NSP2 and NSP6 as structural protein integrity is critical for immune response so we looked for mutations in viral proteins. Nucleocapsid protein is immunogenic phosphoprotein that helps in genome replication and regulation cell signaling pathway. The protein structure is disrupted due to mutation G204R in nucleocapsid and D614G in spike protein. Several studies have revealed that the mutation enhances viral infectivity. The D614G mutation expands the spike protein, which might result to protein instability and might also lead to increase viral infection.

We found 29 mutations in spike protein of delta variant and 14 mutations in alpha variant in genomic sequences of Pakistan SARS-CoV-2 strains when compared with reference strain. The SARS-CoV-2 spike protein is a prominent target for therapeutic & vaccine development due to its interaction in host cell receptor identification, attachment, and entrance (Liu et al., 2020; Ortega et al., 2020). In spike protein of both alpha and delta variants, we discovered a D614G (aspartic acid to glycine) mutation. In a recently study it was revealed that the D614G variant is more pathogenic, with infected individuals having a higher viral load; however, there was no correlation with disease severity (Korber et al., 2020). *In silico studies using pseudo viruses by Li et al. revealed that D614G mutation has a significant role in enhancing the infection (Li et al., 2020)*. Similar findings were reported, with the D614G mutant having the highest cell entrance among the spike variations (Hu et al., 2020). Furthermore, Hou et al. observed that D614G change improves SARS-CoV-2 infection rate & transmission primarily in humans & animal (Hou et al., 2020). The D614G mutation is becoming more common strain around the world.

In Pakistan, there was a 66% increase in SARS-CoV-2 cases & 64.8% increase in deaths in June 2020 when compared with February-May 2020. (72,460 confirmed cases and 1,543 deaths)(Umair et al., 2021). The significant rise in incidence cases & number of deaths may point towards widespread distribution of D614G mutation, that must be further examined (Becerra-Flores & Cardozo, 2020).

Additional molecular epidemiology investigations are required to track the DG614 strain’s circulation in Pakistan, that could also serve to understand impact of SARS-CoV-2 gene mutations on disease severity. Moreover, tracing variations in SARS-CoV-2 spike glycoprotein is critical because of its function in cell receptor interaction, entrance in host cell, and triggering antibody responses, as widespread distribution of the D614G variation throughout world influence on vaccination effectiveness and it has been a major concern, as most vaccines are designed on the D614G variation (Lurie et al., 2020). The same issue has been conveyed through results of Weissman et al., that shows D614G mutation is neutralized at a higher level by serum from vaccinated mice, non-human primates & humans (Weissman et al., 2021). Furthermore, regular monitoring of SARS-CoV-2 spike protein gene mutation is essential for detecting escape variants & in future for vaccine development.

This emphasizes the fact that SARS-CoV-2 strains circulating in Pakistan have mutated and have genetic variation from their origins. So, we suggest whole-genome sequencing of strains found throughout the country for better understanding the viral evolution & identifying strains with distinctive mutational changes.

## Conclusion

The surprising increased rate of SARS-CoV-2 transmission has forced numerous countries to impose a total lockdown due to an unusual occurrence. There is an immediate need to tackle COVID-19, so we want rapid and practical measures. *In silico* techniques are based upon analyzing biological data & using refined predictions and calculations to create a scientific database. In this research, we employed *in silico* computational tools to analyze SARS-CoV-2 genomes from across the world to detect vital variations in structural proteins as well as dynamic changes in all SARS-CoV-2 proteins, mainly spike proteins, produced due to large number of mutations.

Our analysis revealed substantial differences in functional, immunological, physicochemical, & structural variations in SARS-CoV-2 isolates. However real impact of these SNPs can only be determined further by experiments. Our results can aid *in vivo* and *in vitro* experiments in the future.

