## Supplemental Files for "*In-silico* Analysis of SARS-Cov2 Spike Proteins of Different Field Variants"

Muhammad Haseeb^1^, Sadia Sattar^1^, Afreenish Amir^2^, ^3^ r^4^

1. Department of Biosciences, Comsats University, Islamabad.

2. Department of Microbiology National Institute of Health, Islamabad,

3.

**Corresponding author**

**Muhammad Haseeb Tariq**

**National Institute of Health, Islamabad, Pakistan**

****

**Supplementary Figures**

1. **Envelope Protein**

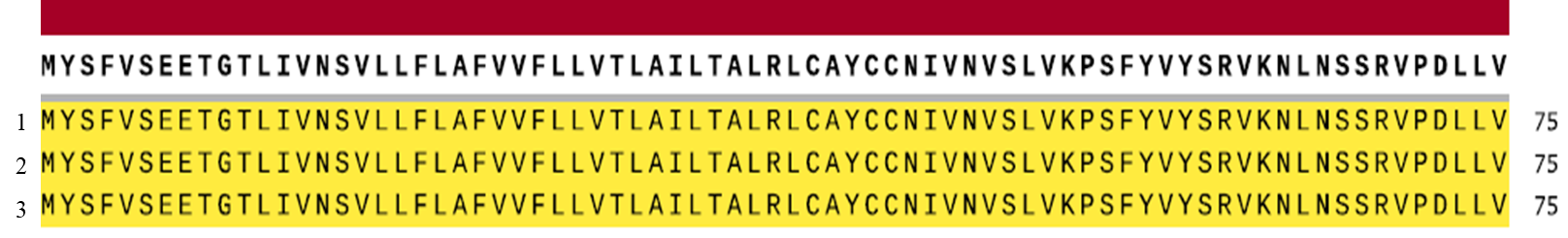

Figure 1 Schematic view of contigs of the envelope protein of 1. Alpha variant (B.1.1.7) (UDQ41840.1) and 2. Delta variant (B.1.617.21) (UDU36748.1) of SARs-CoV-2 with 3. reference strain (Wuhan) (YP_009724392.1)

2. **Membrane Protein**

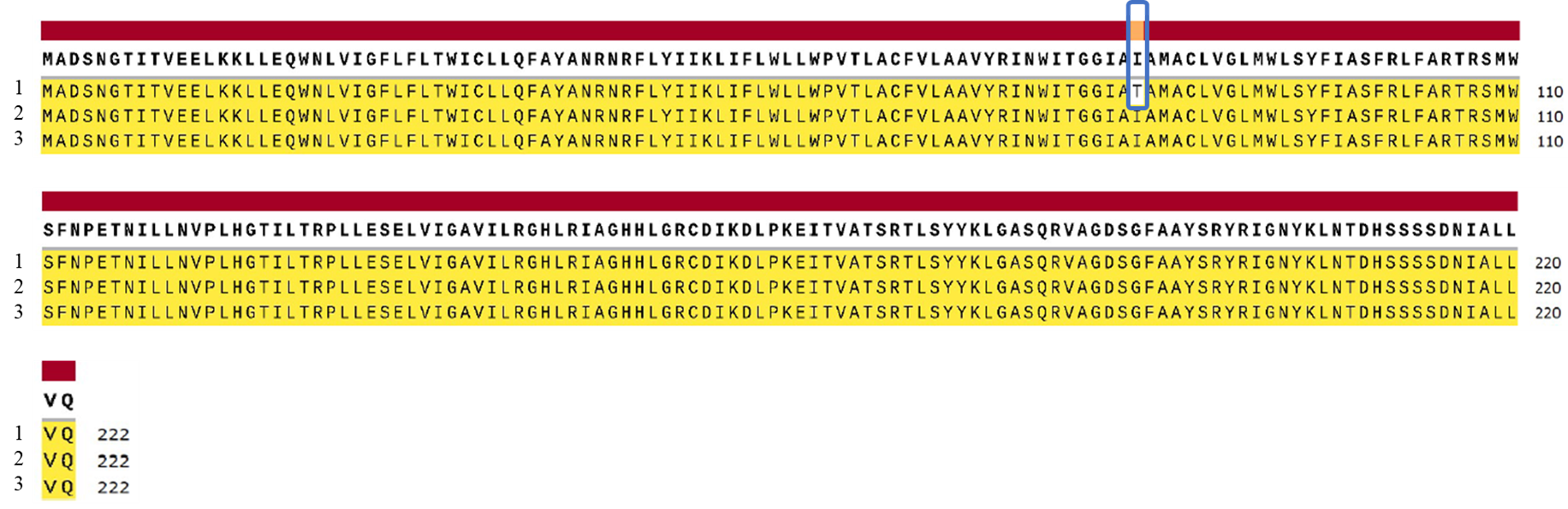

Figure 2: Schematic view of contigs of the membrane glycoprotein of 1. Alpha Variant (B.1.1.7) (UDQ41841.1) and 2. Delta Variant (B.1.617.21) (UDU36749.1) of SARs-CoV-2 with 3. reference strain (Wuhan) (YP_009724393.1)

4. **Nucleocapsid Phosphoprotein**

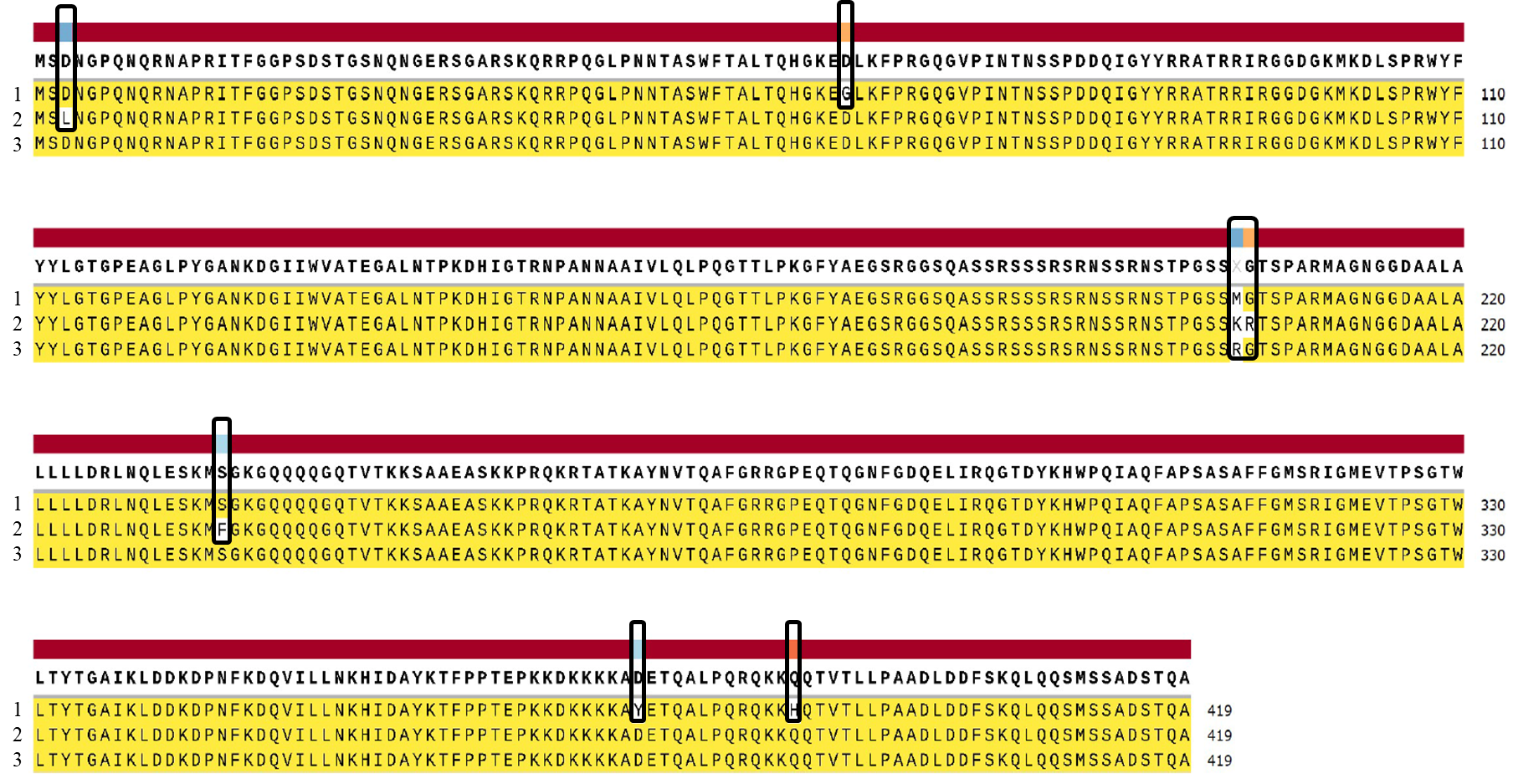

Figure 3: Schematic view of contigs of the Nucleocapsid phosphoprotein of 1. Alpha Variant (B.1.1.7) (UDQ41846.1) and 2. Delta Variant (B.1.617.21) (UDU36754.1) of SARs-CoV-2 with 3. reference strain (Wuhan) (YP_009724397.2)

5. **ORF10 protein**

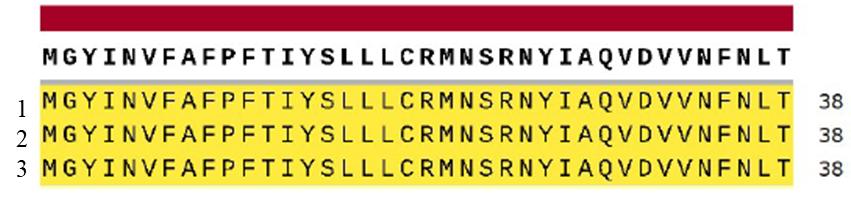

Figure 4: Schematic view of contigs of the ORF10 protein of 1. Alpha Variant (B.1.1.7) (UDQ41847.1) and 2. Delta Variant (B.1.617.21) (UDU36755.1) of SARs-CoV-2 with 3. reference Strain (Wuhan) (YP_009725255.1)

7. **ORF1a polyprotein**

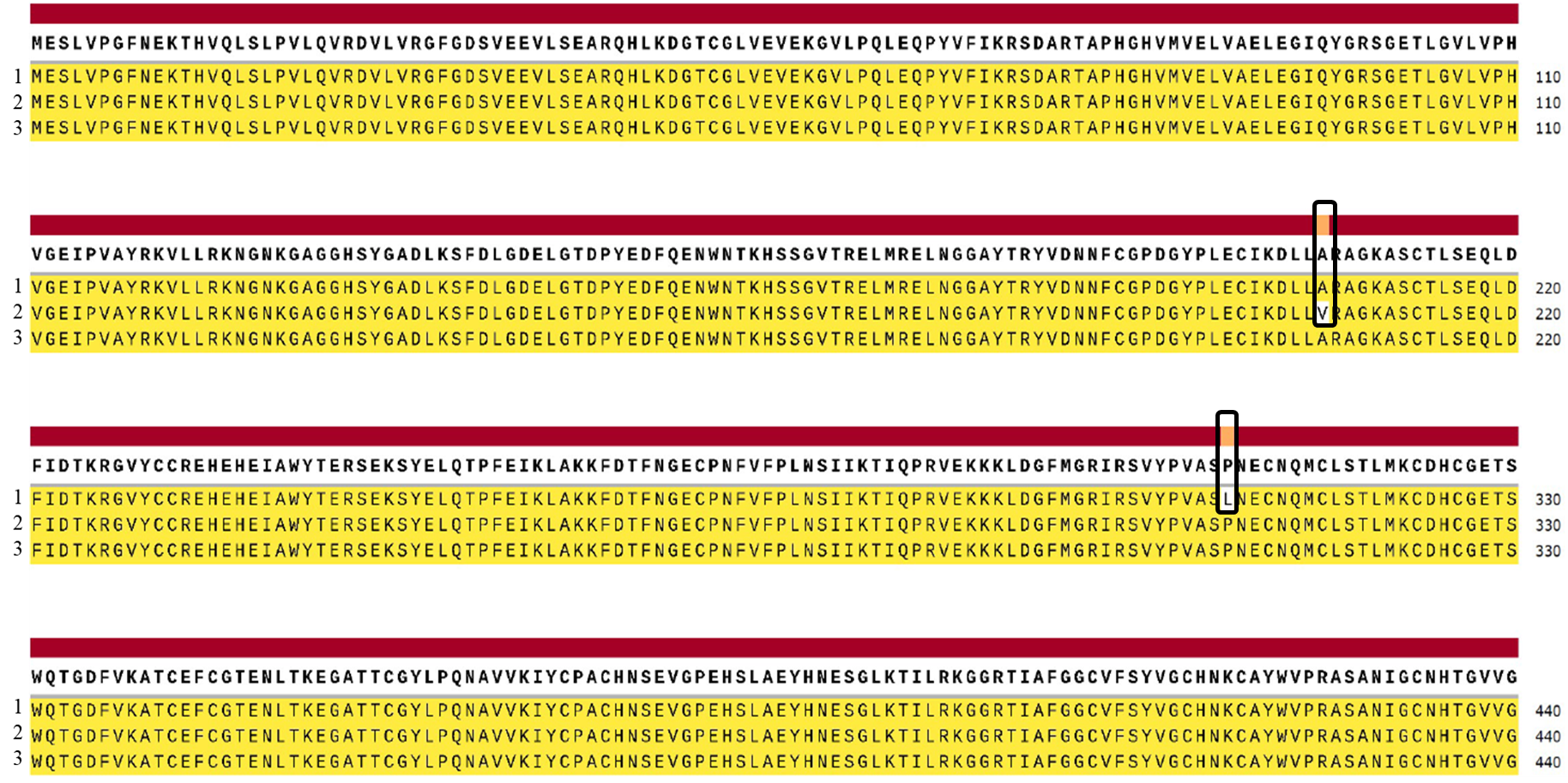

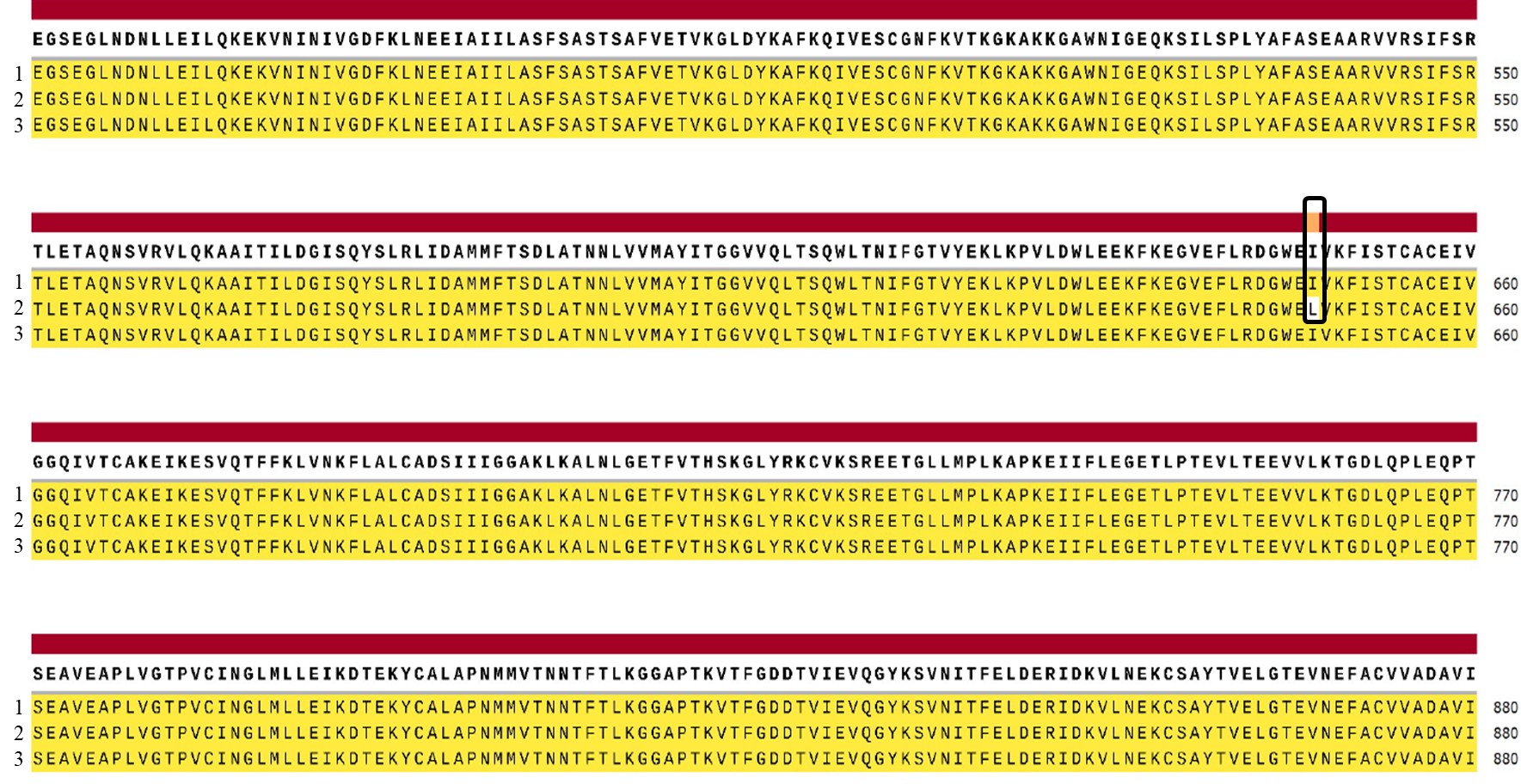

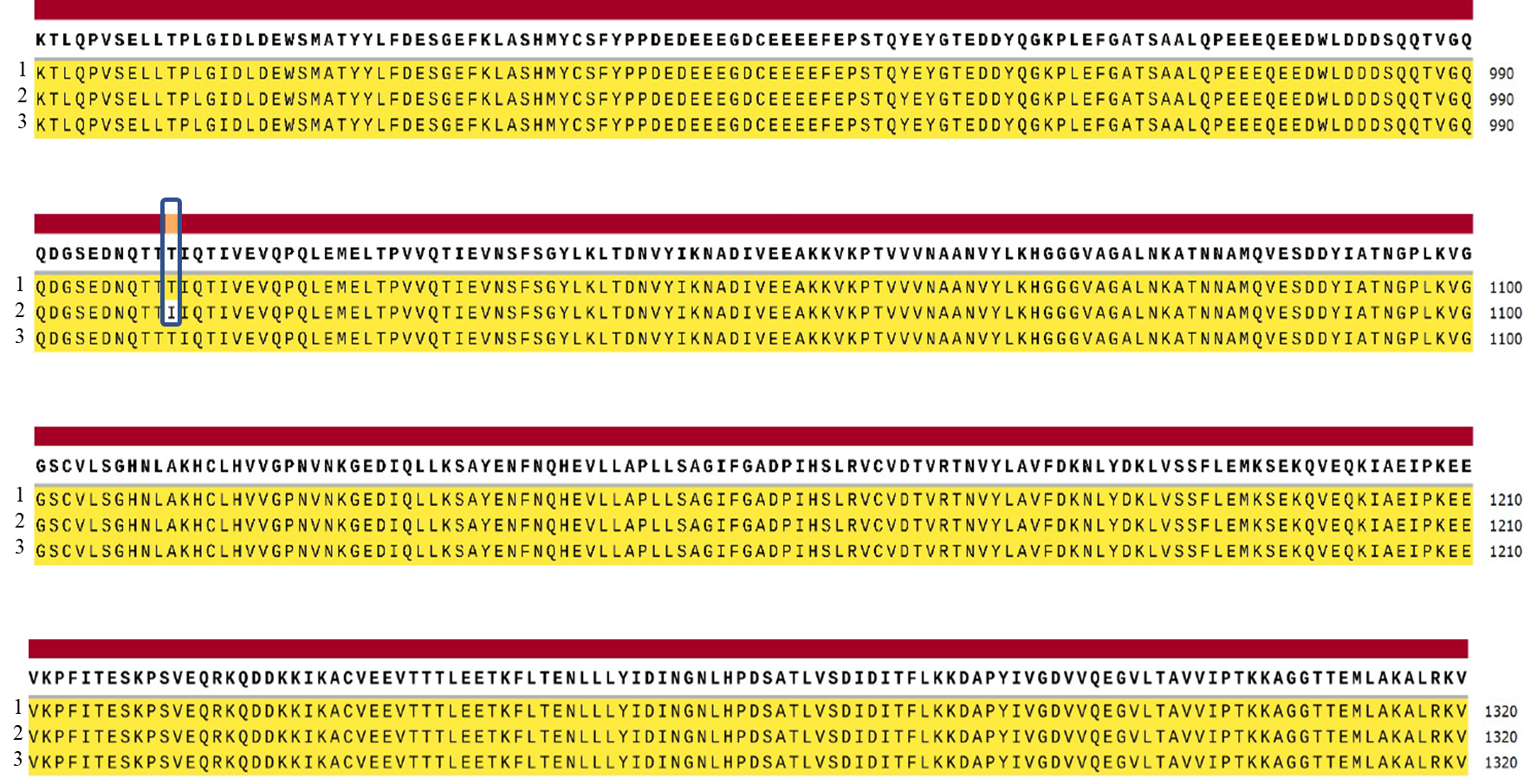

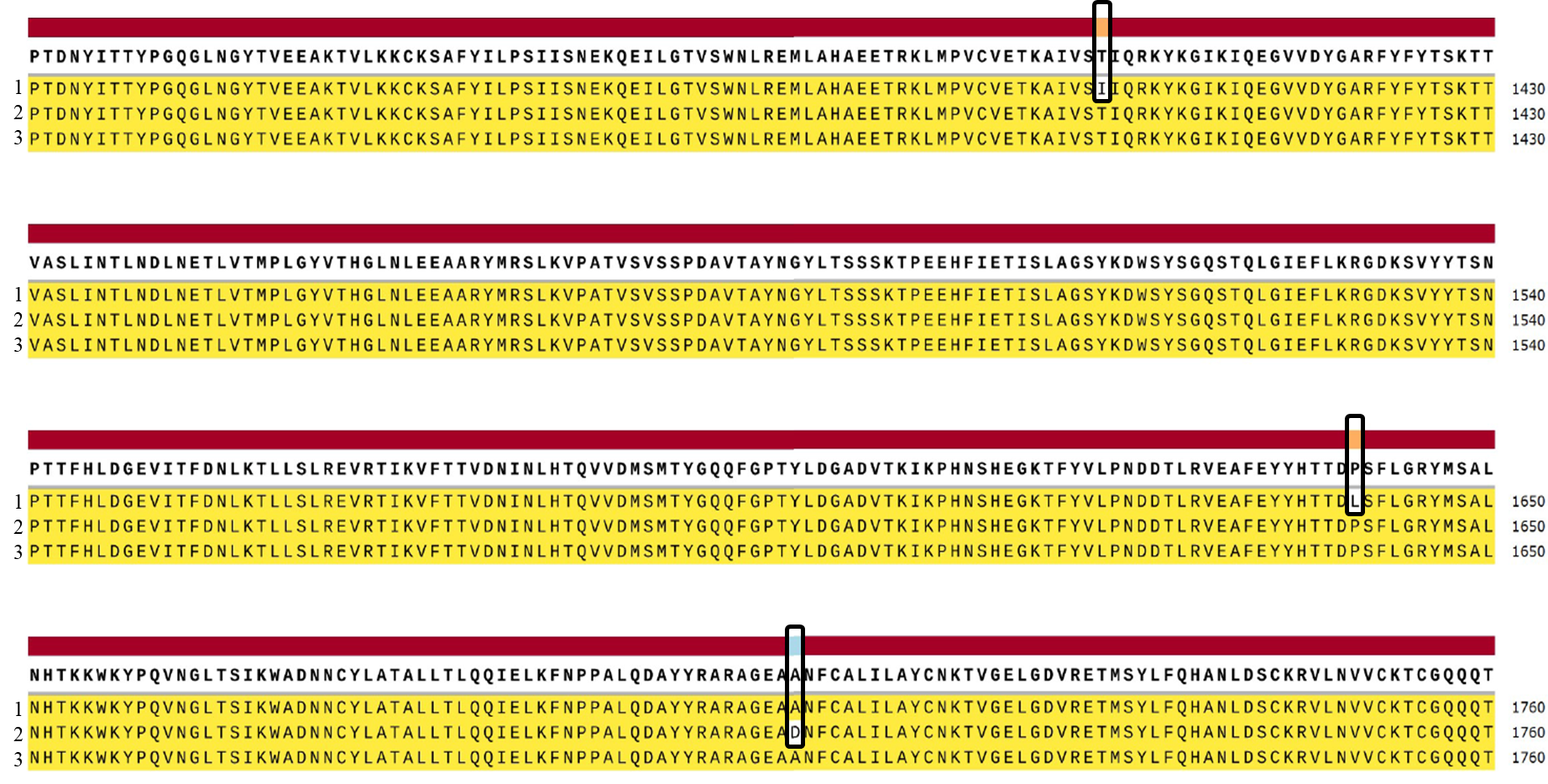

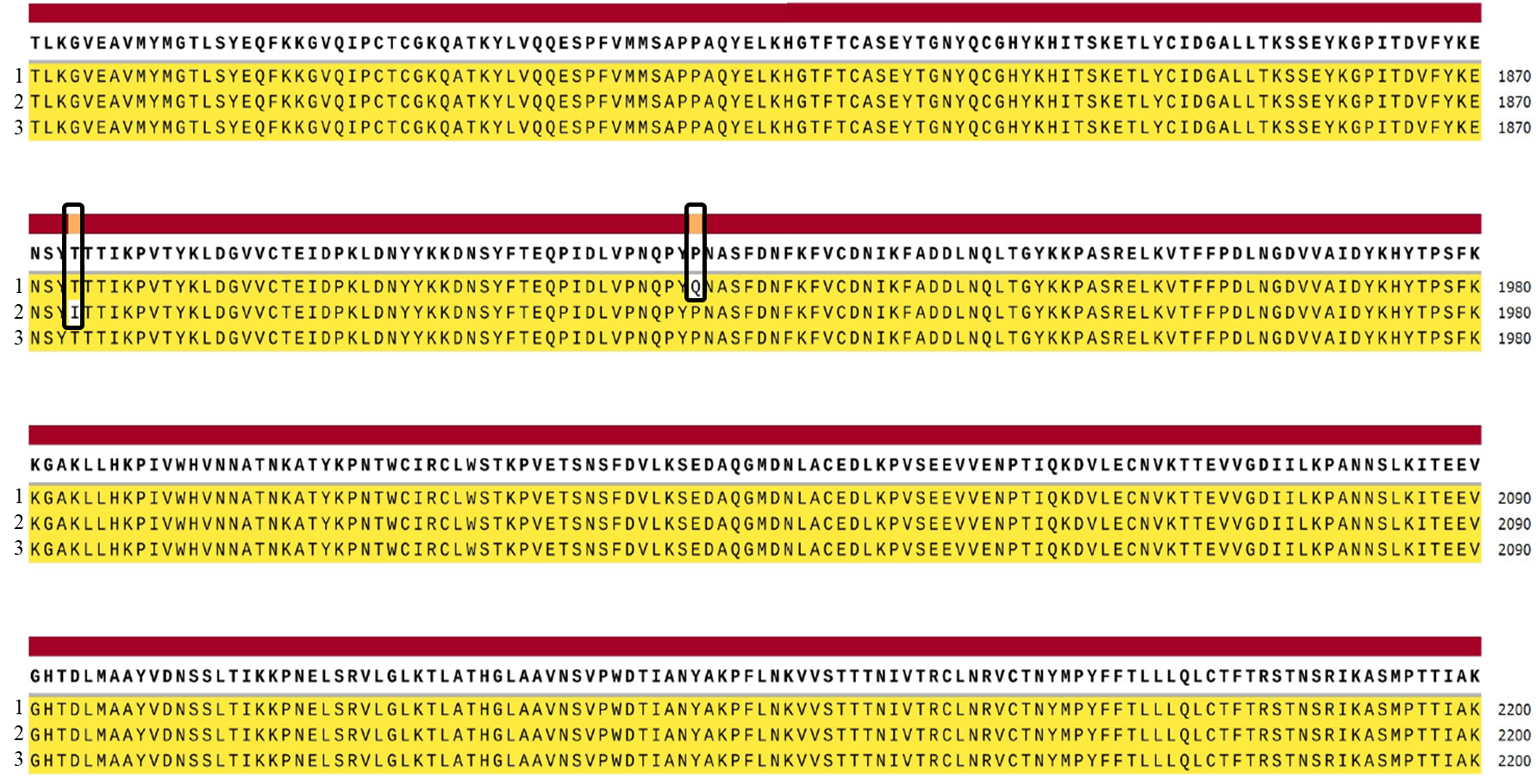

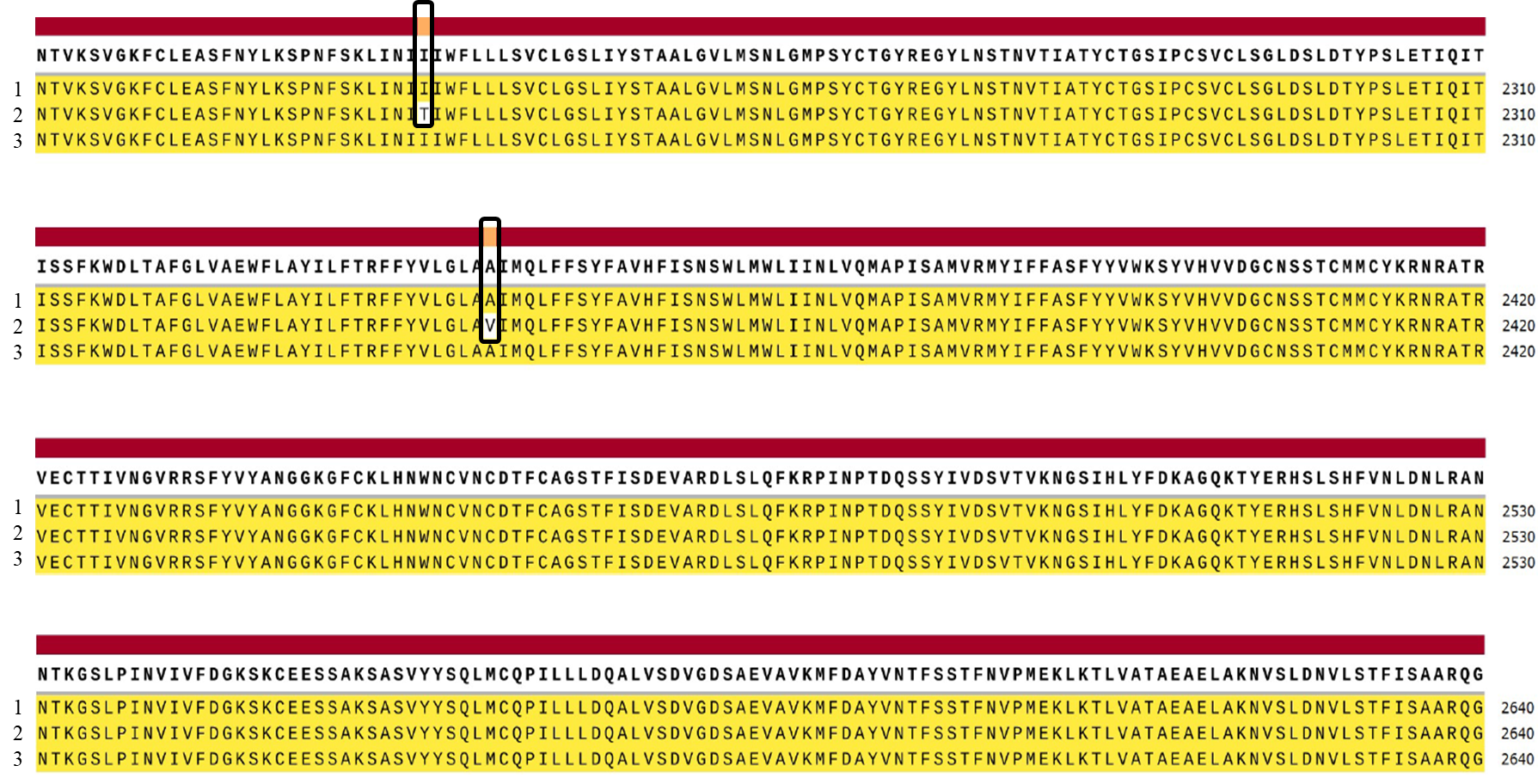

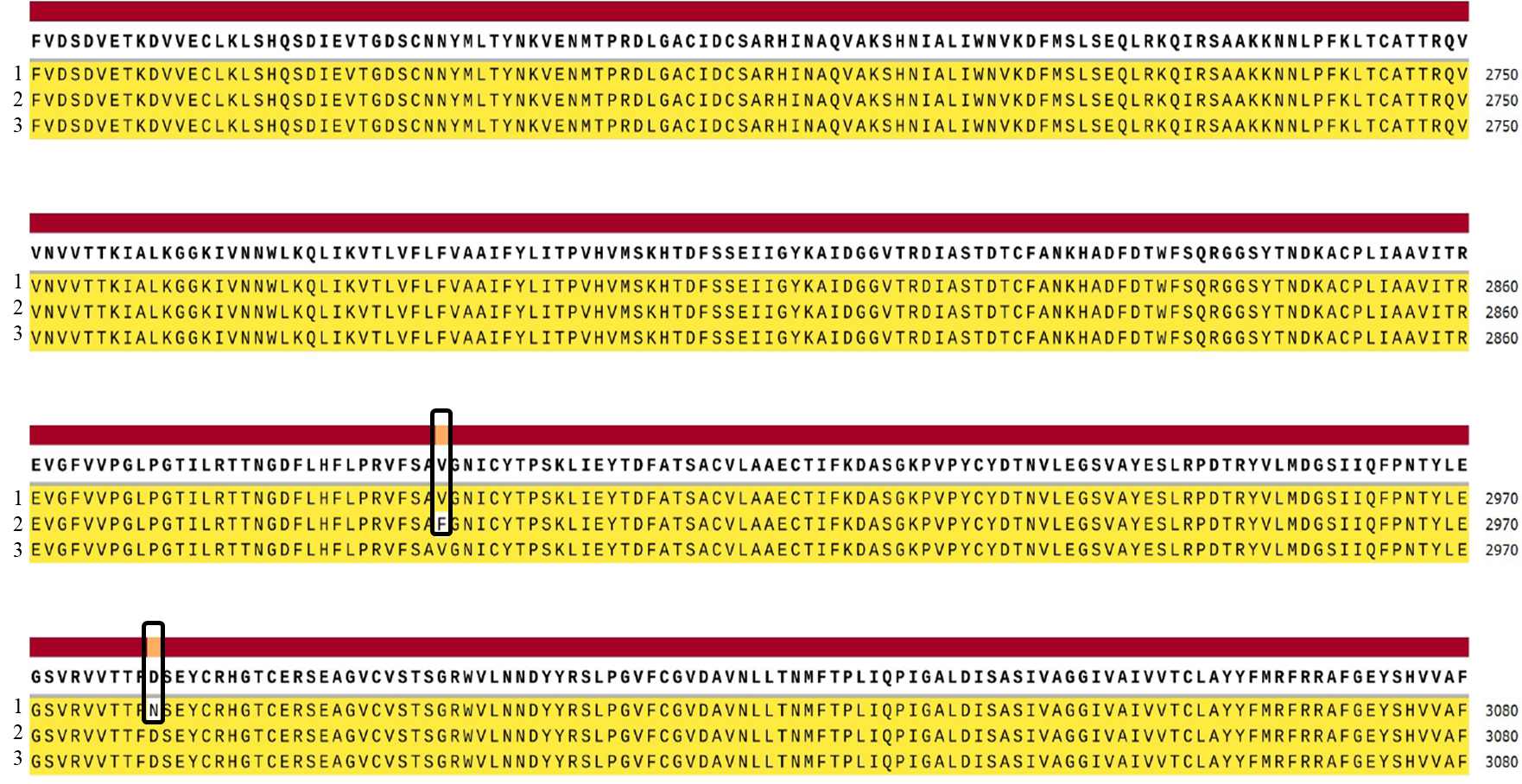

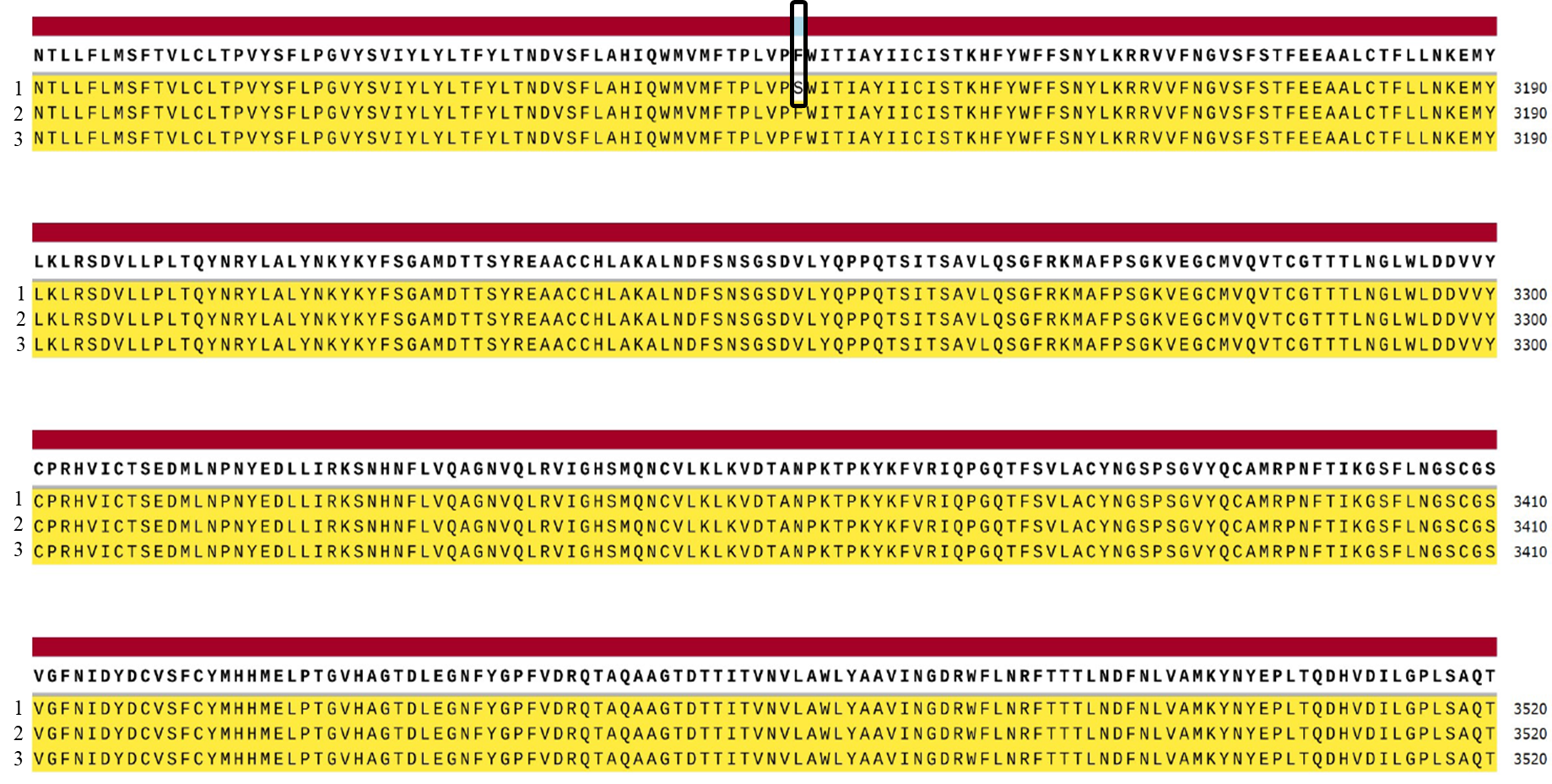

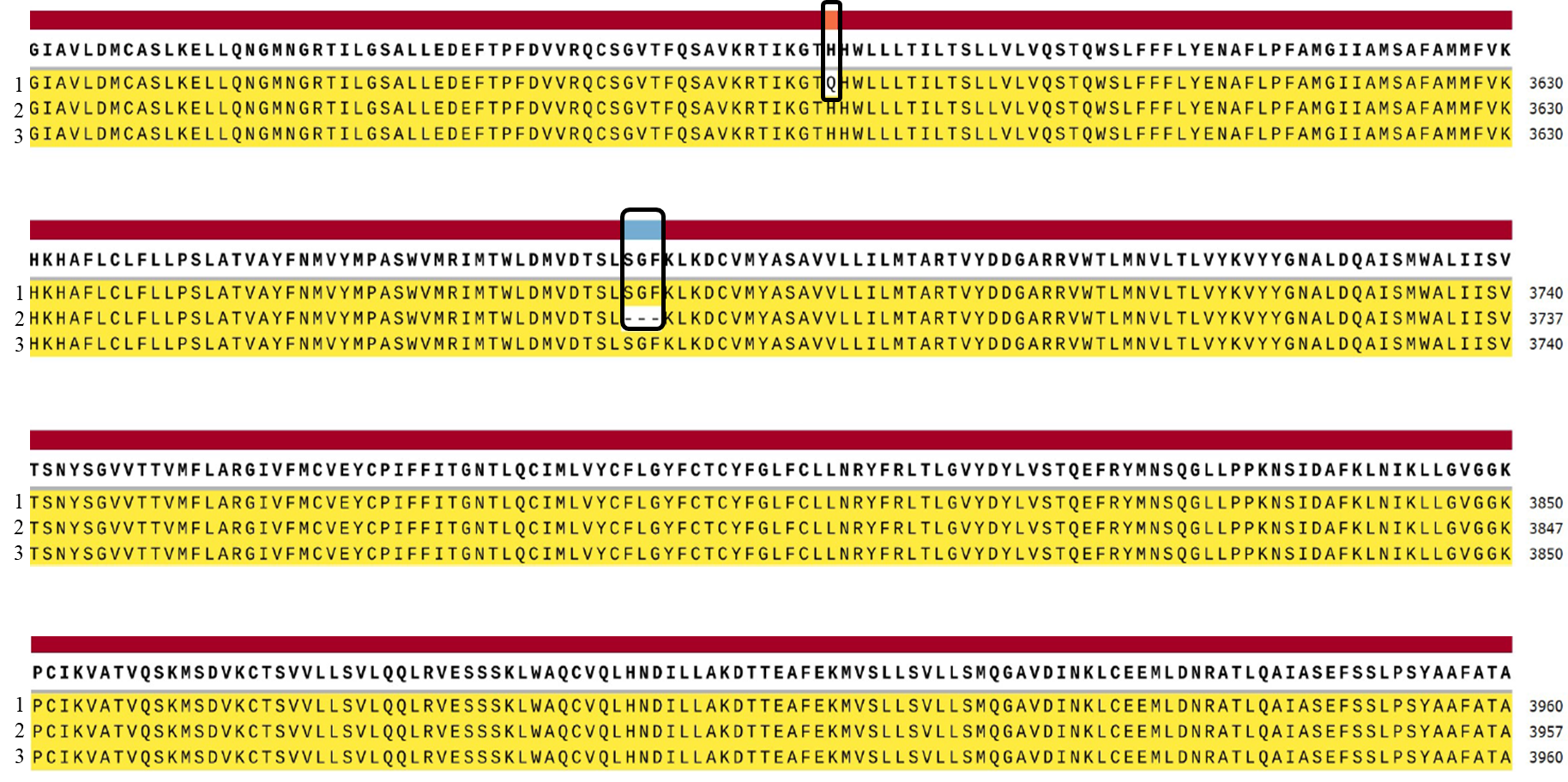

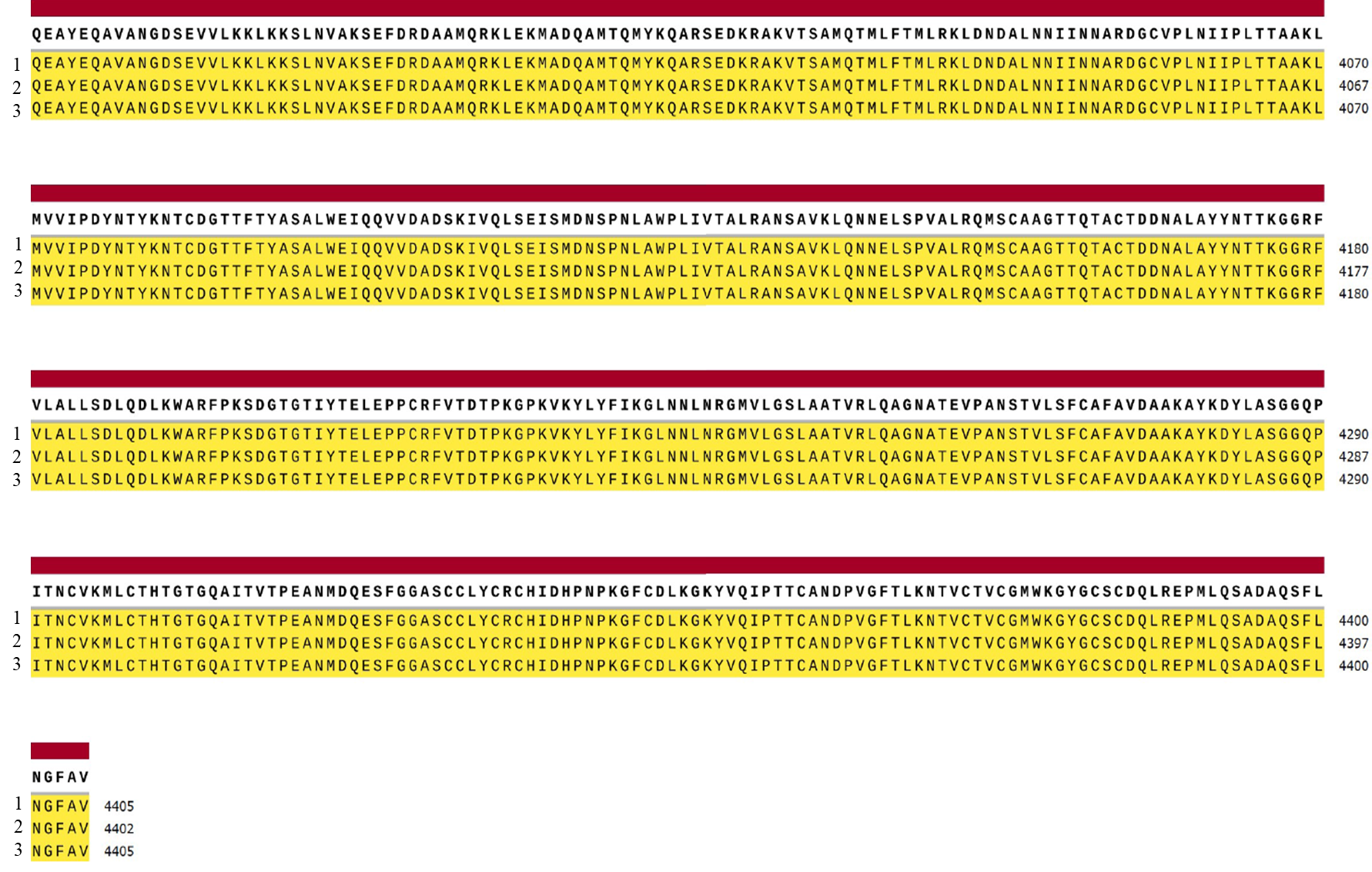

Figure 5Schematic view of contigs of the ORF1a polyprotein of 1. Alpha Variant (B.1.1.7) (UDQ41837.1) and 2. Delta Variant (B.1.617.21) (UDU36745.1) of SARs-CoV-2 with 3. reference Strain (Wuhan) (YP_009725295.1)

8. **ORF1ab polyprotein**

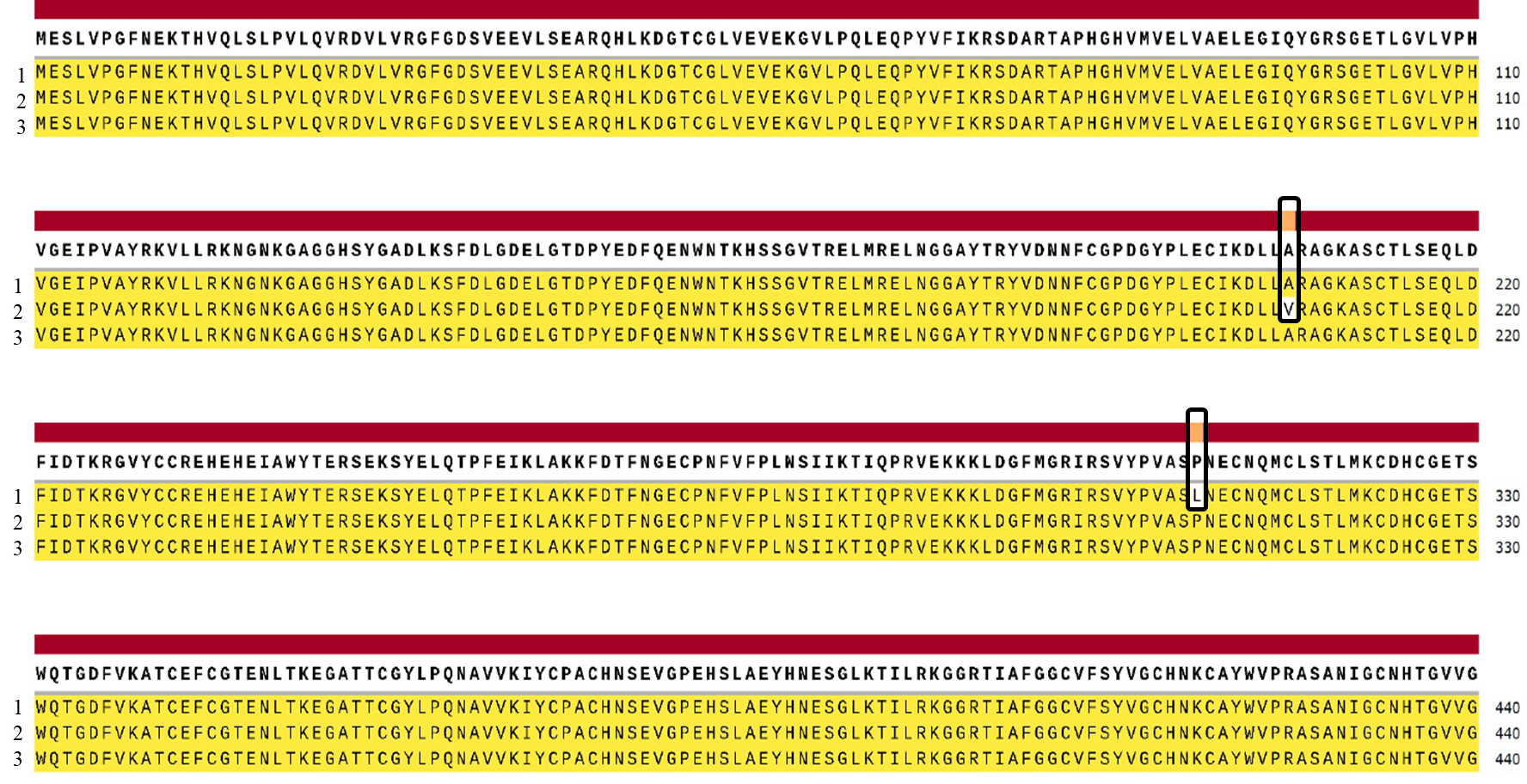

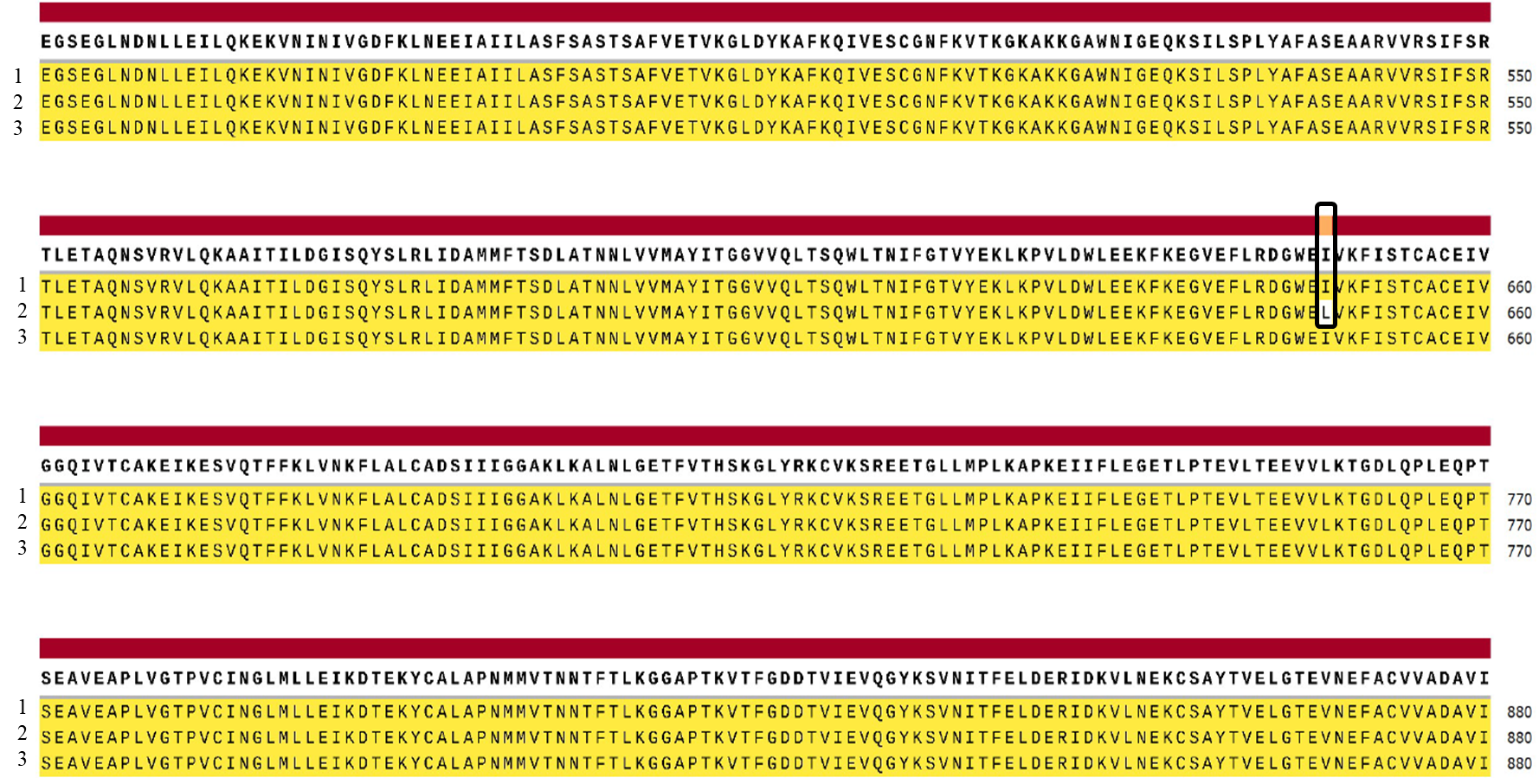

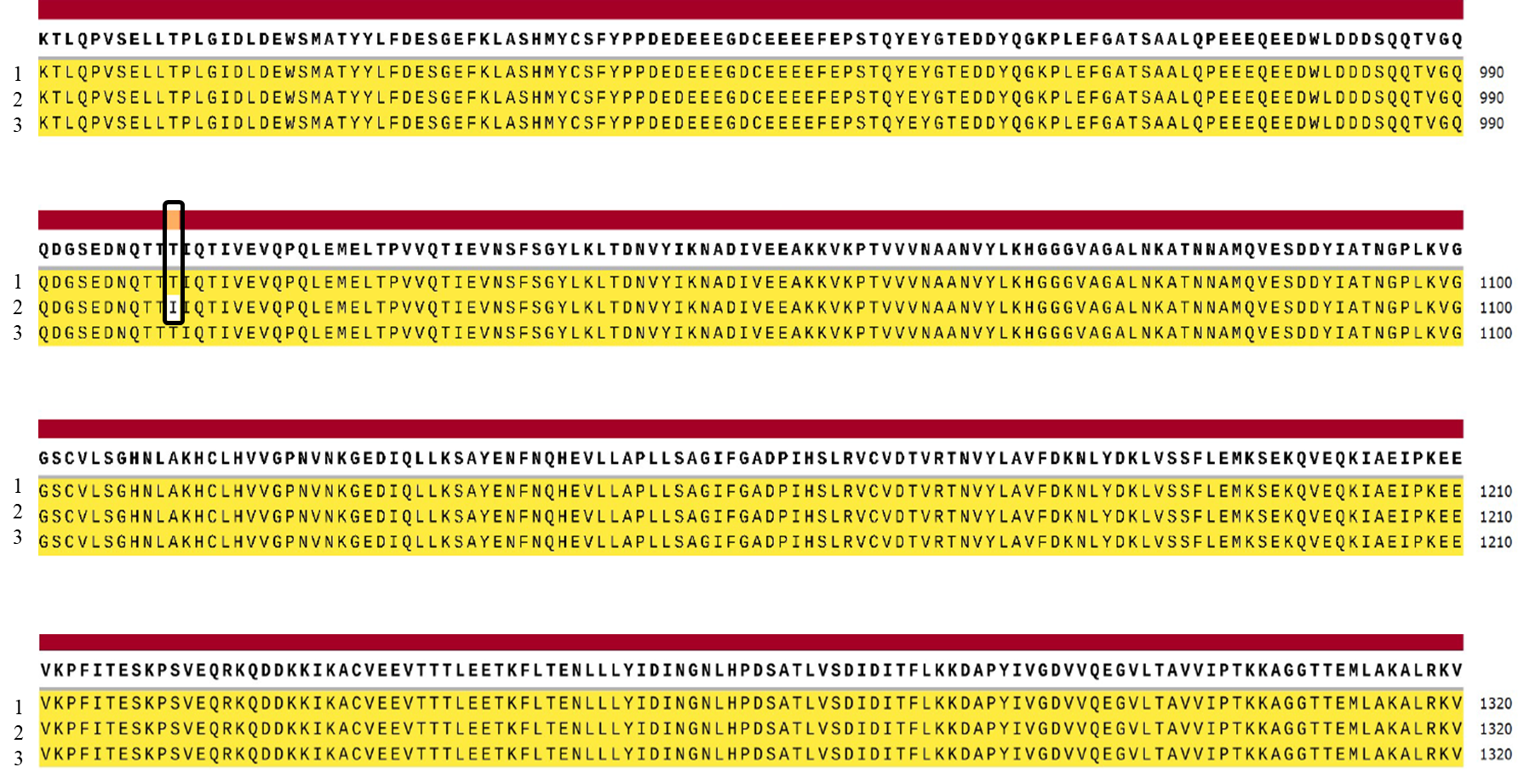

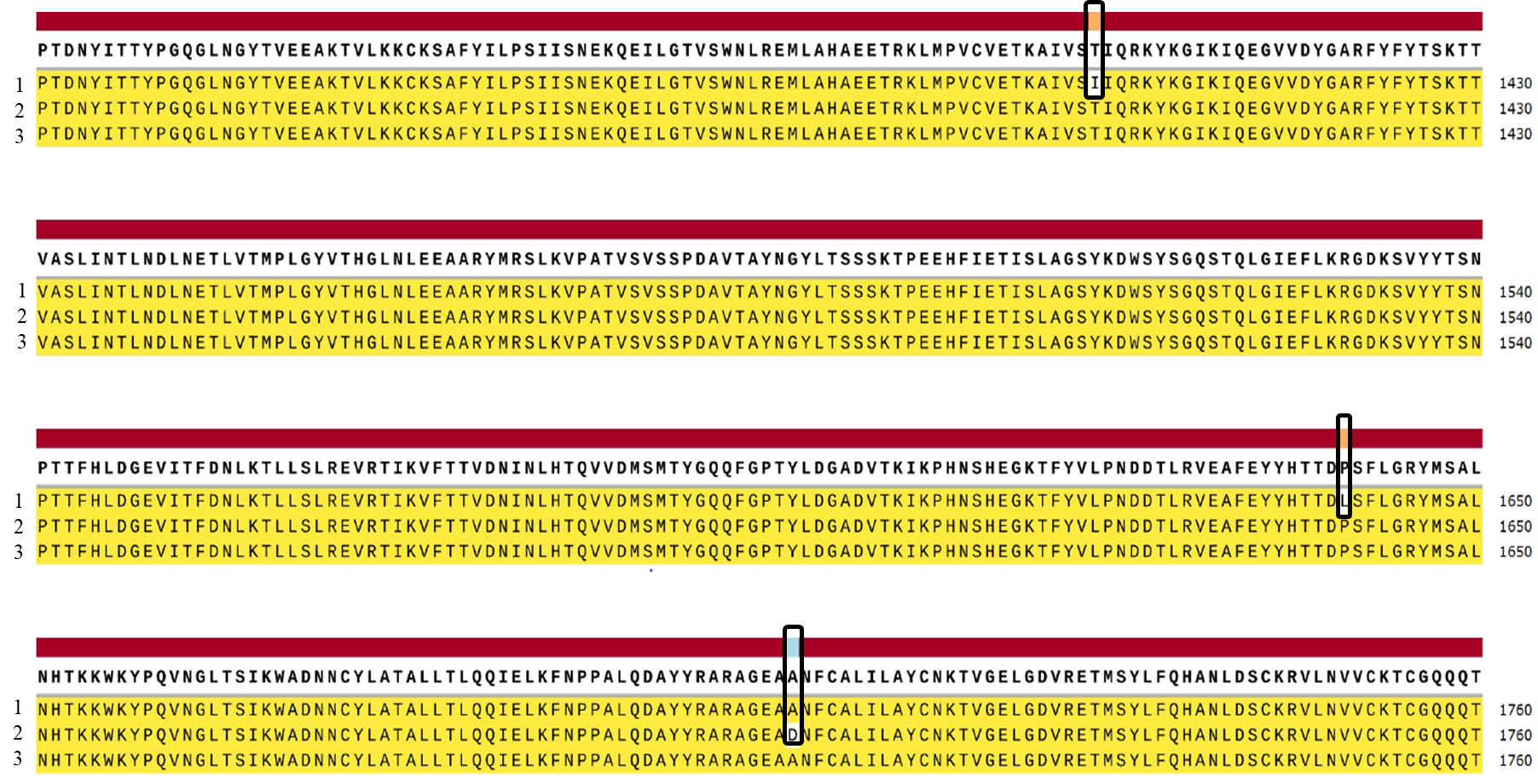

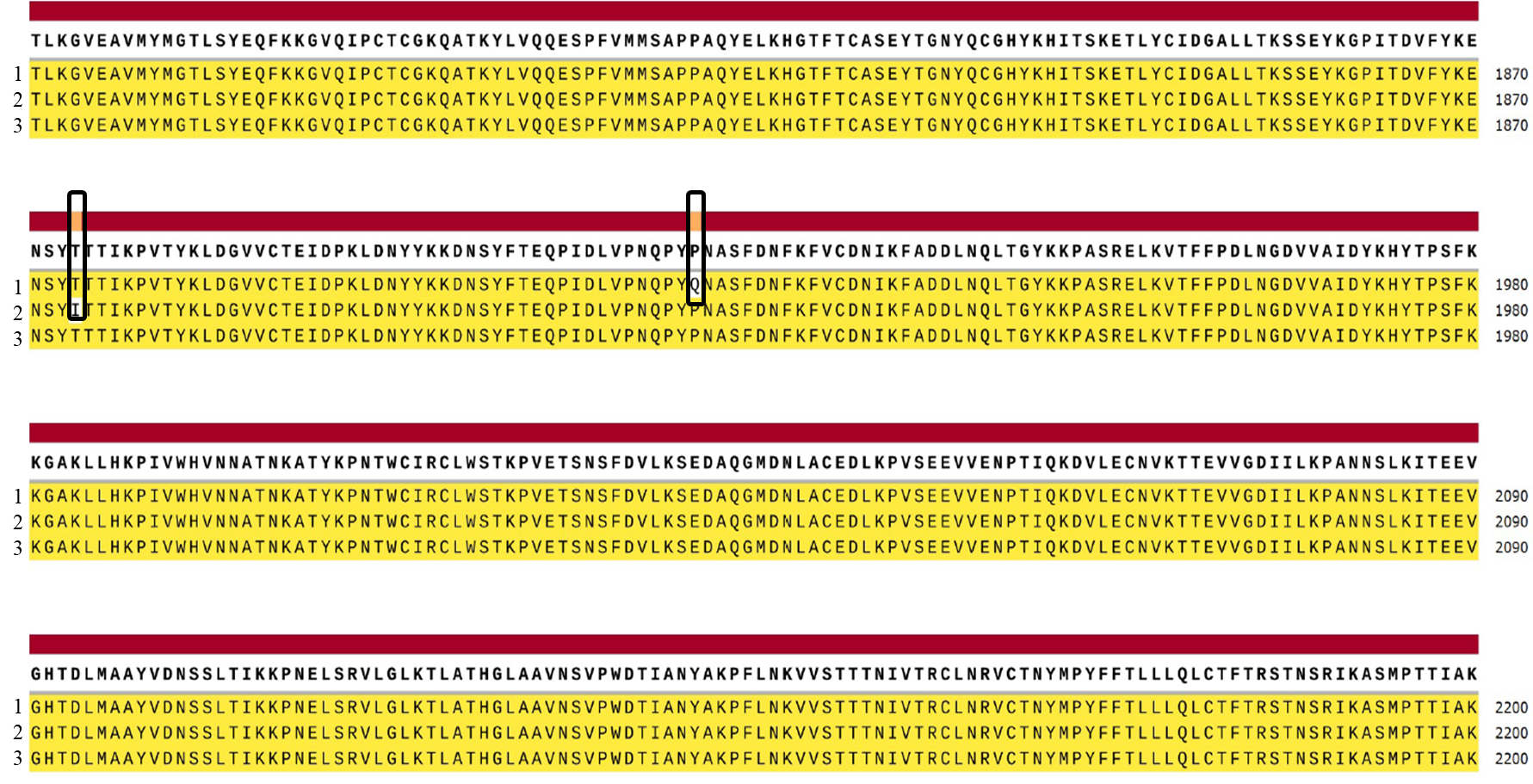

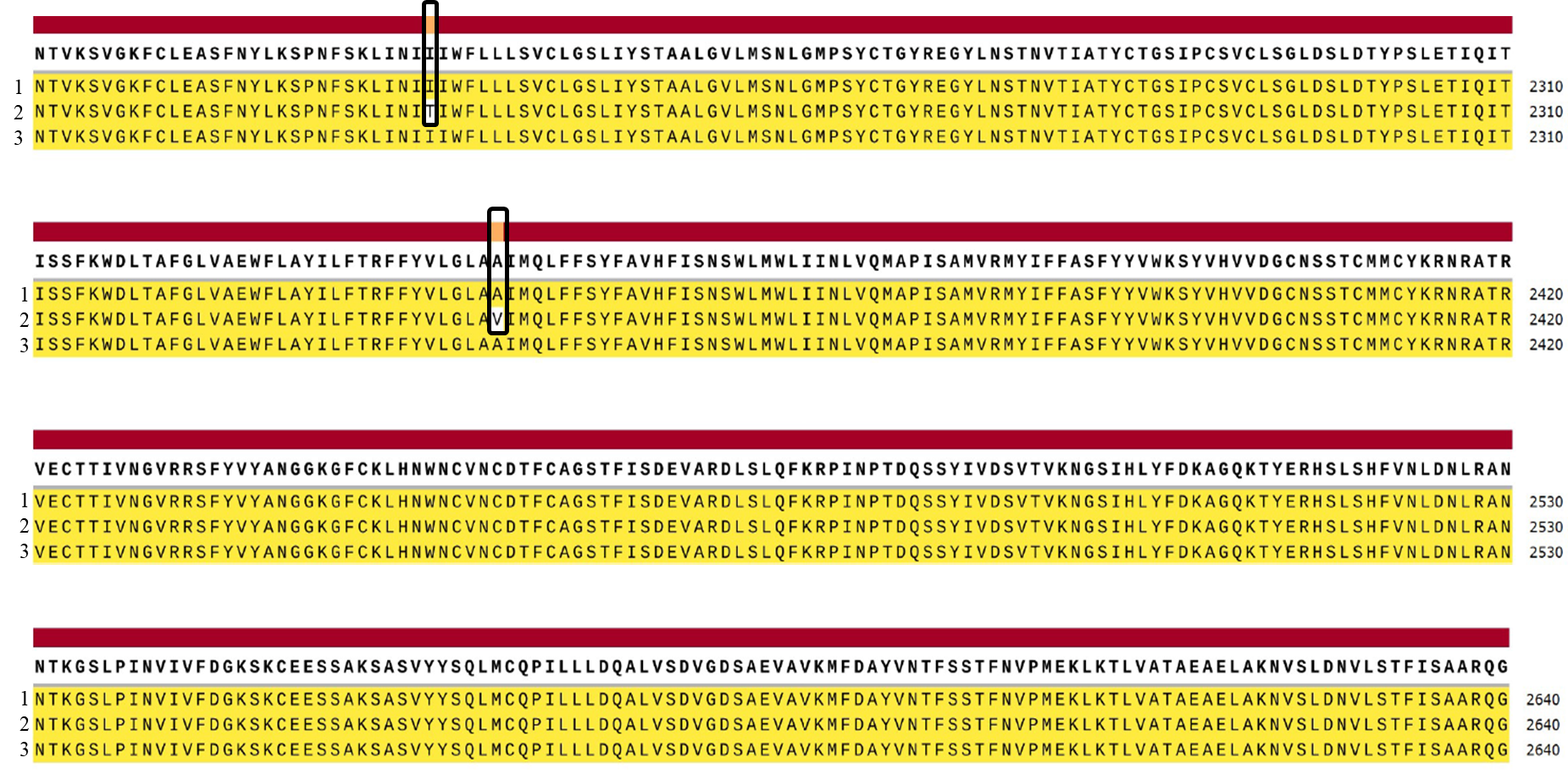

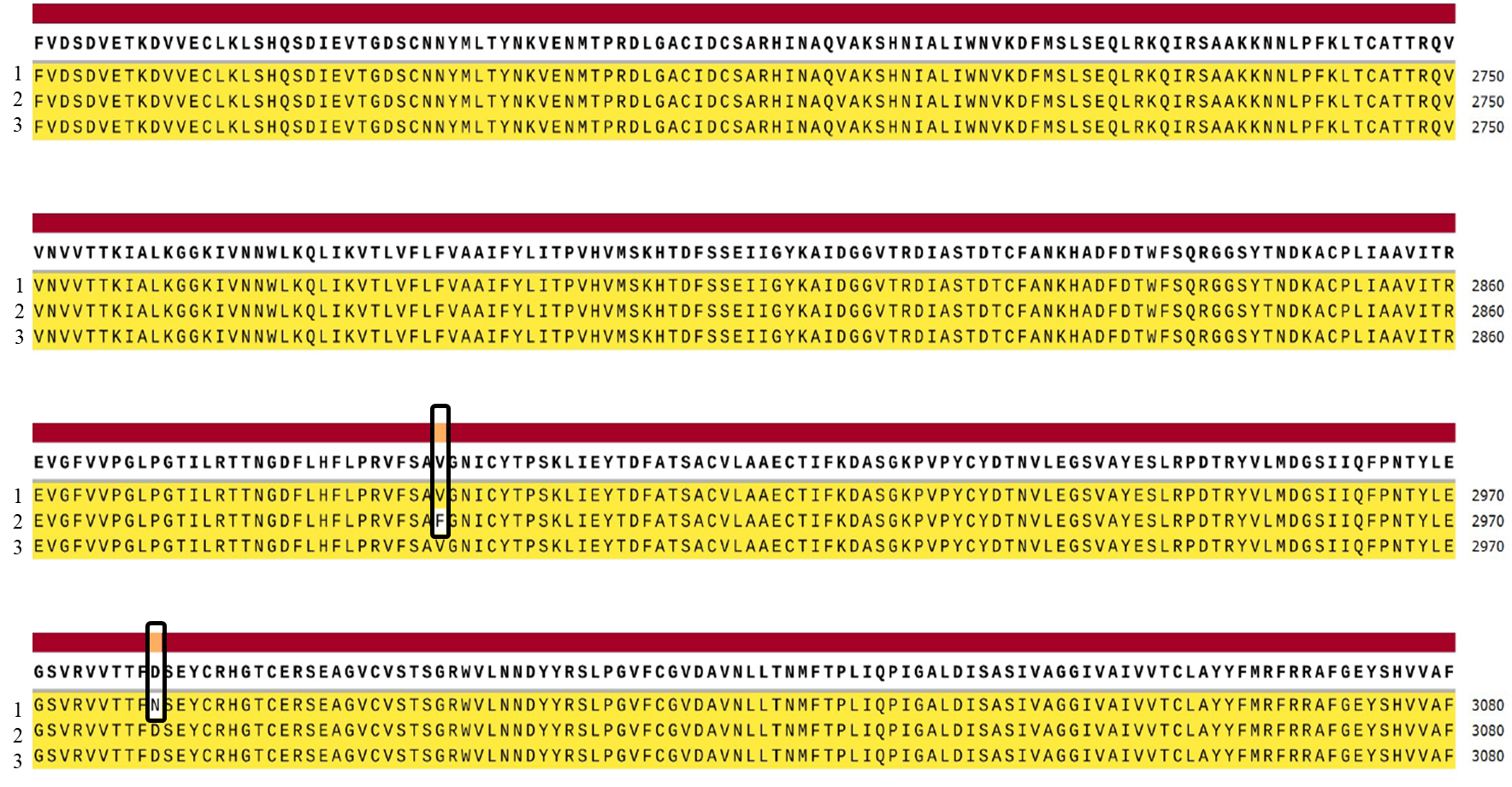

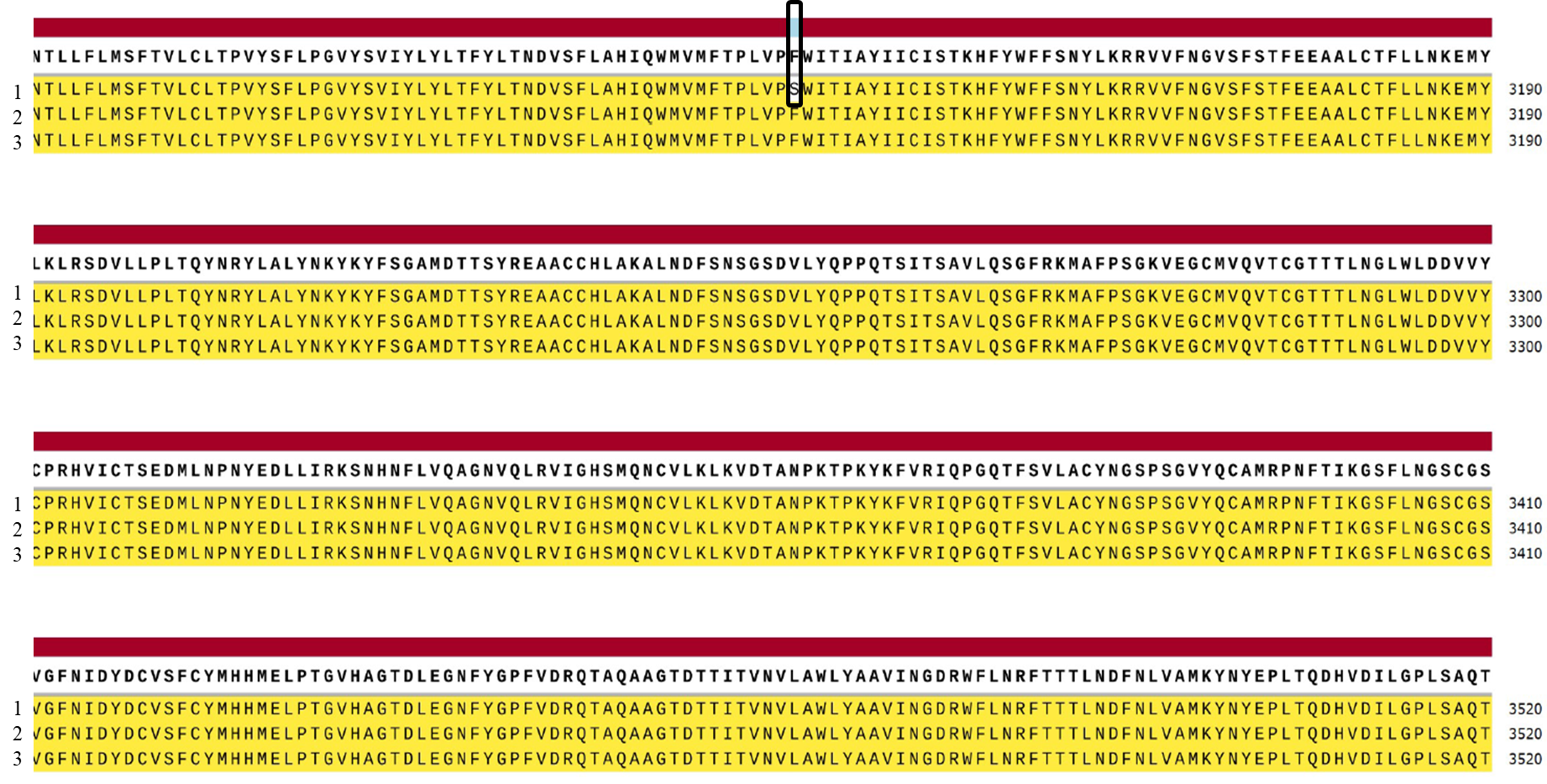

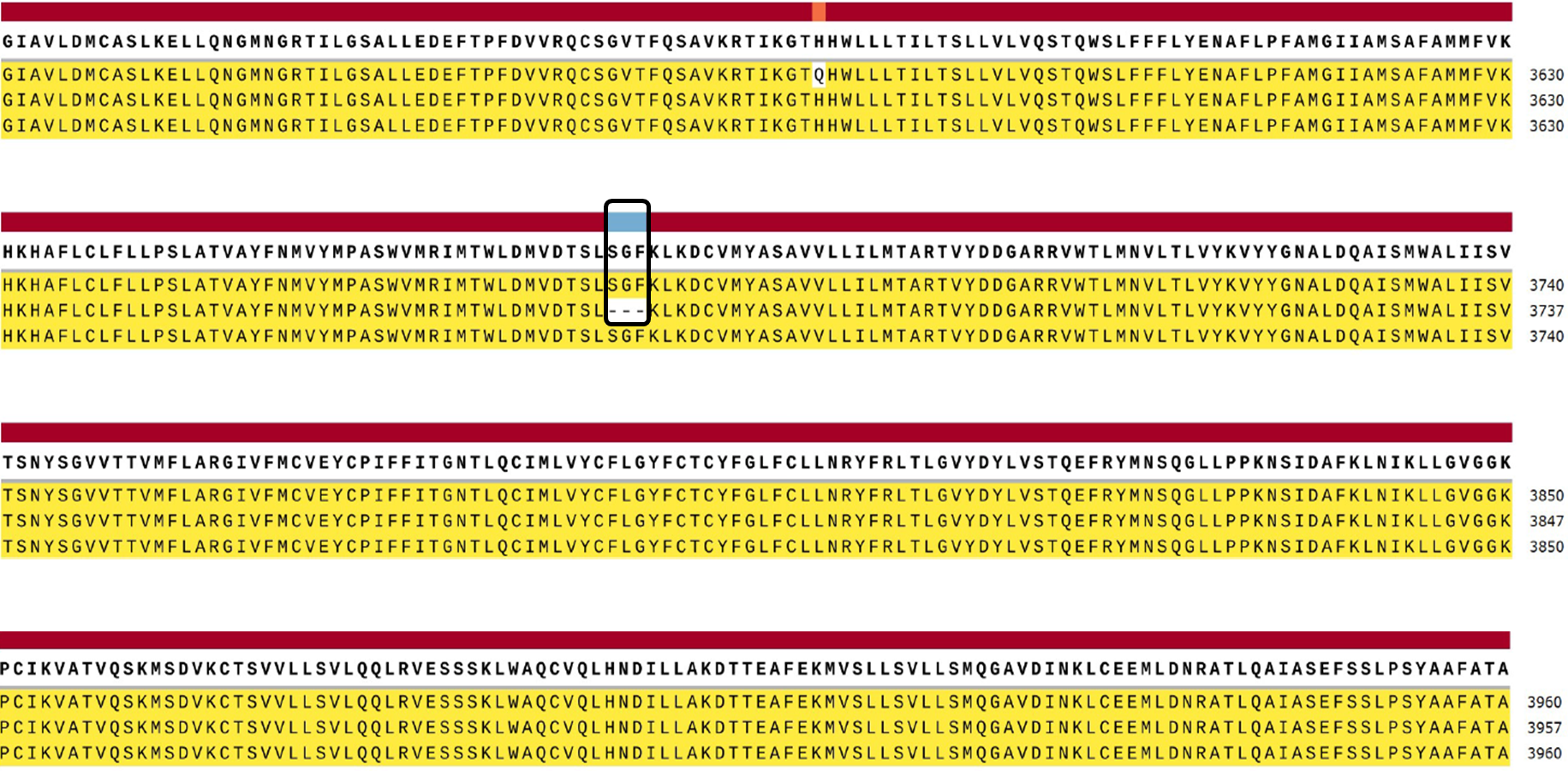

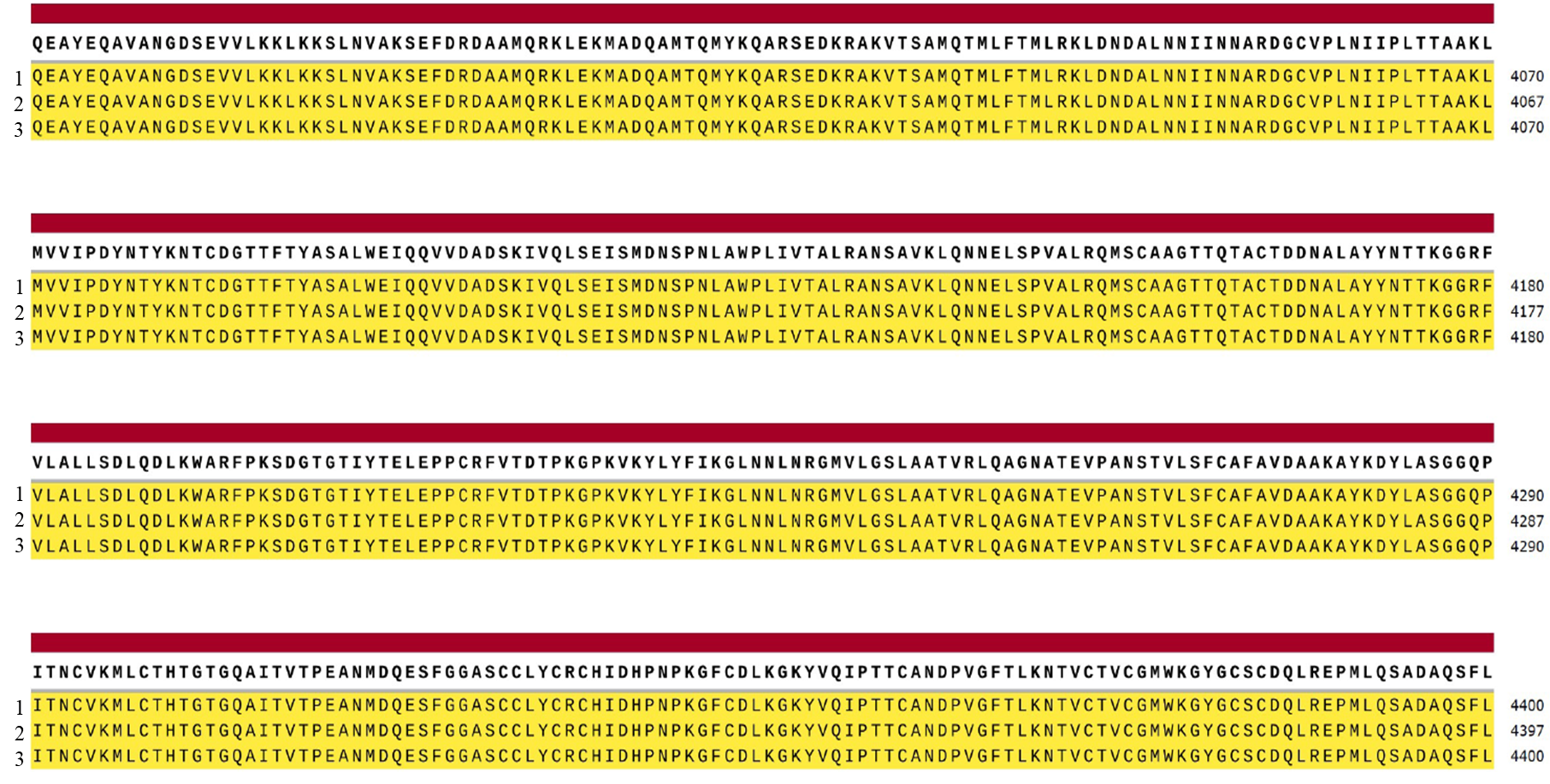

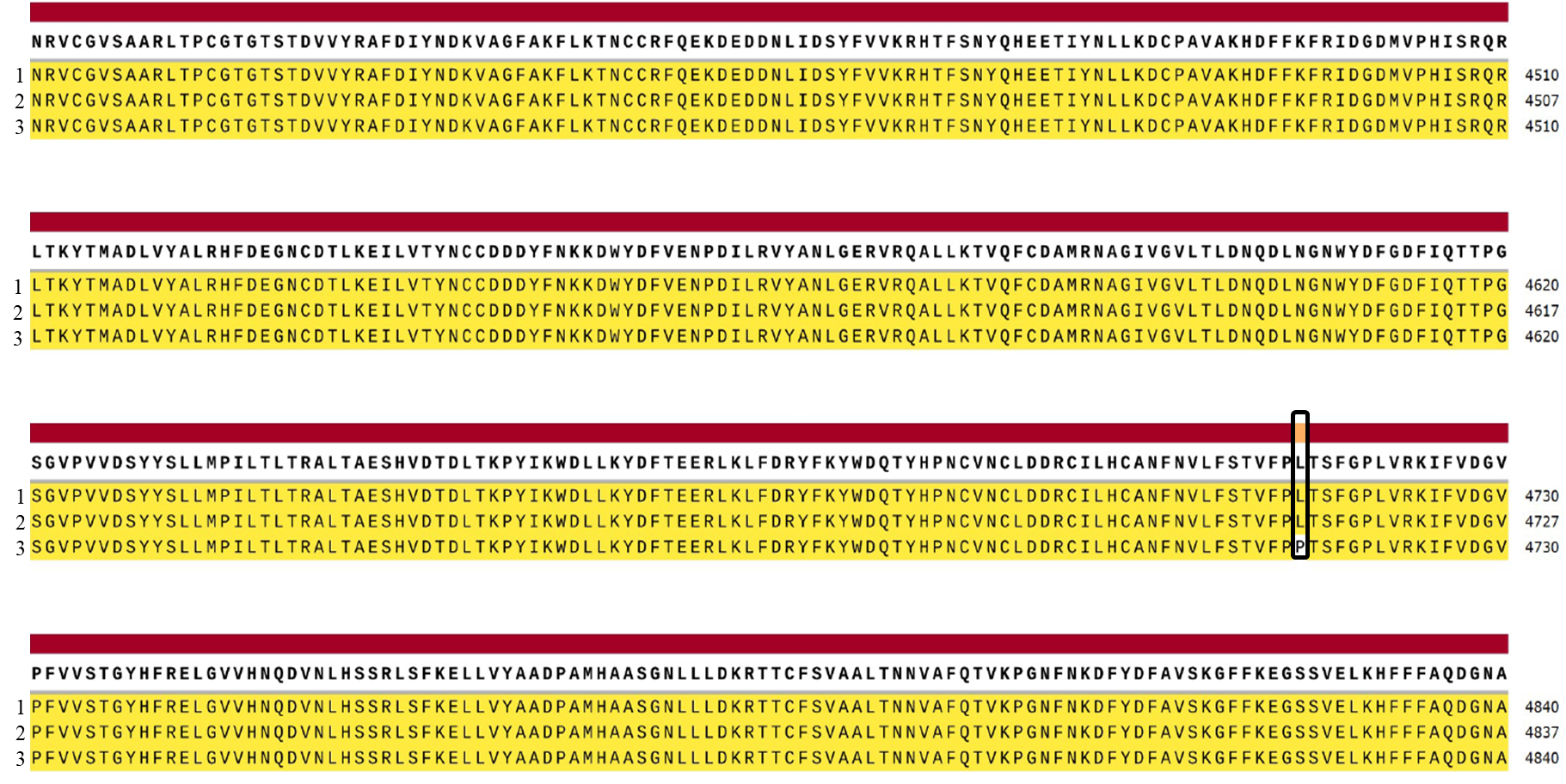

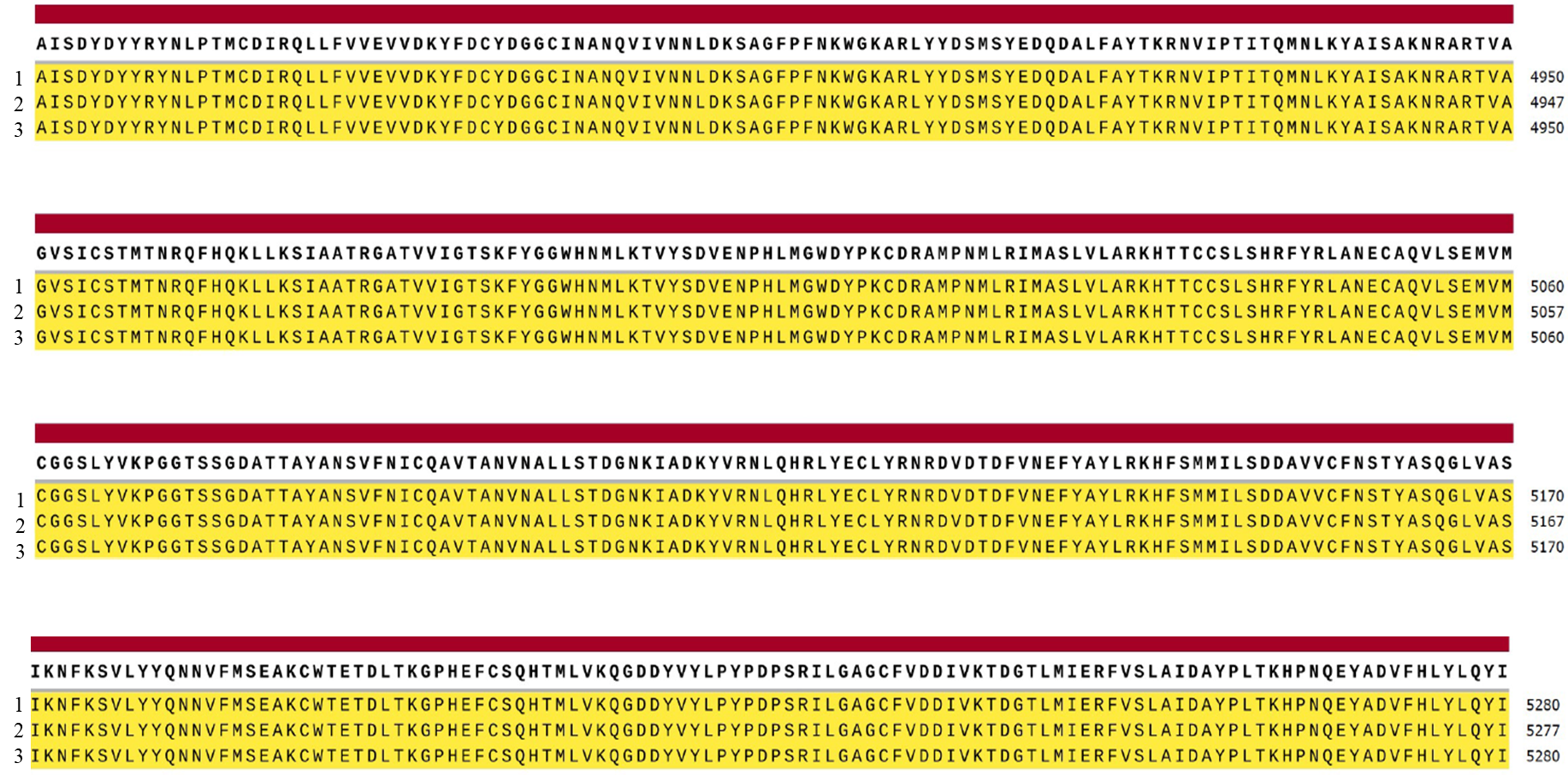

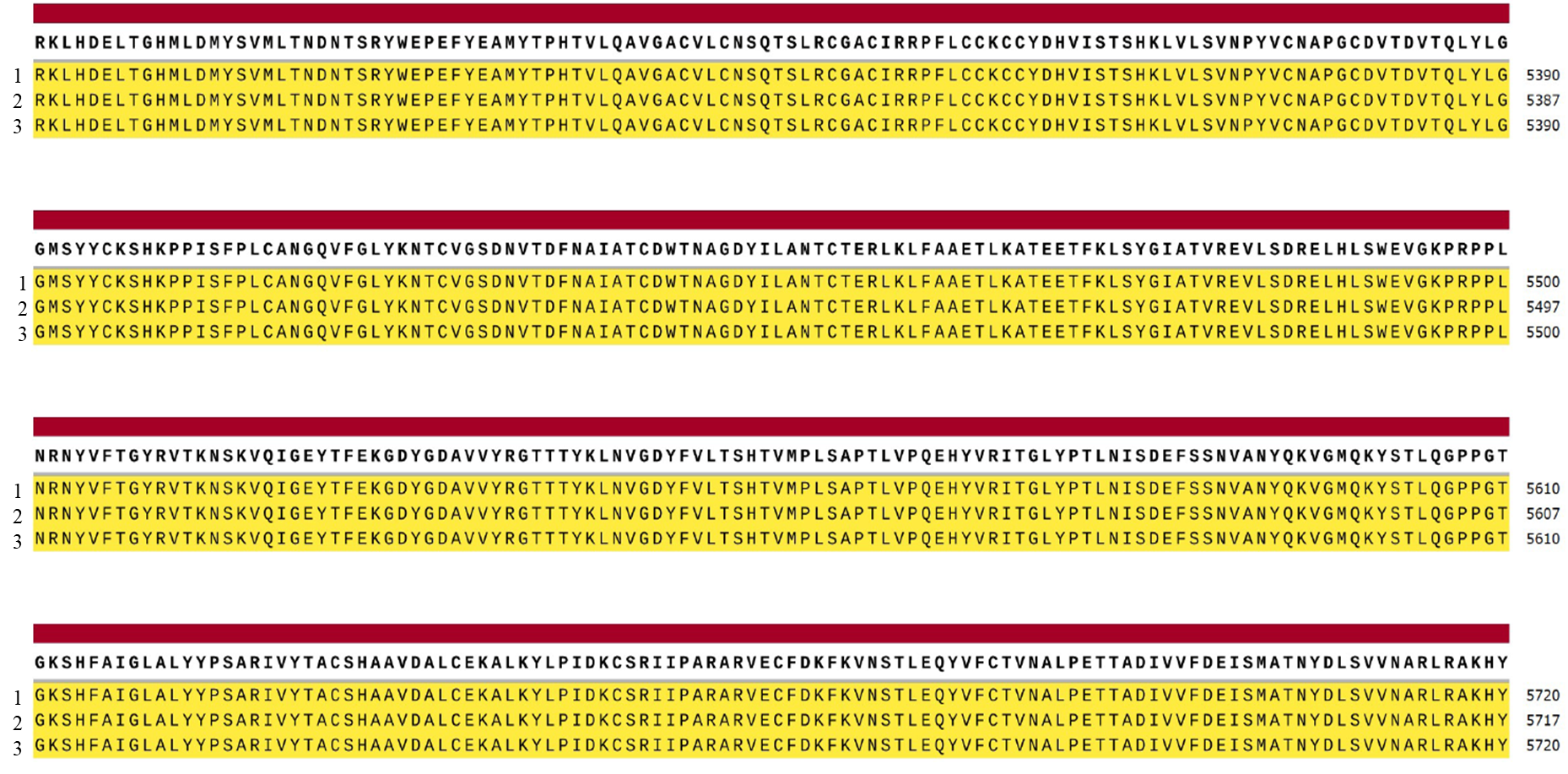

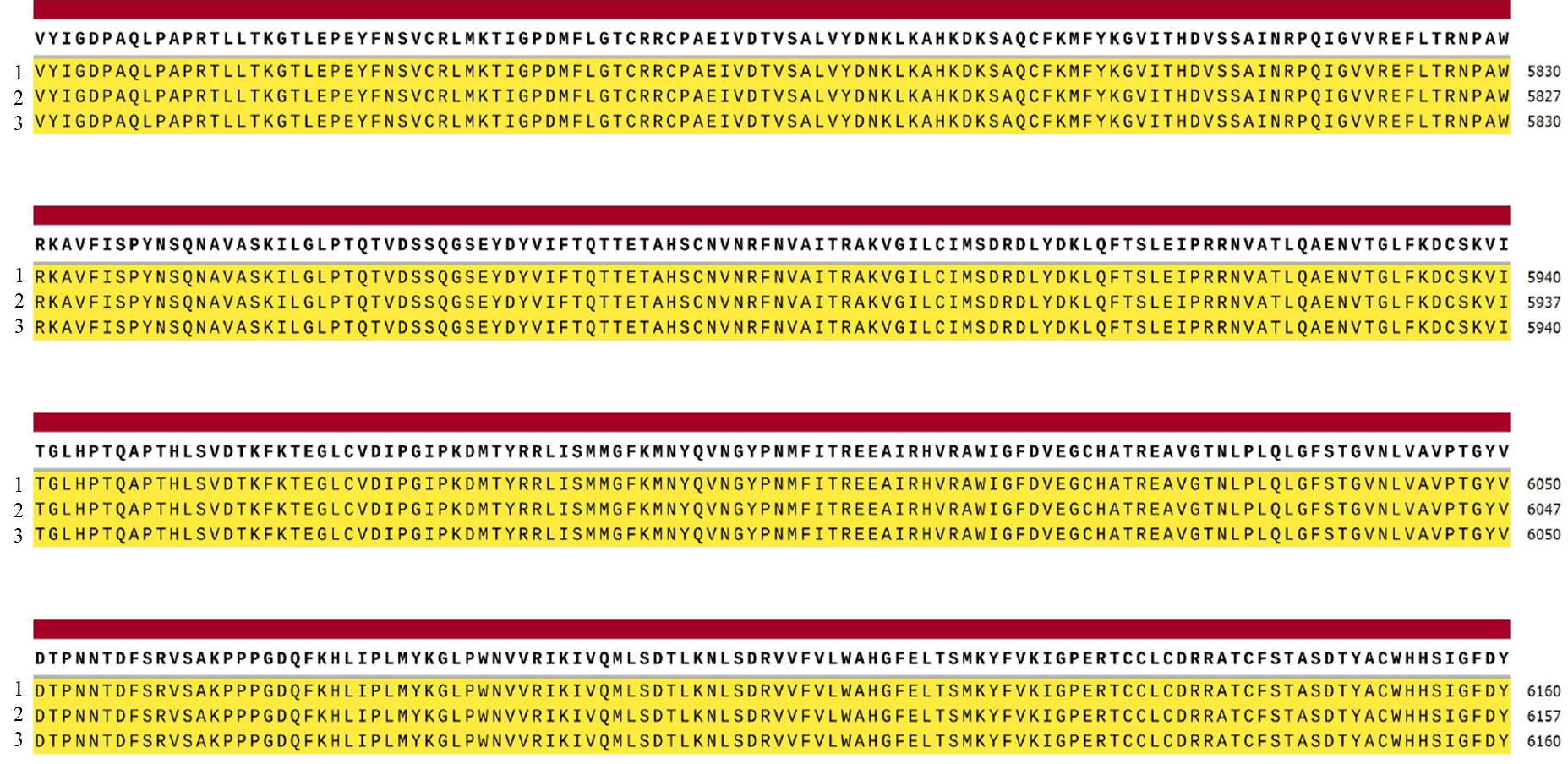

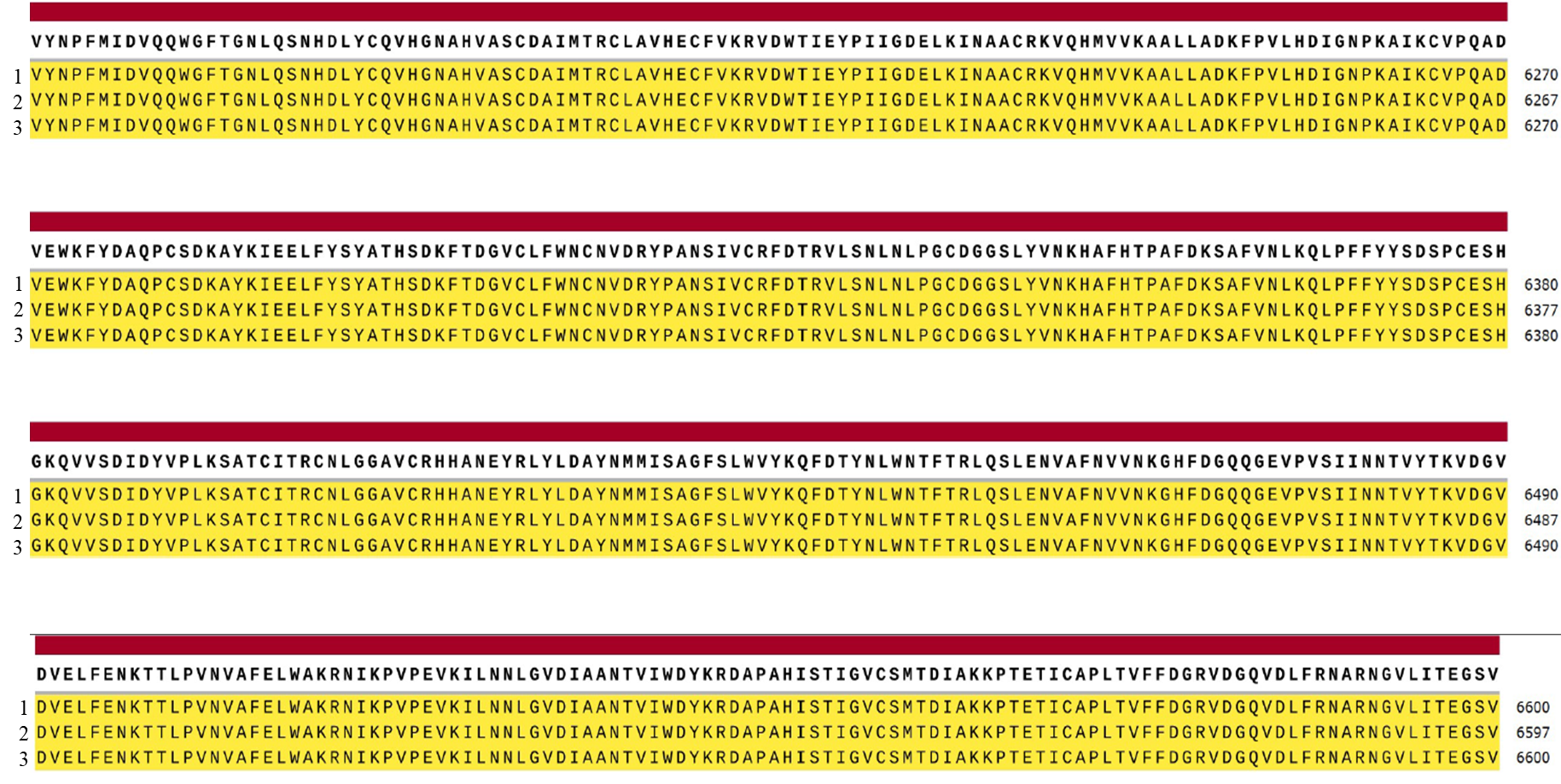

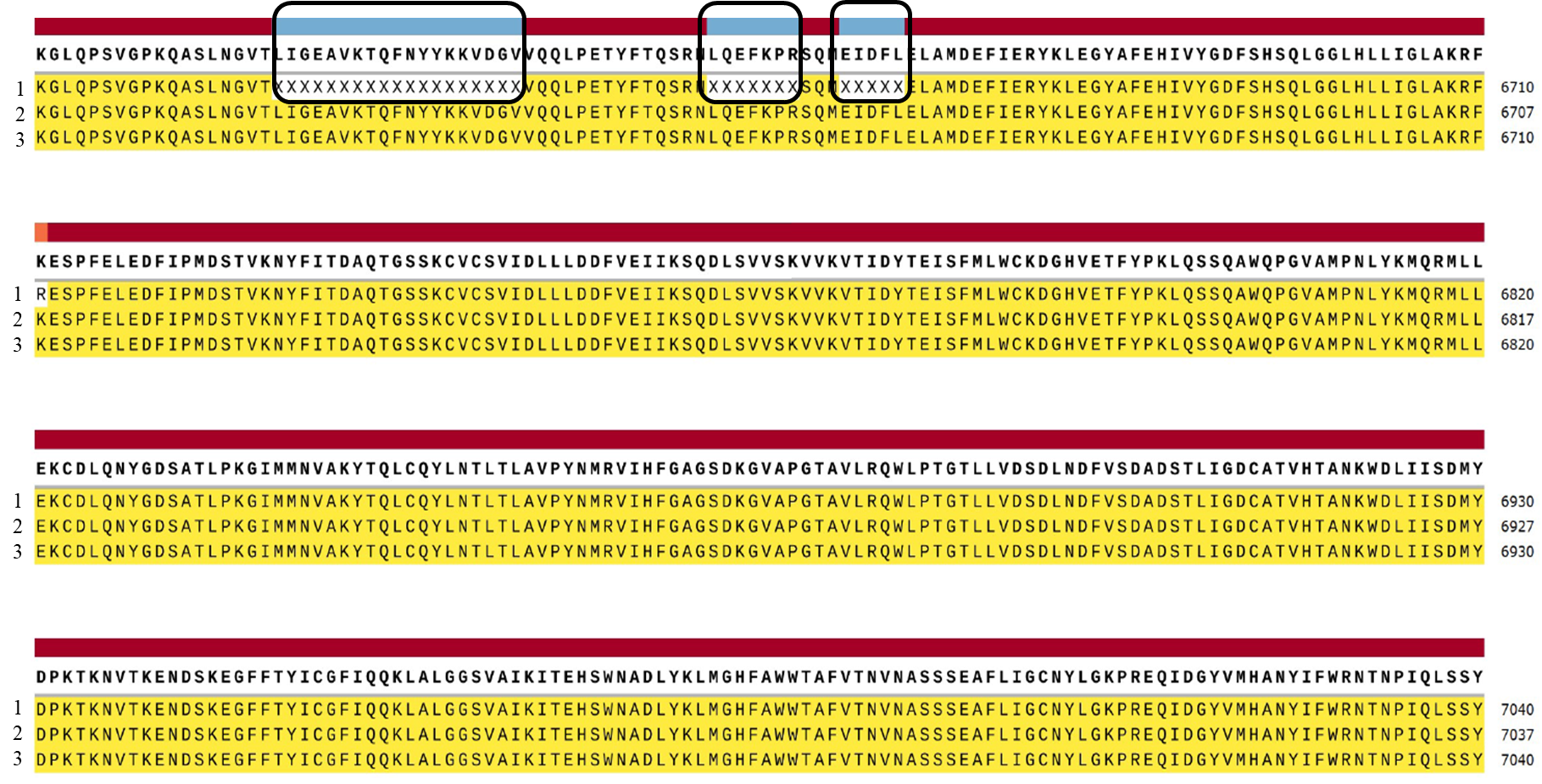

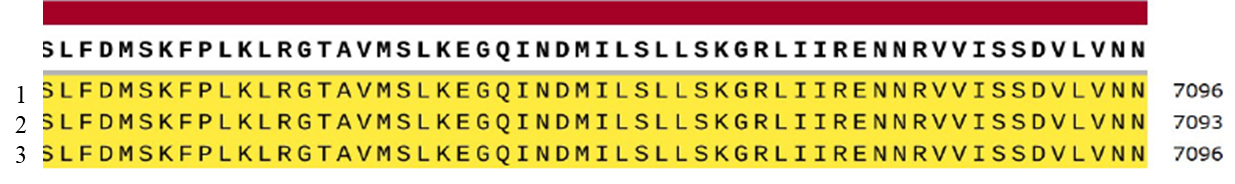

Figure 6: Schematic view of contigs of the ORF1ab polyprotein of 1. Alpha Variant (B.1.1.7) (UDQ41836.1) and 2. Delta Variant (B.1.617.21) (UDU36744.1) of SARs-CoV-2 with 3. reference Strain (Wuhan) (YP_009724389.1)

9. **ORF3a protein**

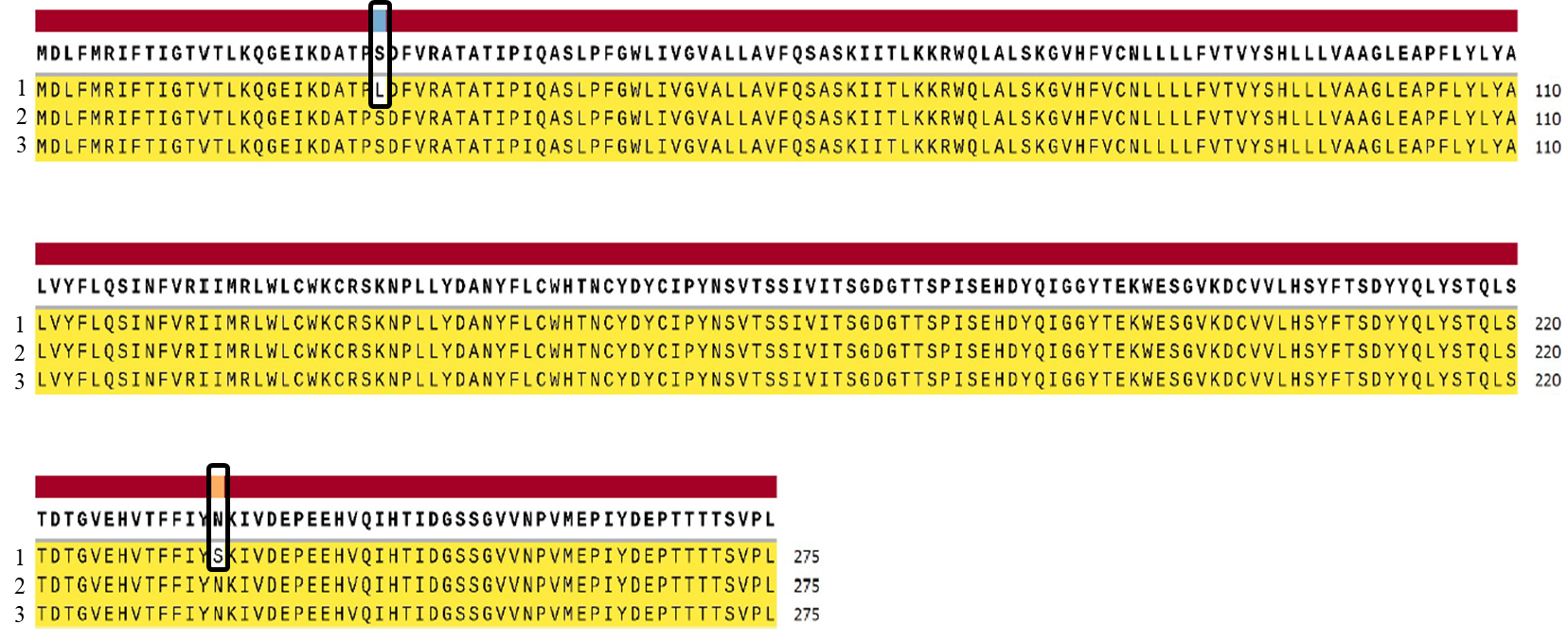

Figure 7: Schematic view of contigs of the ORF3a protein of 1. Alpha Variant (B.1.1.7) (UDQ41839.1) and 2. Delta Variant (B.1.617.21) (UDU36747.1) of SARs-CoV-2 with 3. reference Strain (Wuhan) (YP_009724391.1)

10. **ORF6 protein**

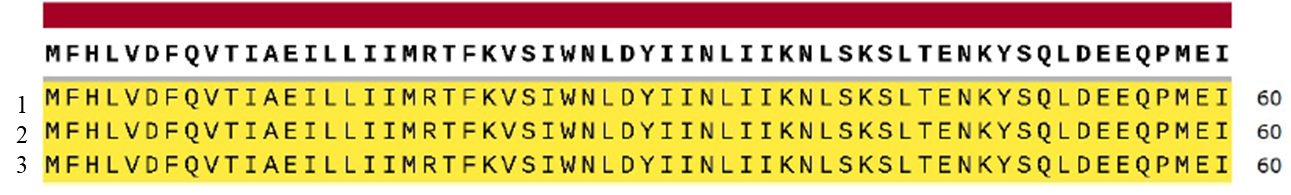

Figure 8: Schematic view of contigs of the ORF6 protein of 1. Alpha Variant (B.1.1.7) (UDQ41842.1) and 2. Delta Variant (B.1.617.21) (UDU36750.1) of SARs-CoV-2 with 3. reference Strain (Wuhan) (YP_009724394.1)

11. **ORF7a protein**

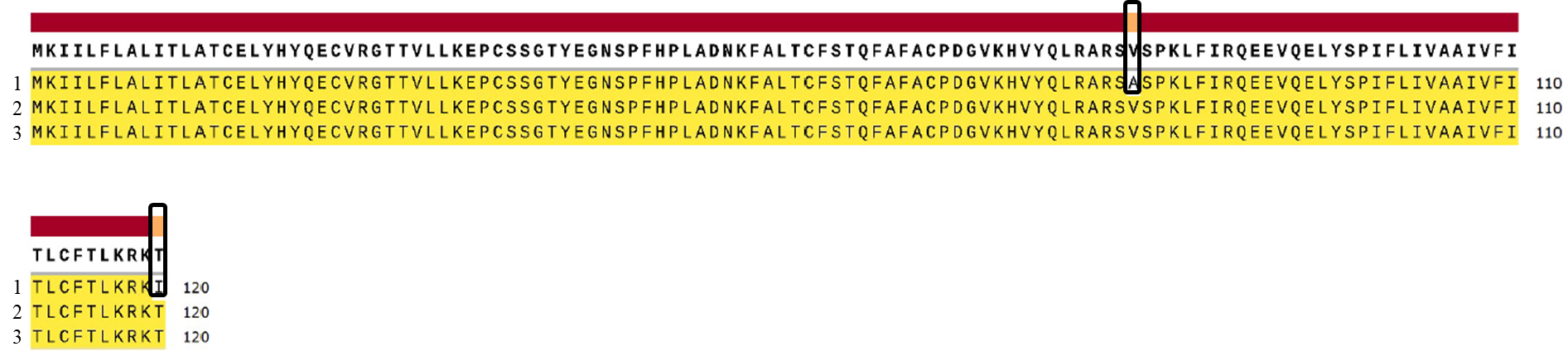

Figure 9: Schematic view of contigs of the ORF7a protein of 1. Alpha Variant (B.1.1.7) (UDQ41843.1) and 2. Delta Variant (B.1.617.21) (UDU36751.1) of SARs-CoV-2 with 3. reference Strain (Wuhan) (YP_009724395.1)

13. **ORF7b protein**

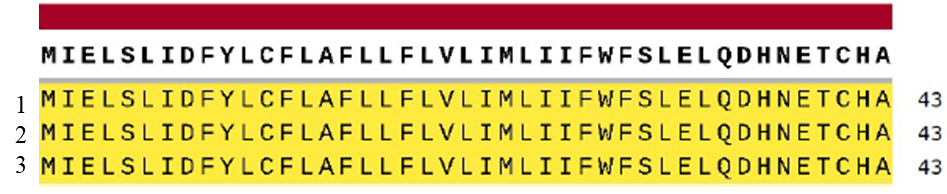

Figure 10: Schematic view of contigs of the ORF7b protein of 1. Alpha Variant (B.1.1.7) (UDQ41844.1) and 2. Delta Variant (B.1.617.21) (UDU36752.1) of SARs-CoV-2 with 3. reference Strain (Wuhan) (YP_009725318.1)

14. **ORF8 protein**

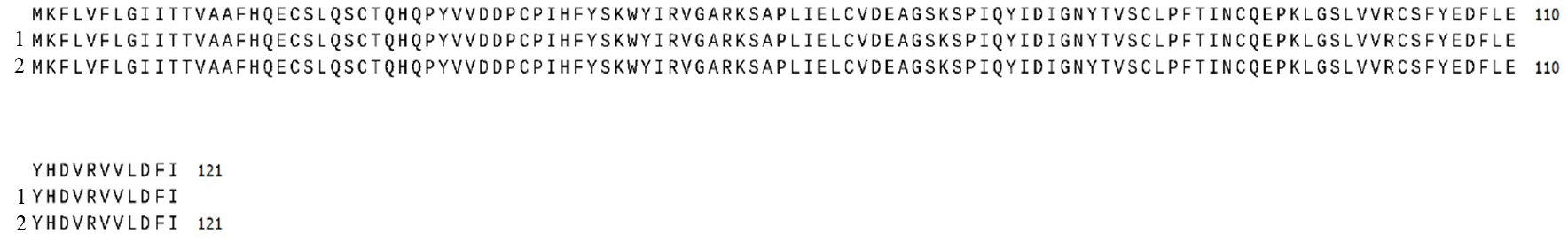

Figure 11: Schematic view of contigs of the ORF8 protein of 1. Delta Variant (B.1.617.21) (UDU36753.1) of SARs-CoV-2 with 2. reference Strain (Wuhan) (YP_009724396.1)

**Membrane Protein**

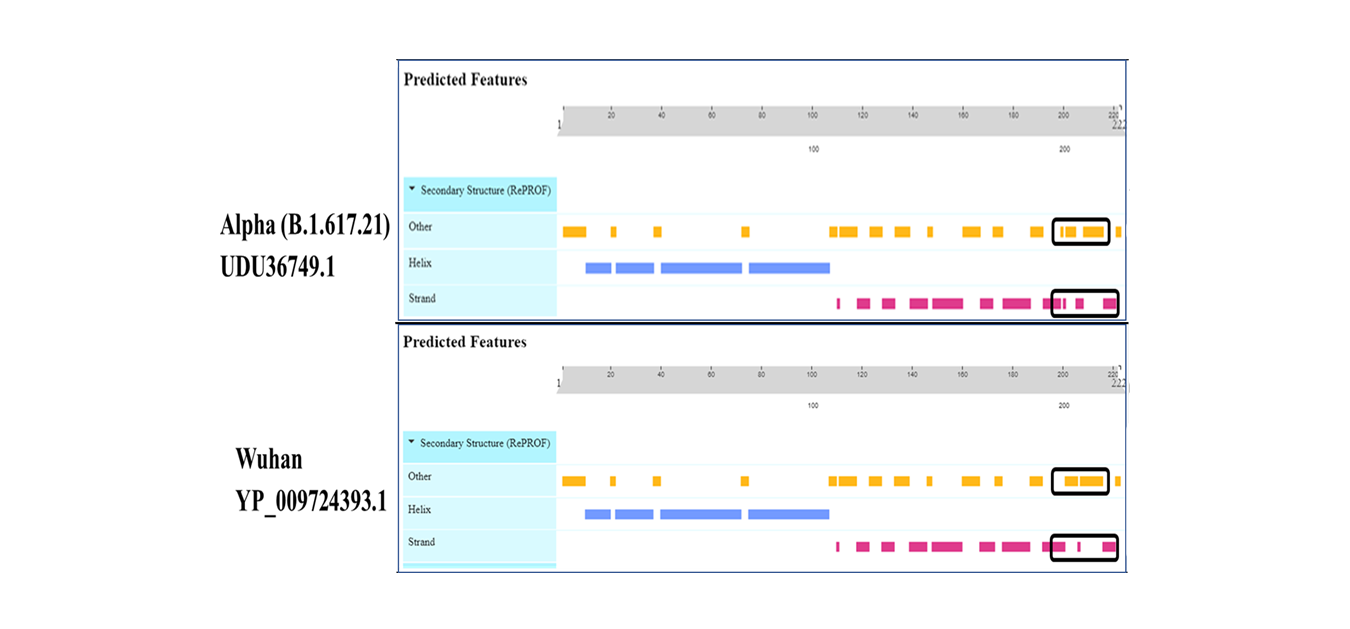

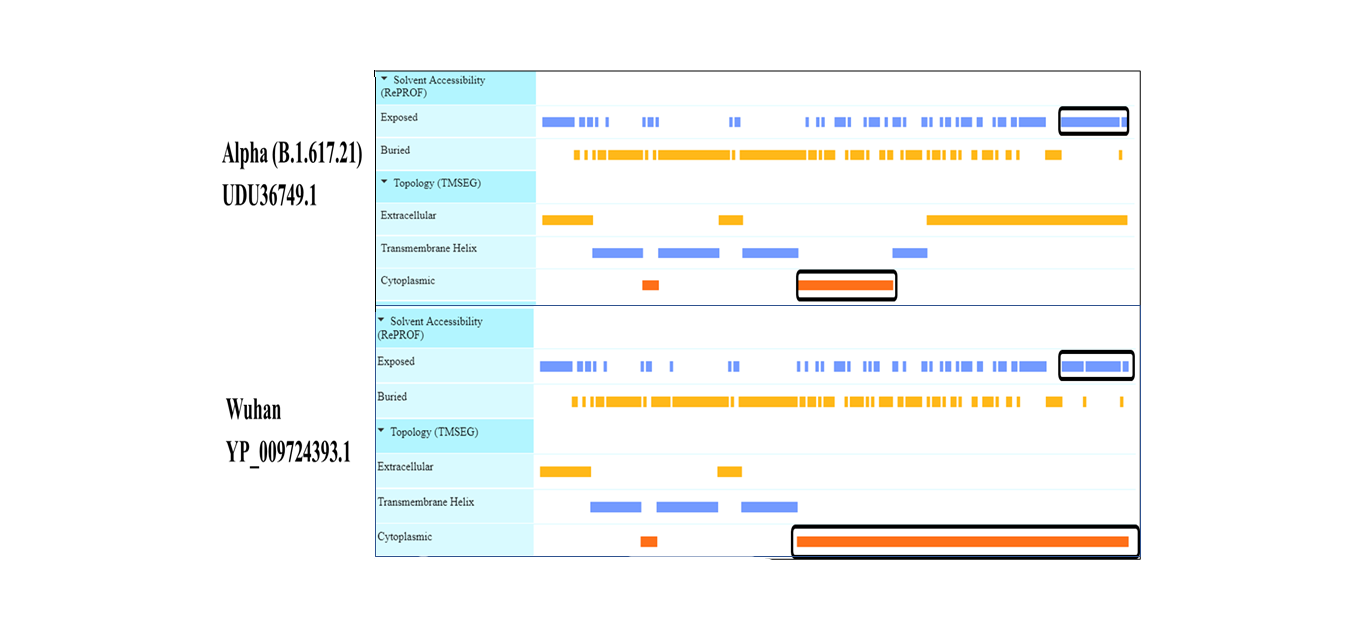

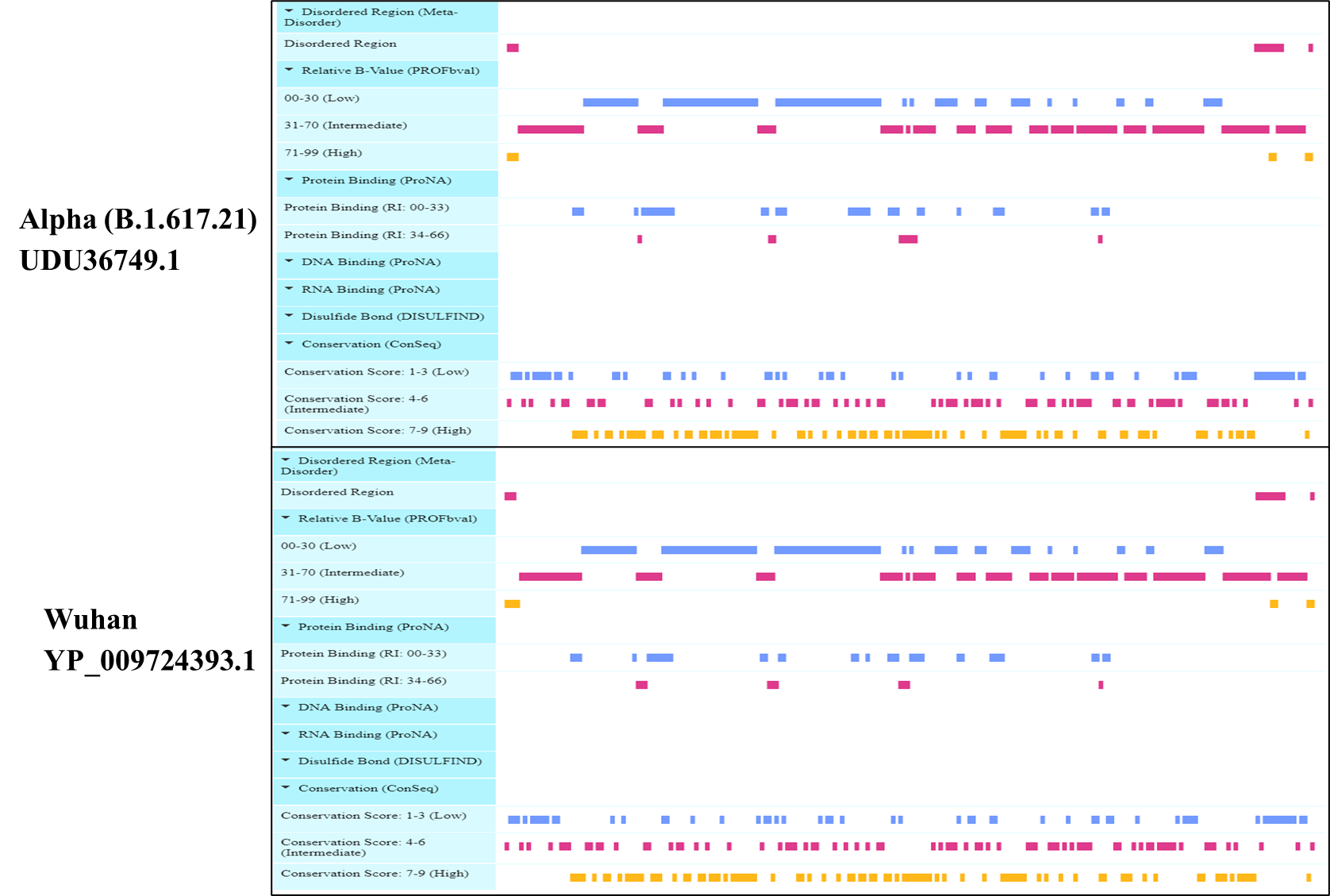

Figure 12 Viewer lays out predicted features of protein structural and functional features of Membrane Glycoproteins.

**Nucleocapsid phosphoprotein**

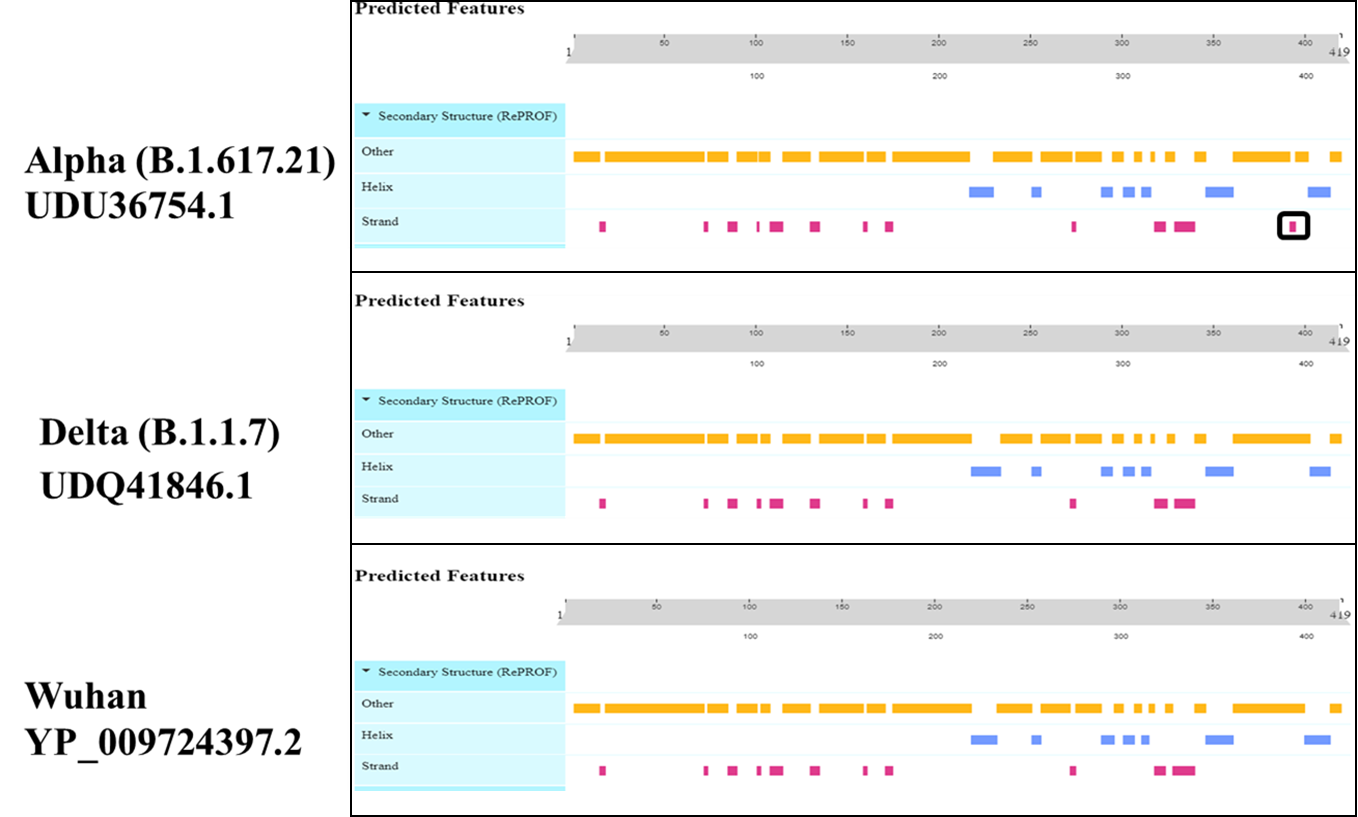

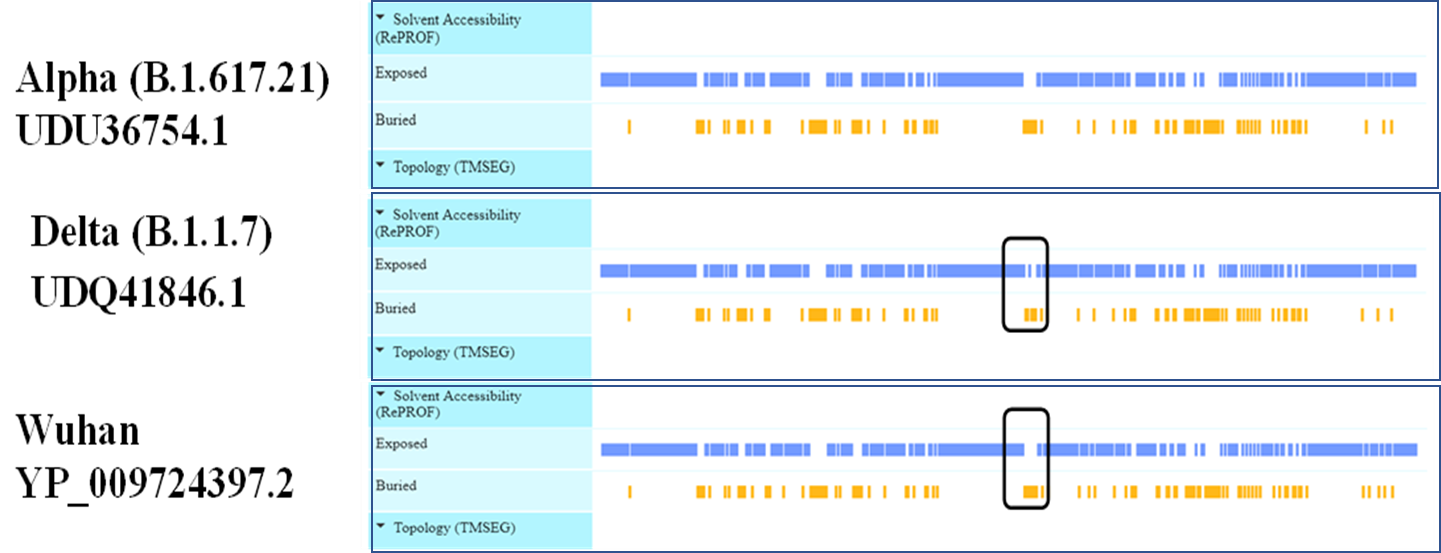

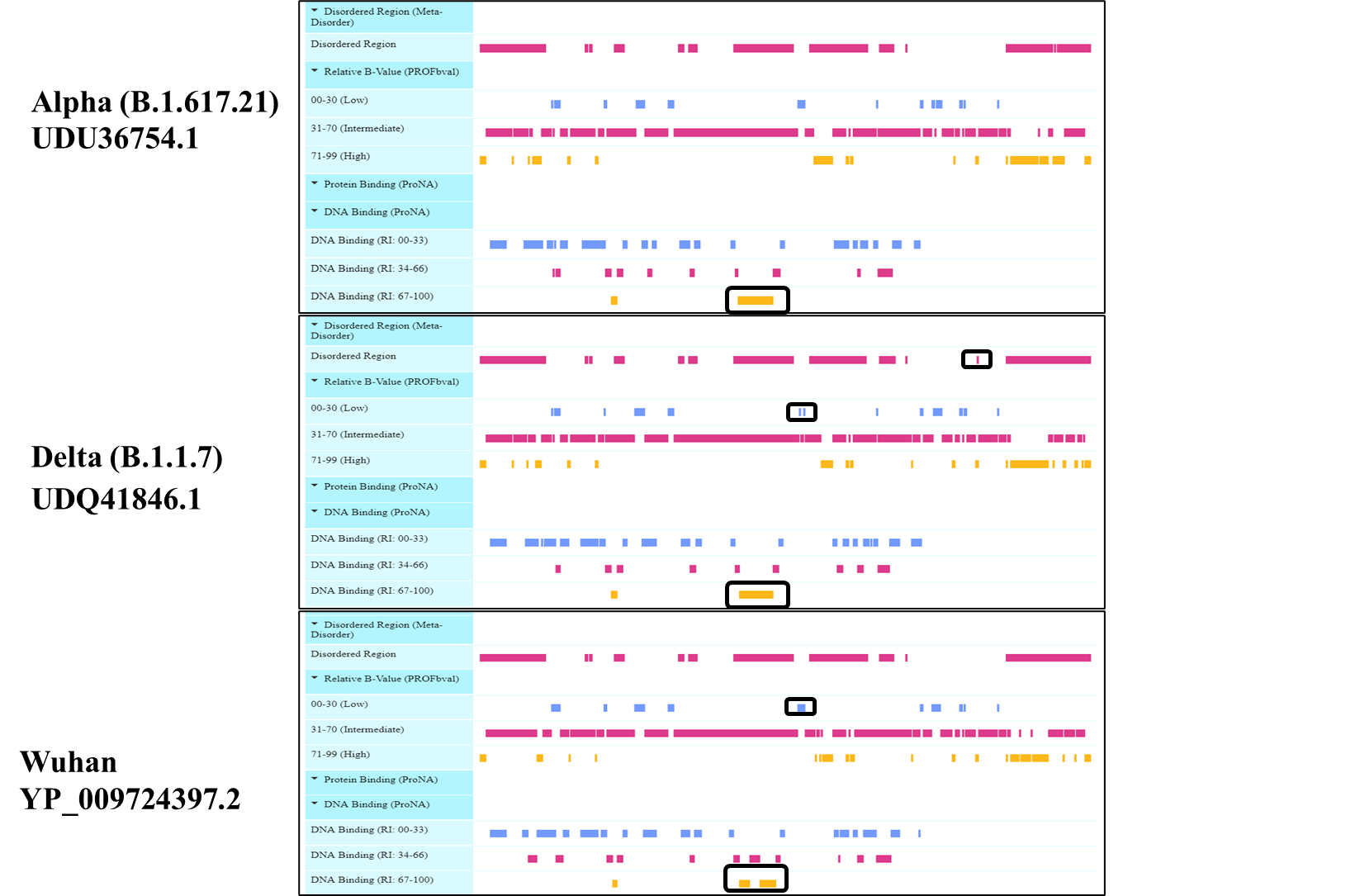

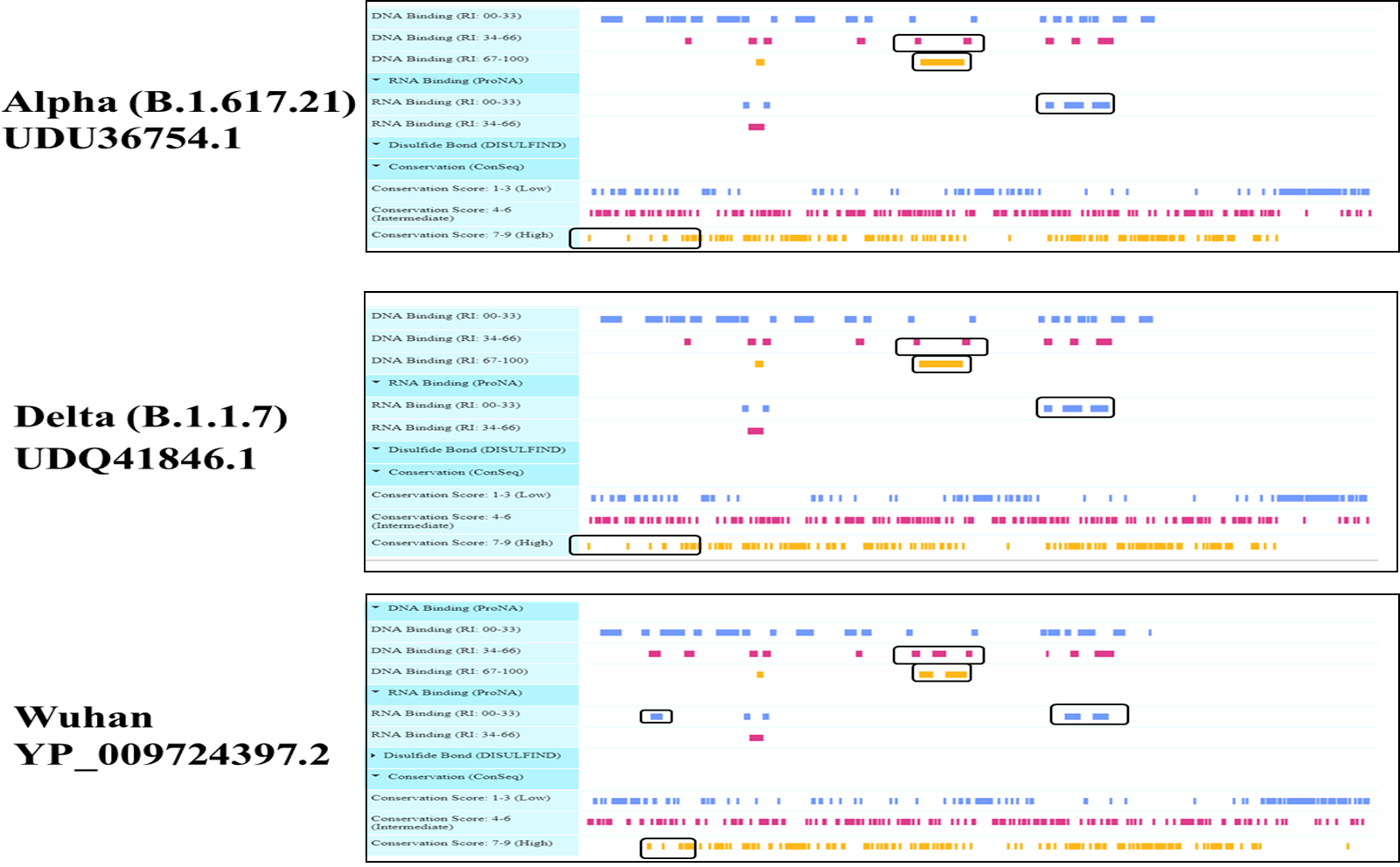

Figure 13 Viewer lays out predicted features of protein structural and functional features of Nucleocapsid phosphoprotein

2. **ORF1a Polyprotein**

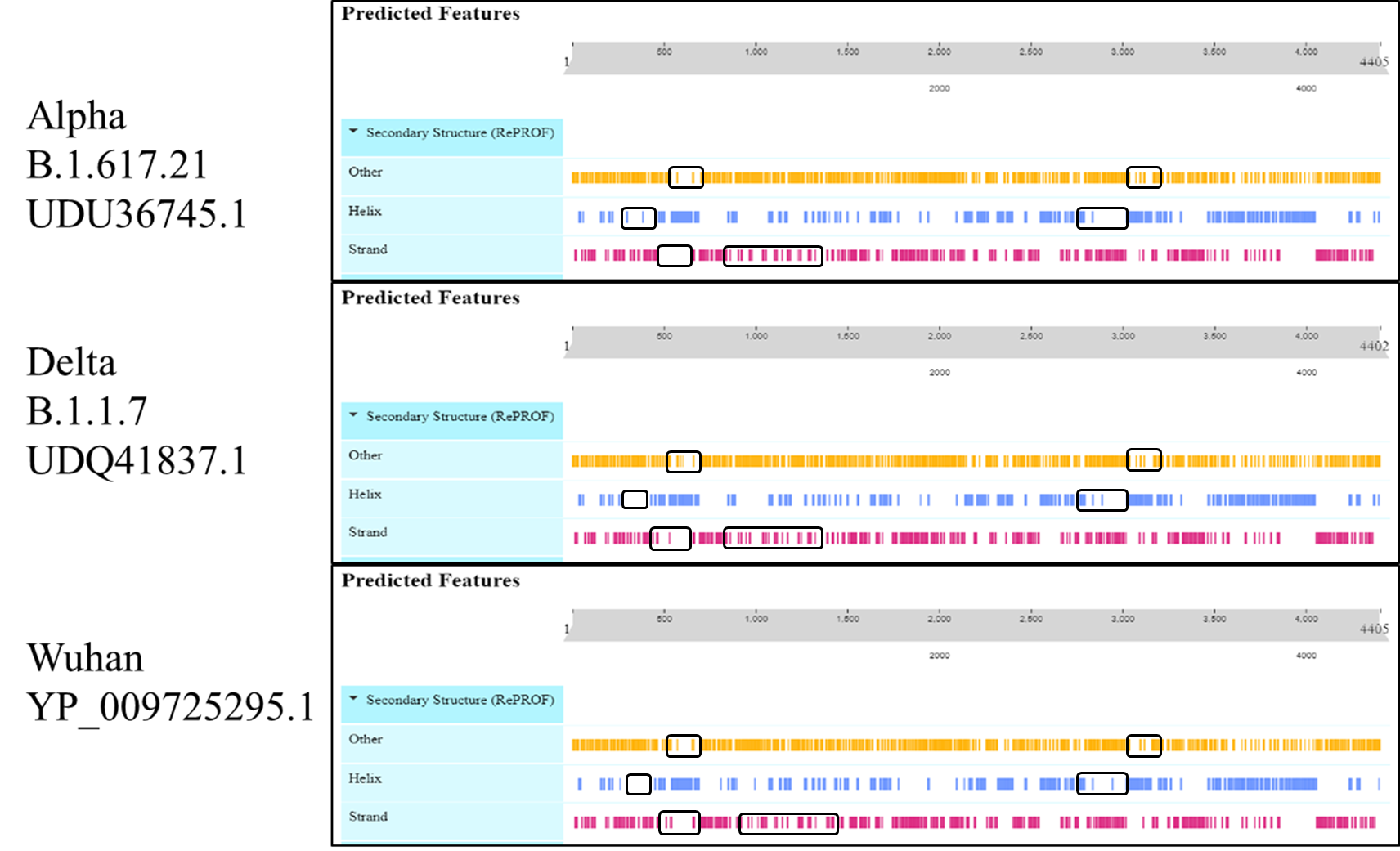

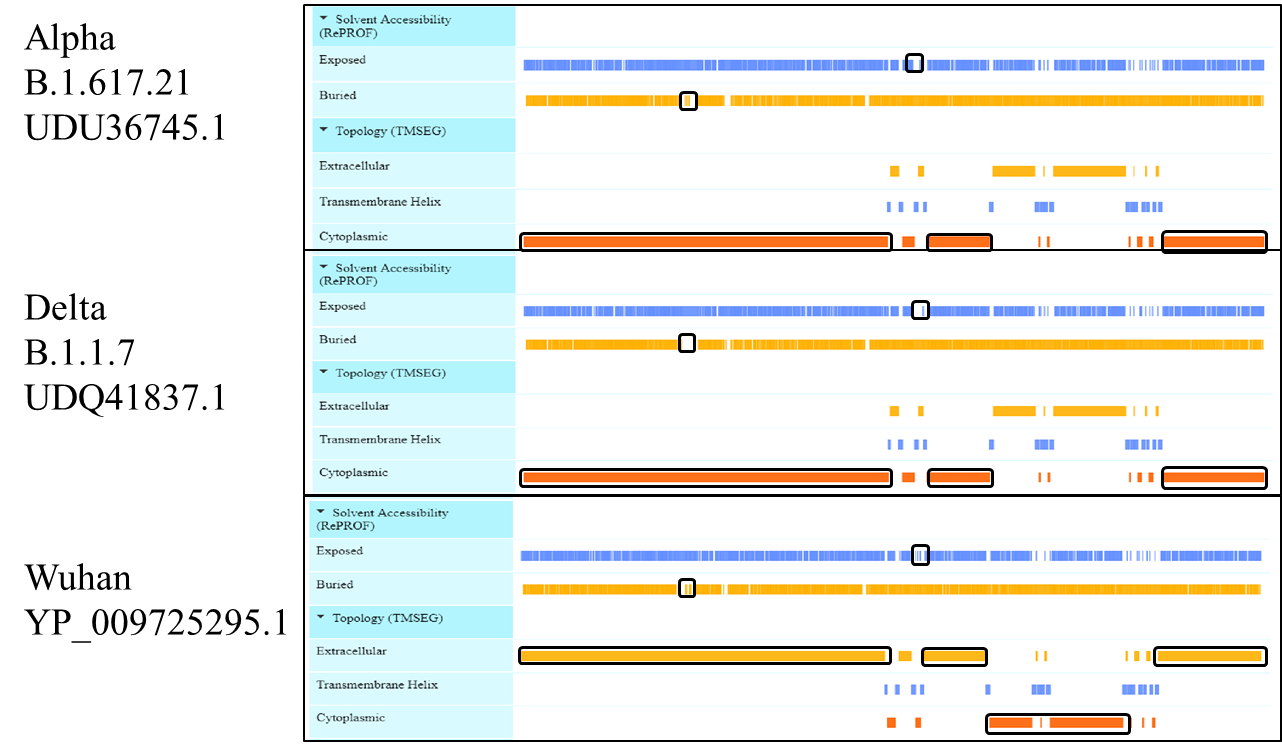

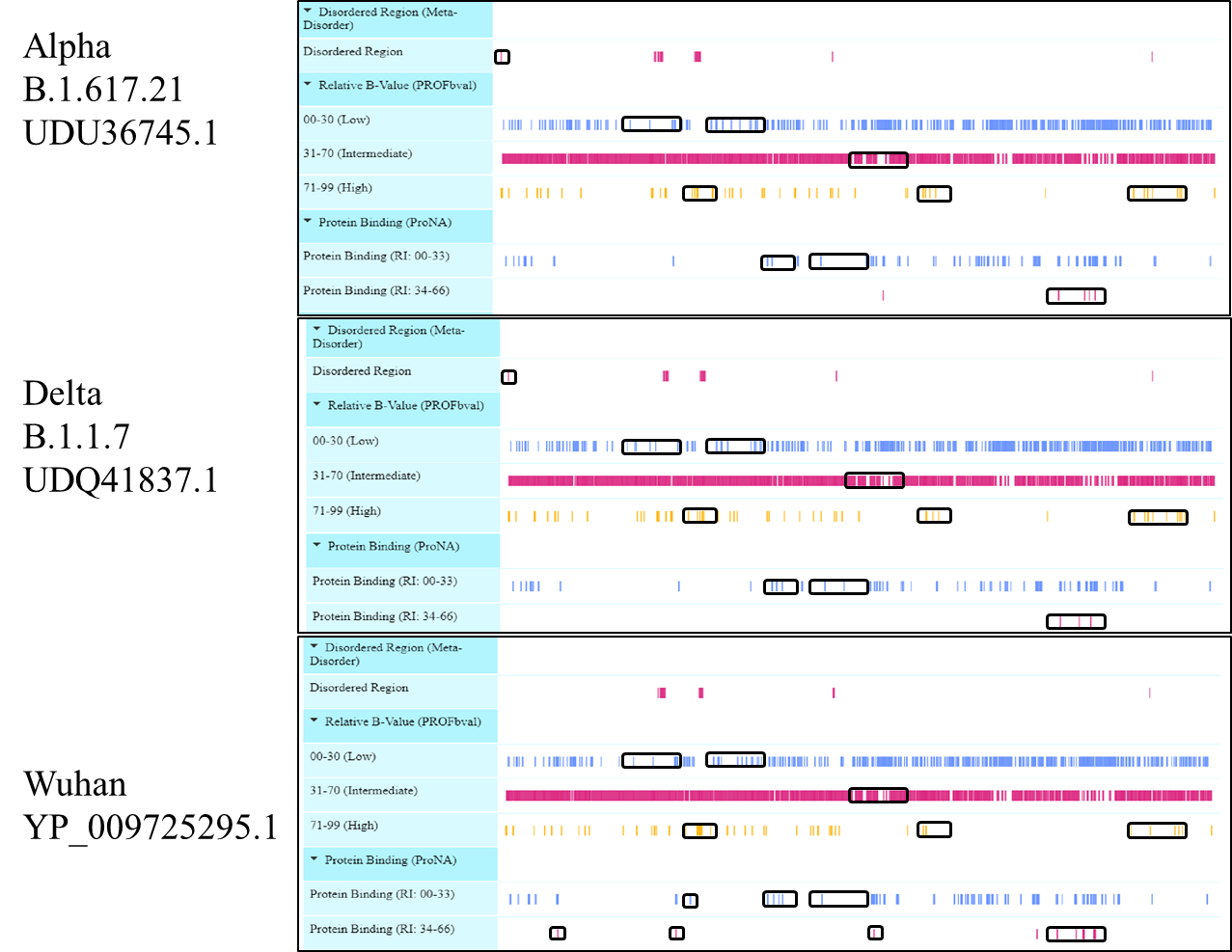

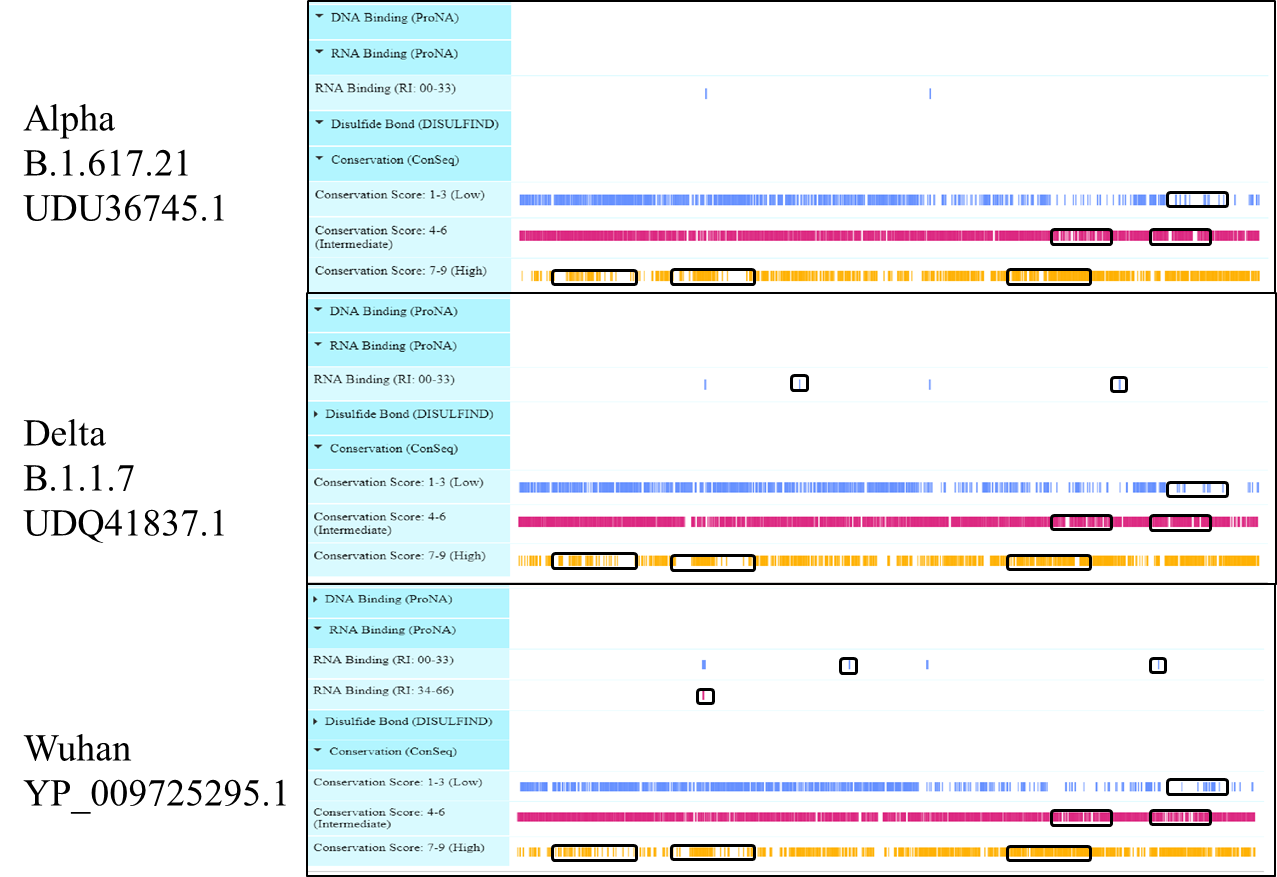

Figure 14 Viewer lays out predicted features of protein structural and functional features of ORF1a Polyprotein

**ORF1ab Polyprotein**

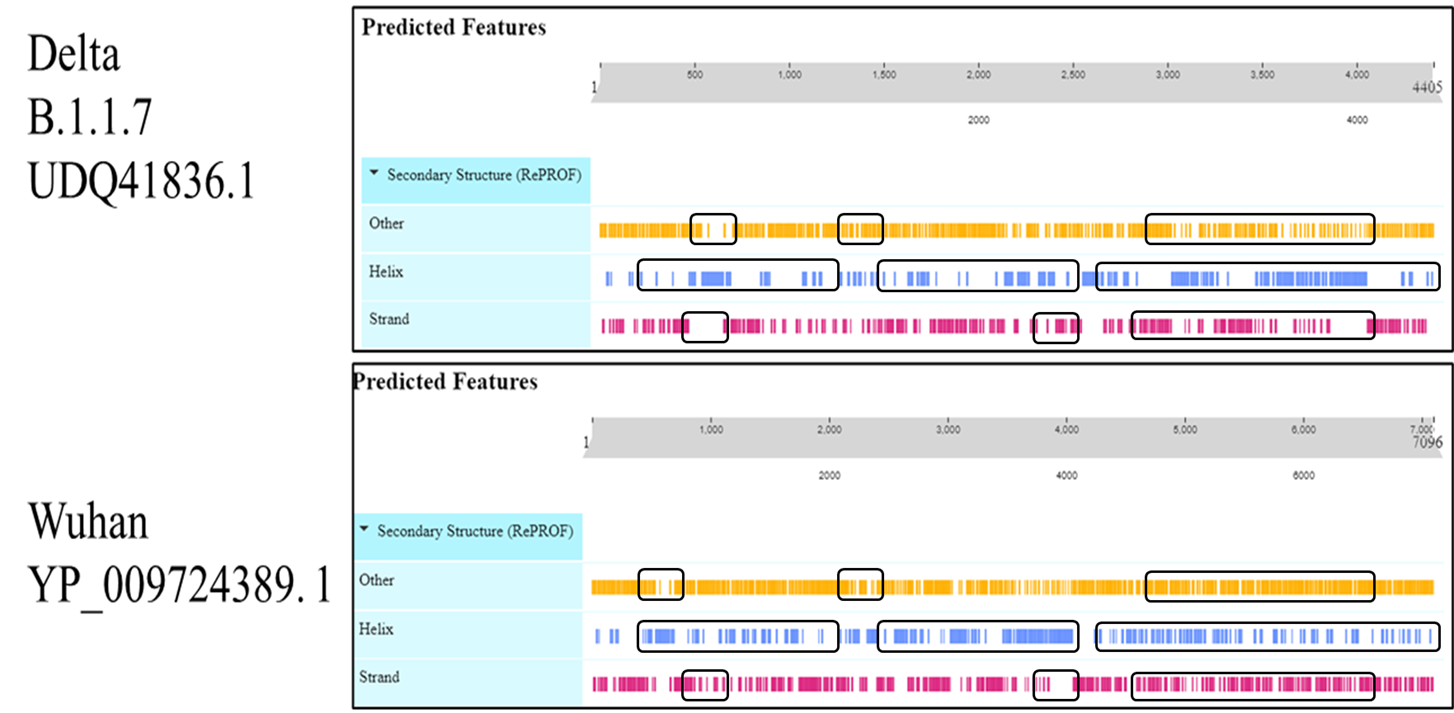

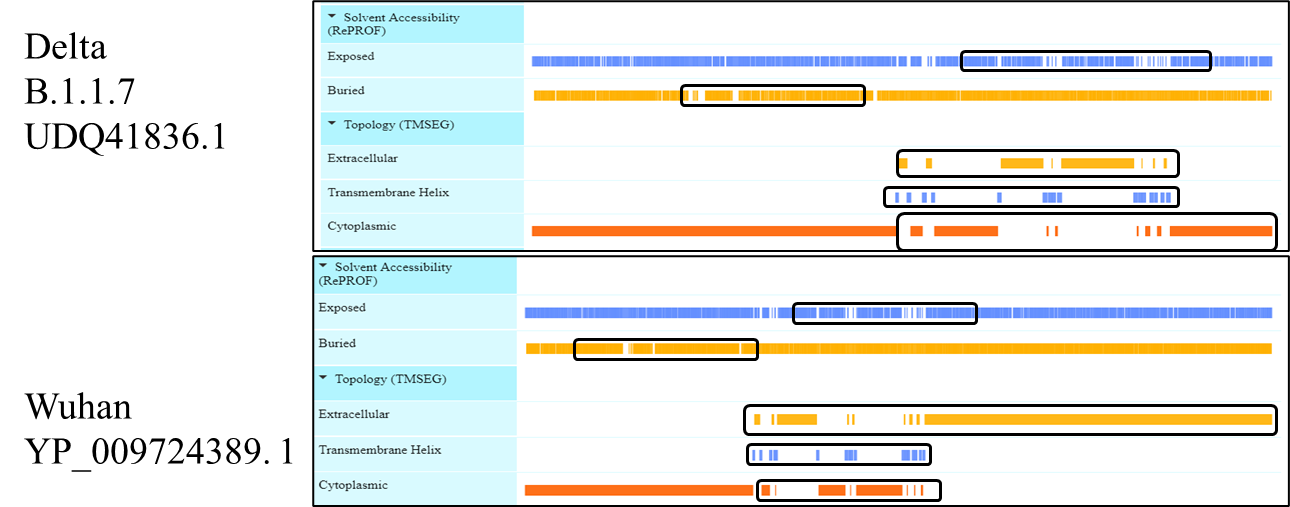

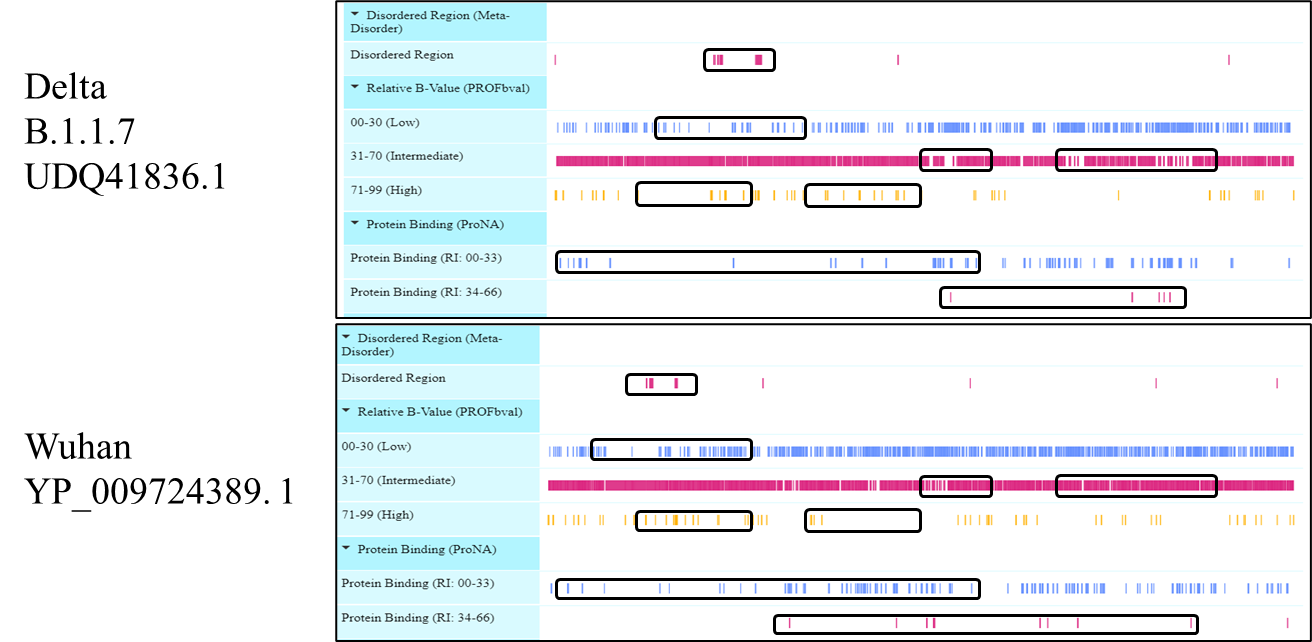

Figure 15 Viewer lays out predicted features of protein structural and functional features of ORF1ab Polyprotein

3. **ORF7a protein**

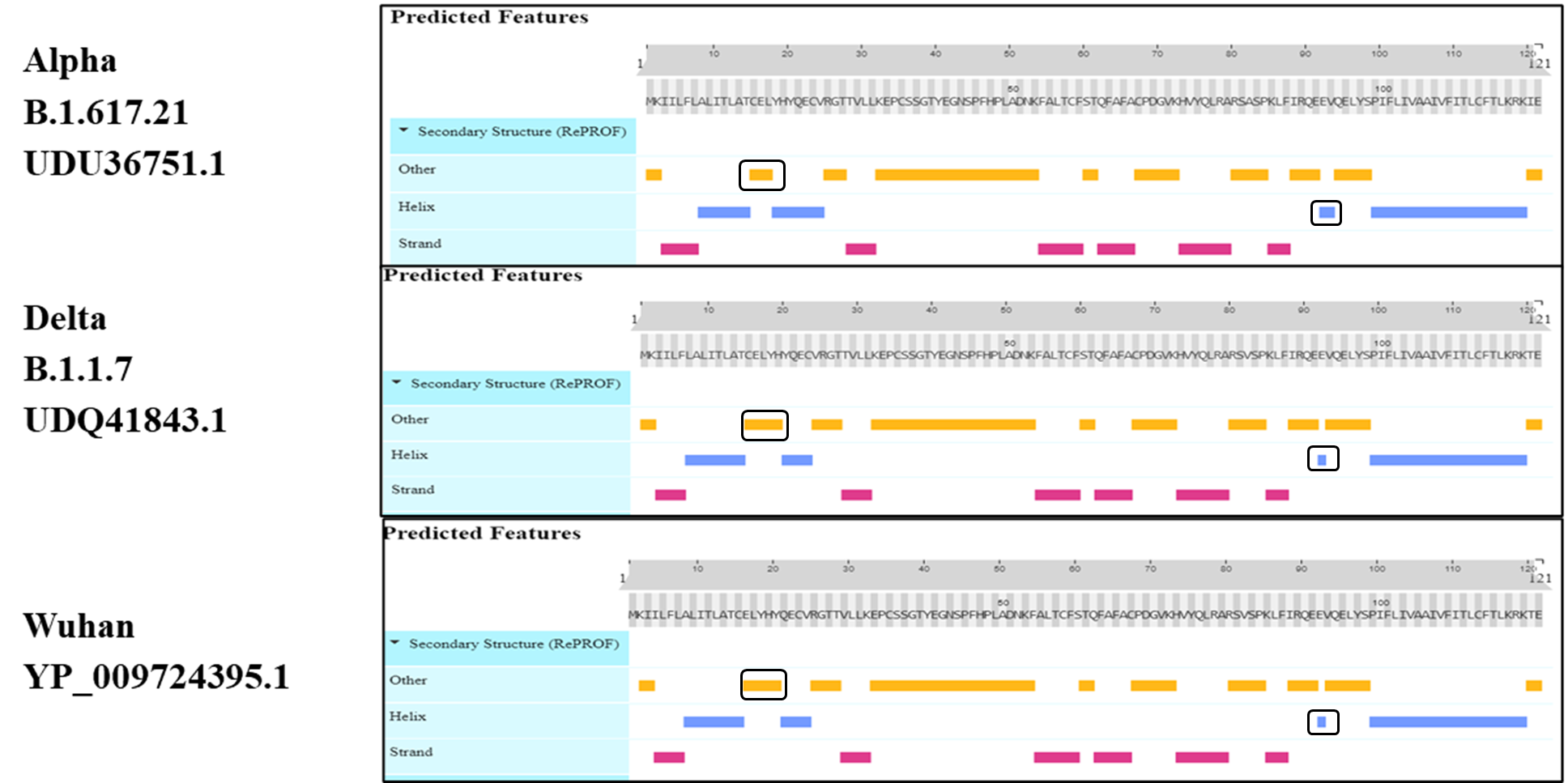

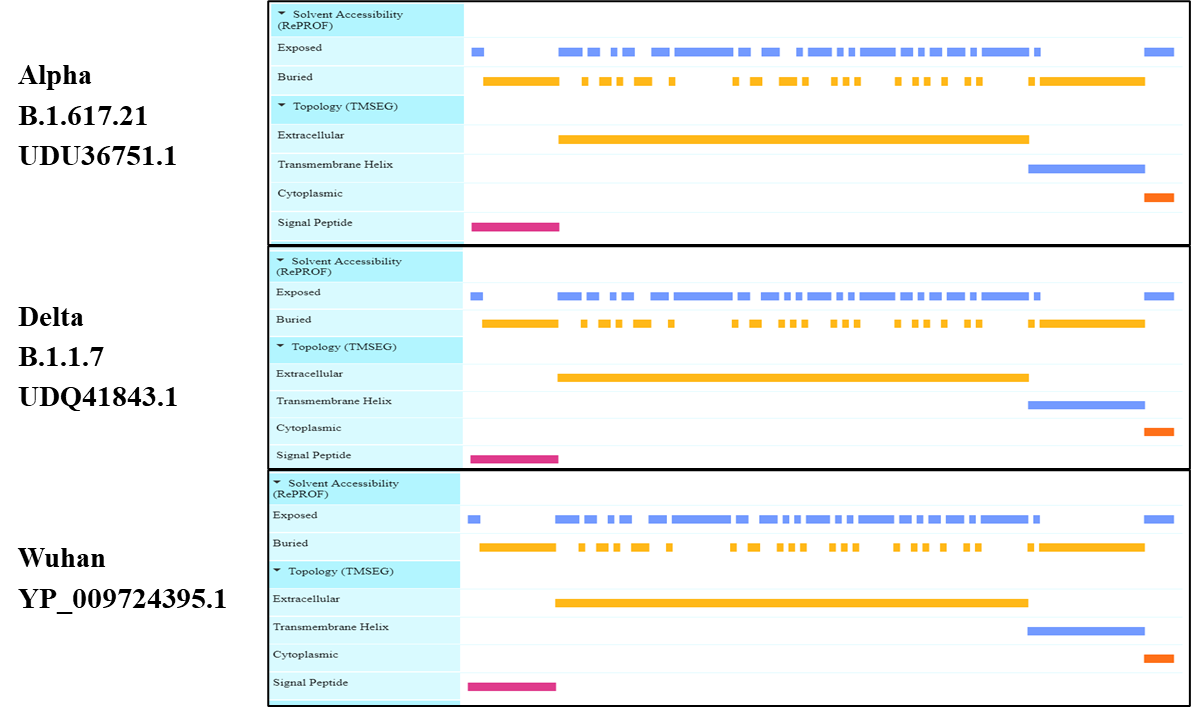

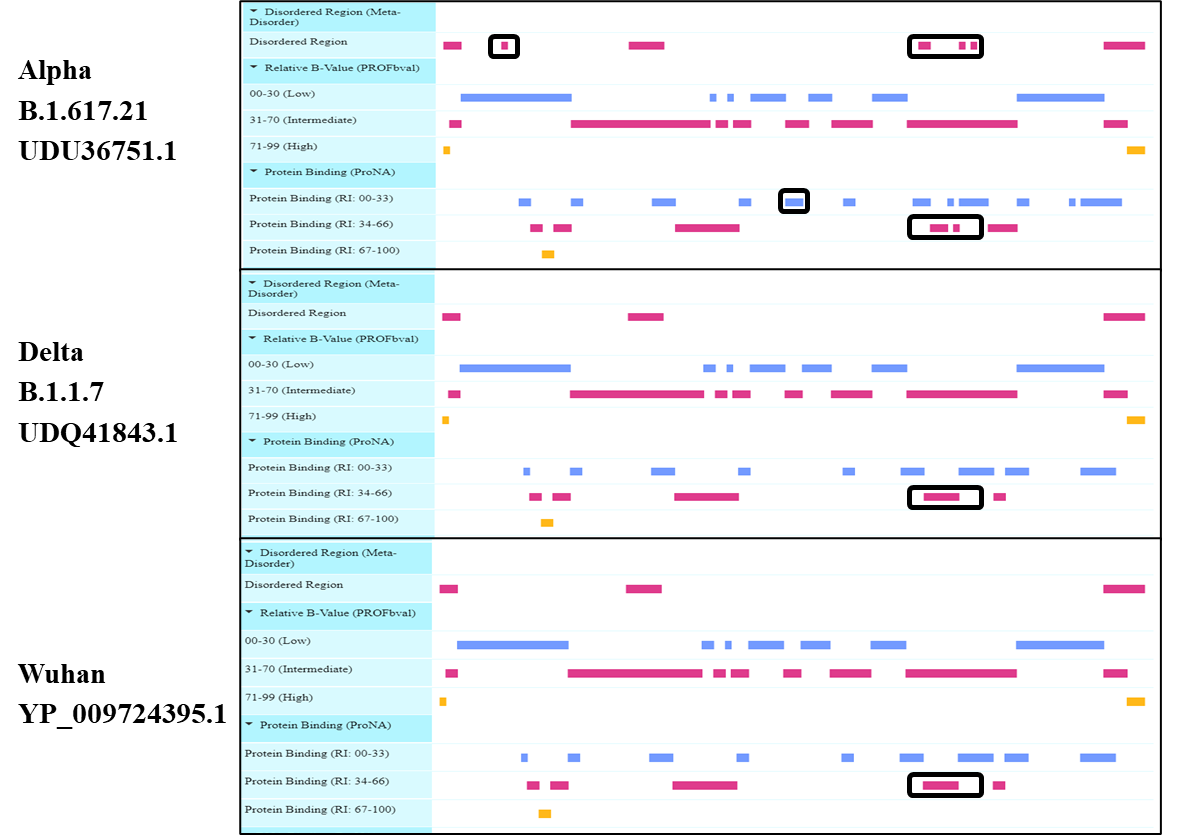

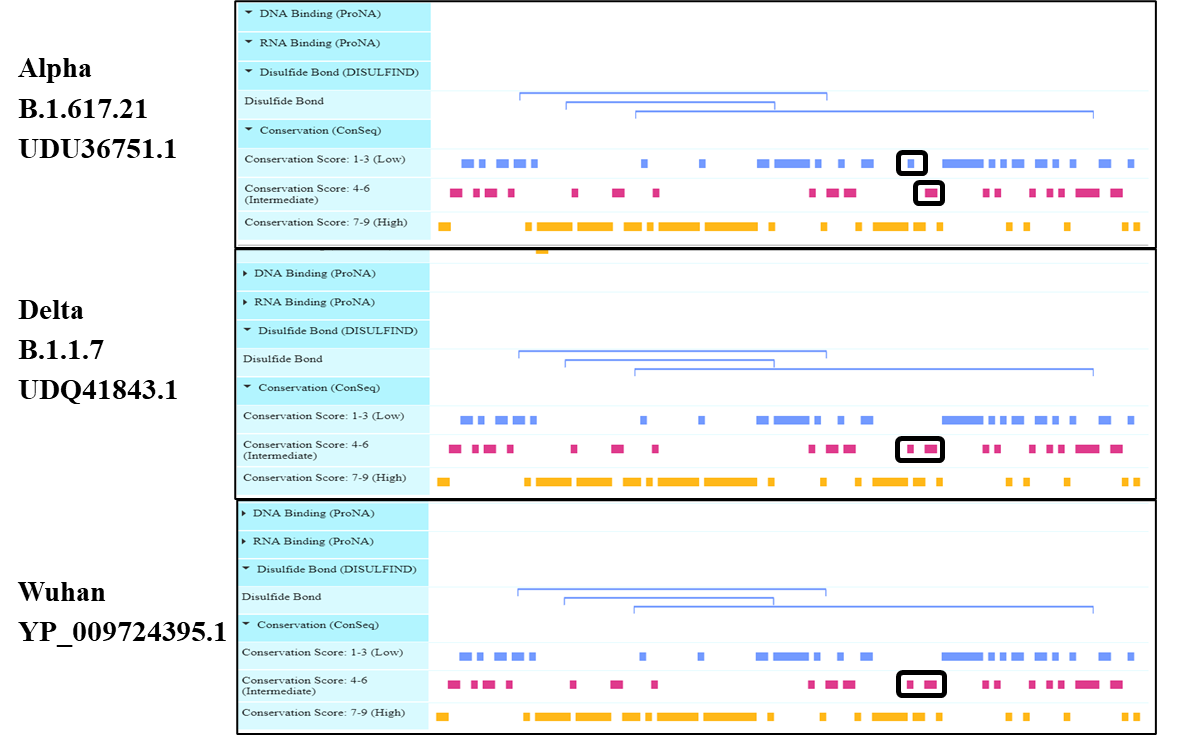

Figure 16 Viewer lays out predicted features of protein structural and functional features of ORF7a protein

**Supplementary Tables**

| **Alpha** | | **Delta** | |
| --- | --- | --- | --- |
| **Mutation** | **Count** | **Mutation** | **Count** |
| **Envelope** | | | |
|  |  | E_V62F | 1 |
| **Membrane** | | | |
|  |  | M_I82T | 8 |
| **Nucleocapsid phosphoprotein** | | | |
| N_D3L | 5 | N_A90S | 1 |
| N_G204R | 5 | N_D377Y | 8 |
| N_Q389H | 1 | N_D63G | 8 |
| N_R203K | 5 | N_G215C | 4 |
| N_S235F | 5 | N_H300Y | 1 |
|  |  | N_R203M | 8 |
|  |  | N_R385K | 1 |
| **NS3** | | | |
| NS3_K16N | 1 | NS3_A23V | 1 |
|  |  | NS3_G49V | 1 |
|  |  | NS3_I118T | 1 |
|  |  | NS3_I62F | 1 |
|  |  | NS3_K16T | 1 |
|  |  | NS3_L65I | 1 |
|  |  | NS3_L73I | 1 |
|  |  | NS3_Q116H | 1 |
|  |  | NS3_S26L | 8 |
|  |  | NS3_T221K | 1 |
|  |  | NS3_Y211H | 1 |
| **NS7a** | | | |
|  |  | NS7a_L116F | 2 |
|  |  | NS7a_T120I | 8 |
|  |  | NS7a_V82A | 8 |
| **NS7b** | | | |
|  |  | NS7b_T40I | 4 |
| **NS8** | | | |
| NS8_R52I | 5 | NS8_Q27stop | 5 |
| NS8_V62L | 1 |  |  |
| NS8_Y73C | 5 |  | |
| **NSP12** | | | |
| NSP12_P323L | 5 | NSP12_G228S | |
|  |  | NSP12_G671S | |
|  |  | NSP12_P323L | |
|  |  | NSP12_Q357H | |
|  |  | NSP12_V111L | |
| **NSP13** | | | |
|  |  | NSP13_M576I | |
|  |  | NSP13_P77L | |
|  |  | NSP13_R392C | |
|  |  | NSP13_S350L | |
| **NSP14** | | | |
|  |  | NSP14_A394V | |
|  |  | NSP14_D144Y | |
|  |  | NSP14_M72I | |
|  |  | NSP14_P46L | |
|  |  | NSP14_T113I | |
| **NSP15** | | | |
|  |  | NSP15_G229C | |
|  |  | NSP15_H234Y | |
|  |  | NSP15_V66L | |
| **NSP16** | | | |
|  |  | NSP16_K160R | |
|  |  | NSP16_M270I | |
| **NSP2** | | | |
| NSP2_E345K | 1 | NSP2_A386S | |
| NSP2_L550F | 1 | NSP2_P129L | |
|  |  | NSP2_Y16H | |
| **NSP3** | | | |
| NSP3_A1321V | 1 | NSP3_A416V | 1 |
| NSP3_A1819V | 1 | NSP3_A488S | 4 |
| NSP3_A1941V | 1 | NSP3_H1274Y | 1 |
| NSP3_A890D | 5 | NSP3_K1693N | 1 |
| NSP3_I1412T | 5 | NSP3_P1228L | 4 |
| NSP3_P153L | 1 | NSP3_P1469S | 4 |
| NSP3_R586C | 1 | NSP3_P822L | 4 |
| NSP3_T183I | 5 | NSP3_S1285F | 1 |
| NSP3_T423I | 1 | NSP3_S1370F | 1 |
| NSP3_T779I | 1 | NSP3_S1424F | 1 |
|  |  | NSP3_V245F | |
| **NSP4** | | | |
|  |  | NSP4_A446V | 4 |
|  |  | NSP4_T492I | 4 |
|  |  | NSP4_V167L | 4 |
| **NSP5** | | | |
|  |  | NSP5_V86L | 1 |
| **NSP6** | | | |
| NSP6_F108del | 5 | NSP6_T181I | 2 |
| NSP6_G107del | 5 | NSP6_T77A | 4 |
| NSP6_S106del | 5 | NSP6_V149A | 4 |
| **Spike** | | | |
| Spike_A570D | 5 | Spike_A1078V | 1 |
| Spike_A67S | 1 | Spike_A222V | 2 |
| Spike_A688V | 1 | Spike_C1250W | 1 |
| Spike_D1118H | 5 | Spike_D138Y | 1 |
| Spike_D614G | 5 | Spike_D215H | 1 |
| Spike_H69del | 5 | Spike_D574Y | 1 |
| Spike_N501Y | 5 | Spike_D614G | 8 |
| Spike_P681H | 5 | Spike_D950N | 6 |
| Spike_S982A | 5 | Spike_E156G | 8 |
| Spike_S98F | 1 | Spike_E484Q | 1 |
| Spike_T716I | 5 | Spike_F157del | 8 |
| Spike_V70del | 5 | Spike_G142D | 6 |
| Spike_S98F | 1 | Spike_I850L | 1 |
| Spike_Y144del | 5 | Spike_L1141W | 1 |
|  |  | Spike_L452R | 8 |
|  |  | Spike_P681R | 8 |
|  |  | Spike_Q613H | 1 |
|  |  | Spike_R158del | 8 |
|  |  | Spike_T19R | 8 |
|  |  | Spike_T478K | 8 |
|  |  | Spike_T95I | 4 |
|  |  | Spike_V483A | 1 |

Table 2: Mutation of amino acids in Alpha & Delta Variant of SARs-CoV-2 from Pakistan with reference strain (hCoV-19/Wuhan/WIV04/2019).
